# Spatial organization of a SPINK1-associated tumor-stromal ecosystem links metabolic remodeling to immune restriction in bladder cancer

**DOI:** 10.64898/2026.11.18.767479

**Authors:** Qiyi Yu, Yifan Wu, Xuedan Han, Yu Lu, Huilong Li, Yi Da Douglas Jiang, Jingyi Zhou, Yan Li, Hang Huang, Wei Chen, Xiaoman Li, Fangdie Ye, Lufeng Zheng

**Author notes:** These authors contributed equally to this work (Qiyi Yu, Yifan Wu, Xuedan Han). Corresponding authors: Lufeng Zheng; Fangdie Ye; Xiaoman Li.

## Abstract

Spatially organized tumor–stromal interactions contribute to metabolic remodeling and immune suppression in bladder cancer (BLCA). Using spatial transcriptional programs to define single-cell states, we identified a niche comprising SPINK1-high epithelial cells (E2) and TPM2-high cancer-associated fibroblasts (CAFs) (C1). SPINK1 promoted tumor growth and myofibroblastic CAF programs, with EGFR and TGF-β signaling implicated in tumor–CAF communication. Inferred metabolic reactions and spatial metabolite distributions supported a candidate circuit linking tumor-derived lactate to CAF oxidative metabolism and glutamine provision, followed by tumor glutamine utilization for reductive carboxylation and fatty acid synthesis. SPINK1 perturbation altered extracellular glutamine and lactate concentrations. Predicted chemokine interactions implicated C2/C3 CAFs in T cell recruitment, whereas the E2– C1 niche exhibited checkpoint and exhaustion-associated features. RHCG-positive tumor regions contained higher proportions of TOX⁺CXCR4⁺ cells among CD3⁺ T cells, suggesting a possible association between ammonia handling and T cell dysfunction. *In vivo*, *Spink1* loss attenuated tumor progression, increased cytotoxic CD8⁺ T-cell infiltration, and enhanced tumor control during PD-1 blockade. These findings identify a spatially organized tumor–stromal niche associated with metabolic cooperation and immune restriction, supporting SPINK1 as a candidate therapeutic target in combination with immunotherapy.

## Introduction

Bladder cancer (BLCA) is a common urinary tract malignancy with substantial morbidity and mortality worldwide [1]. Approximately 75% of patients present with non-muscle-invasive disease, which frequently recurs and can progress to muscle-invasive cancer [2]. Disease progression and treatment response reflect interactions between malignant cells and the surrounding microenvironment. Examining these interactions within intact tissues can reveal relationships between cellular states that are obscured by analyses of malignant cells alone.

Metabolic profiling has identified alterations in amino acid, lipid, nucleotide, and energy metabolism in BLCA [3, 4], although bulk measurements do not resolve their cellular sources or spatial distribution. Stromal nutrient provision supports tumor metabolism in other malignancies, exemplified by alanine secretion from pancreatic stellate cells [5], whereas competition for glucose can impair T cell effector function [6]. Local metabolite accumulation can impose additional constraints on immunity; extracellular ammonia, for example, promotes T cell exhaustion in colorectal cancer [7]. These observations indicate that metabolic conditions supporting tumor growth can also influence neighboring immune cells, motivating the joint investigation of nutrient provision and immune function within defined tissue regions.

Single-cell studies have identified diverse epithelial and cancer-associated fibroblast (CAF) states in BLCA, including CAF populations associated with adverse clinical features [8]. In metastatic urothelial cancer, fibroblast TGF-β response signatures were associated with T cell exclusion and resistance to PD-L1 blockade [9]. More recent work implicated IFN-γ-responsive CAFs and CXCL12 signaling in the recruitment of CXCR4-expressing T cells in muscle-invasive BLCA [10]. Chemokine-mediated recruitment brings T cells into environments whose nutrient availability, metabolite composition, and regulatory signals may constrain their effector function. Molecular annotations and overall cell fractions do not fully capture the spatial relationships through which particular tumor and CAF states may shape these conditions. Resolving these relationships is necessary to assess whether local metabolic support for tumor growth is accompanied by impaired antitumor immunity.

Spatial transcriptomics and mass spectrometry imaging provide complementary measurements of gene expression and metabolite-associated signals across tissue regions [11]. Here, we used spatial transcriptional programs to guide single-cell state definition and integrated spatial metabolomics, multiplex imaging, and functional perturbations to characterize tumor–stromal interactions in BLCA. We identified a niche comprising SPINK1-high epithelial cells and TPM2-high CAFs. Functional analyses implicated SPINK1 in tumor growth and myofibroblastic CAF programs, while spatial analyses linked this niche to potentially complementary metabolic programs and exhaustion-associated T cell phenotypes. Genetic *Spink1* loss suppressed tumor progression and increased CD8⁺ T cell infiltration, with improved tumor control in combination with PD-1 blockade. These findings support further evaluation of SPINK1 inhibition alongside immunotherapy.

## Results

### A multimodal atlas reveals cellular and metabolic organization in human BLCA

To establish a comprehensive molecular atlas of BLCA, we assembled a multi-omics cohort spanning diverse clinical and pathological contexts (**Figure 1A; Figure S1A**). We integrated single-cell RNA sequencing (scRNA-seq) data from 82 samples, comprising 66 tumors and 16 adjacent tissues, with a total of 528,112 cells. The cohort encompassed substantial clinical heterogeneity, including tumors of different grades and TNM stages, primary and recurrent disease, and samples collected across distinct treatment settings. Specifically, tumor grade was available for 25 cases, TNM stages for 21 cases, and the cohort included 23 primary and 17 recurrent tumors, as well as samples from BCG-naïve (n = 14), BCG-treated (n = 14), epirubicin-naïve (n = 4), and epirubicin-treated (n = 4) settings (**Table S15-S20**). Matched whole-exome sequencing (WES) of paired tumor and health tissue specimens was available from 23 patients (**Table S6-S10**). To expand the spatial coverage across diverse histopathological contexts, we further integrated nine publicly available Visium datasets comprising 57 bladder tissue samples and 256,531 spatial spots. After quality control, requiring at least 500 counts and 300 detected genes per spot, 214, 326 spots were retained. These samples included 53 tumors, three adjacent normal tissues, and one healthy bladder tissue, with the tumor specimens encompassing pure urothelial carcinoma as well as squamous, glandular, micropapillary, and sarcomatoid variants (**Table S21-S27**). To complement these public datasets with matched spatial multi-omics measurements, we generated spatial transcriptomic and spatial metabolomic profiles from three pairs of tumor and adjacent tissue sections (**Table S1**) and performed multiplex immunofluorescence (mIF) profiling across 14 BLCA specimens using multiple antibody panels. An independent tissue microarray (TMA) comprising 96 samples from 63 patients provided an additional cohort for spatial validation (**Table S28**).

**Figure 1.**
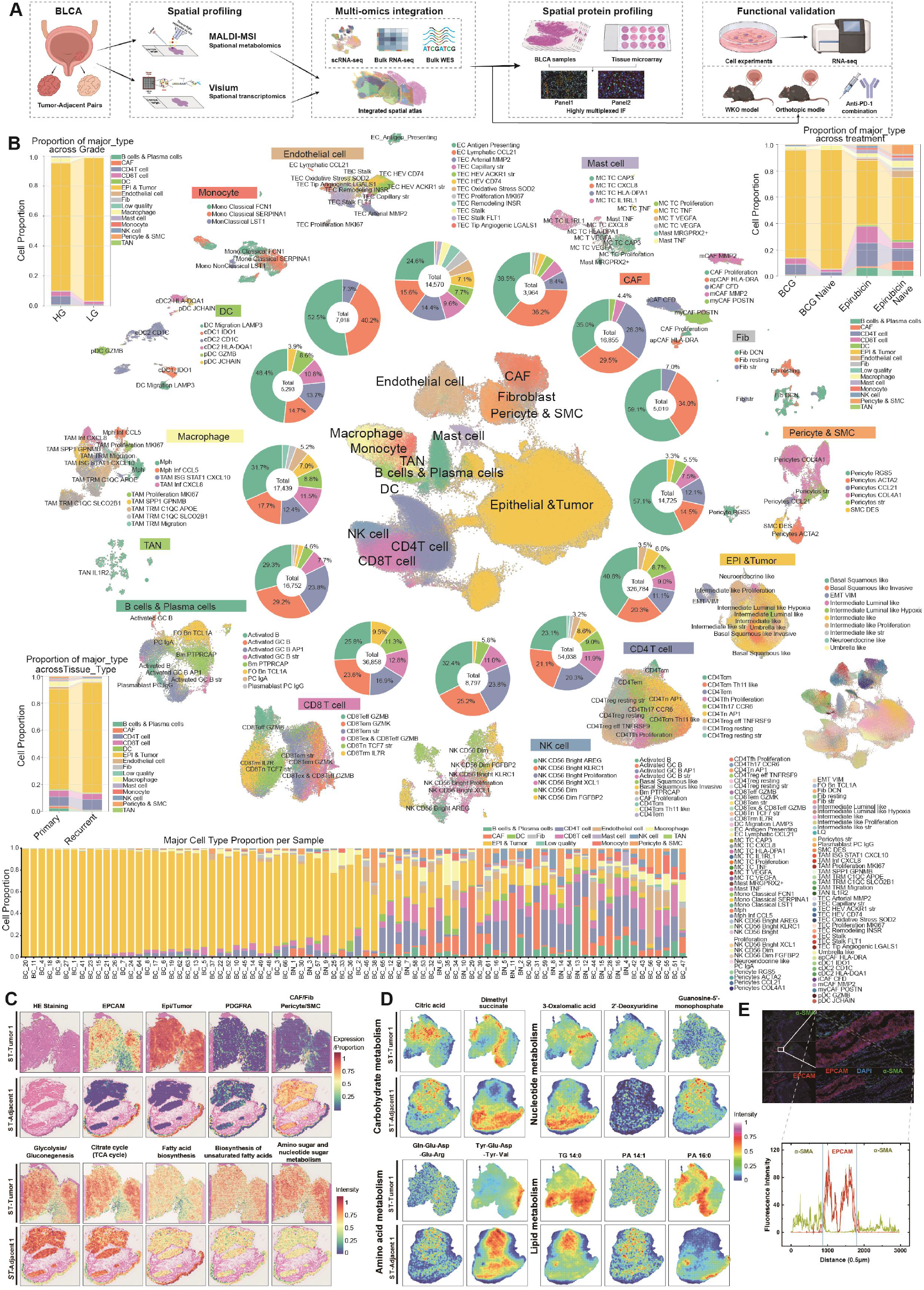
Single-cell, spatial transcriptomic, and spatial metabolic landscapes of BLCA. (A) Schematic overview of study design and analytical workflow, including data sources, cell subset identification, spatial organization, spatial metabolic profiling, metabolic reprogramming analysis, and combination immunotherapy strategies. (B) UMAP plots showing major cell clusters and subsets in BLCA versus normal tissues, with proportional distributions across high-grade versus low-grade, primary versus recurrent, different treatment modalities, and all samples. (C) Spatial maps showing expression of gene signatures in BLCA versus adjacent tissues. (D) Spatial maps showing distributions of different metabolite types in BLCA versus adjacent tissues. (E) Spatial distribution of stromal and tumor compartments in BLCA tissues.

Unsupervised analysis of the scRNA-seq compendium resolved the major cellular lineages of the BLCA ecosystem, including epithelial/tumor cells, fibroblasts, endothelial cells, myeloid cells, and lymphoid cells, which were further stratified into transcriptionally distinct states (**Figure 1B; Figures S1B** and **S1C**). Comparative compositional analyses revealed substantial variation in cell-state representation according to tumor grade, primary versus recurrent disease, and treatment status, indicating marked differences of the cellular ecosystem across clinical contexts (**Figure 1B**).

Spatial transcriptomics further revealed regionalized epithelial, stromal, and immune transcriptional programs that aligned with the tissue architecture defined by paired H&E staining (**Figure 1C; Figure S1D**). Complementary spatial metabolomics demonstrated pronounced spatial heterogeneity in metabolite-associated ion distributions between tumor and adjacent regions, including features associated with amino acid, lipid, and tricarboxylic acid (TCA) cycle metabolism (**Figure 1D; Figure S1E**). Segmentation of regions of interest further delineated tumor and stromal compartments within individual BLCA sections, enabling integrated interrogation of transcriptional and metabolic features across spatially defined tissue regions (**Figure 1E**). This multimodal resource established a spatially and molecularly resolved foundation for investigating tumor stromal organization, metabolic remodeling, and immune states in BLCA.

### Spatially coupled C1 CAF and E2 tumor epithelial states mark clinically aggressive BLCA

To identify cellular compartments most extensively remodeled in BLCA, we quantified transcriptional perturbation across major cell populations. Epithelial cells and CAFs were among the most transcriptionally altered compartments between adjacent and tumor tissues, with CAF-associated differentially expressed genes prominently enriched for extracellular matrix organization (**Figure 2A**). We therefore focused on resolving epithelial and stromal states and their spatial relationships.

**Figure 2.**
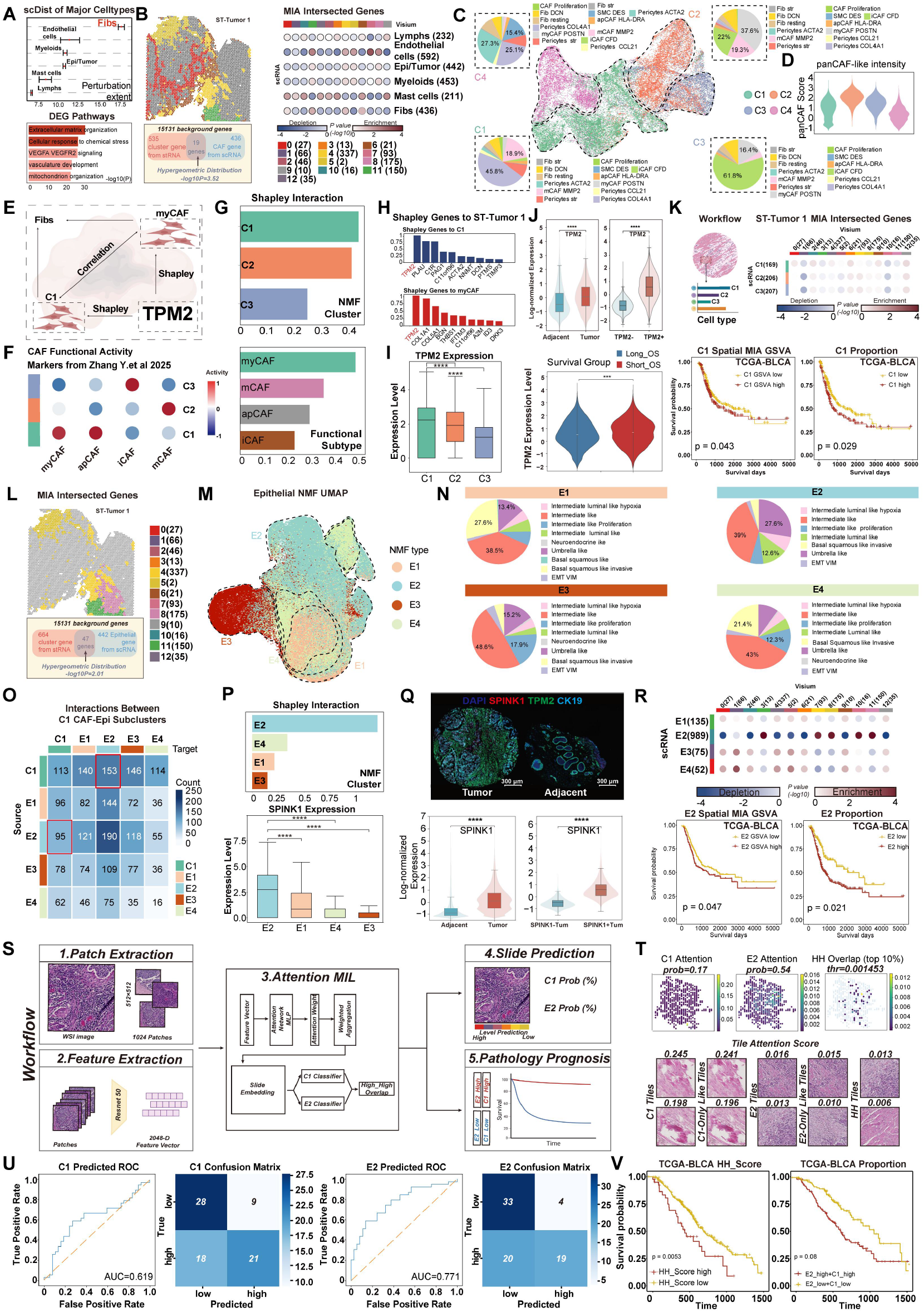
Spatial and clinical analyses prioritize C1 CAF and E2 epithelial states in BLCA. (A) Cell-state perturbation between adjacent and BLCA tissues, with pathway enrichment of CAF-associated differentially expressed genes. (B) Multimodal intersection analysis (MIA) linking CAF signatures to spatial clusters in ST-Tumor 1. (C) UMAP of NMF-defined CAF states C1–C4 and their functional-state composition. (D) panCAF scores across CAF states. (E) SHAP workflow evaluating the contributions of CAF states, functional programs, and gene-level features to the C1–myCAF pattern. (F) myCAF, apCAF, iCAF, mCAF, and panCAF module scores across C1–C3. (G) SHAP contributions of CAF states and functional programs to spatial CAF abundance. (H) Top MIA derived genes ranked by their SHAP contributions to the spatial C1 and myCAF patterns. (I) TPM2 expression across C1–C3. (J) TPM2 fluorescence intensity in adjacent and tumor TMA tissues and TPM2-defined groups. (K) Spatial MIA workflow, C1 enrichment across Visium regions, and overall survival according to the C1 MIA-GSVA score and inferred C1 abundance. (L) MIA linking epithelial signatures to spatial clusters in ST-Tumor 1. (M) UMAP of epithelial states E1–E4. (N) Composition of annotated epithelial populations within E1–E4. (O) CellChat-inferred ligand–receptor interactions among C1 CAFs and E1–E4 epithelial states. (P) SHAP contributions of epithelial states to spatial epithelial abundance and SPINK1 expression across E1–E4. (Q) Representative mIF of SPINK1, TPM2, and CK19, with SPINK1 fluorescence intensity quantified in adjacent, tumor, and SPINK1-defined regions. (R) Spatial MIA enrichment of epithelial states and overall survival according to the E2 MIA-GSVA score and inferred E2 abundance. (S) Attention-based multiple-instance learning workflow predicting C1 and E2 abundance from TCGA-BLCA H&E slides. (T) Representative C1, E2, and joint attention maps and high-attention tiles. (U) ROC curves and confusion matrices for C1 and E2 classification. (V) Overall survival according to joint C1–E2 attention and concordant C1/E2 abundance. Survival differences were assessed using two-sided log-rank tests; all other comparisons used two-sided Wilcoxon tests. Exact P values and scale bars are indicated.

Given that CAF phenotypes are influenced by their local tumor–stroma context, particularly their close spatial association with malignant epithelial cells, we sought to incorporate spatial relevance into feature selection rather than relying solely on highly variable genes derived from dissociated scRNA-seq data. Multimodal intersection analysis (MIA) was first used to identify spatial regions enriched for scRNA-seq-derived fibroblast signatures. Marker genes shared between these CAF-enriched spatial regions and scRNA-seq-defined fibroblasts were then retained as a spatially informed CAF feature set, thereby prioritizing genes associated with both fibroblast identity and their spatial organization within the tumor. NMF based on this feature set resolved CAFs into four transcriptional states (C1–C4) (**Figures 2B** and **2C**). Among these, C4 exhibited the lowest pan-CAF module score, indicating a comparatively attenuated canonical CAF program (**Figure 2D; Figures S2A** and **S2B**).

Among the remaining states, C1 combined the strongest myCAF program with perivascular fibroblast features, consistent with a spatially integrated stromal state surrounding the tumor vascular (**Figure 2E)**. SHAP modeling prioritized both C1 and the myCAF program in spatial CAF enrichment, with *TPM2* emerging as a leading gene associated with molecular and spatial patterns (**Figures 2F–2H**). Consistently, *TPM2* expression was highest in C1 relative to C2 and C3, nominating TPM2 as a marker of the C1 myCAF-like state (**Figure 2I**). Multiplex imaging confirmed higher TPM2 fluorescence intensity in tumor than in adjacent tissue (**Figure 2J**). A C1-associated signature derived by intersecting C1 differentially expressed genes with spatially associated genes was subsequently evaluated in TCGA-BLCA. Both inferred C1 abundance and the corresponding GSVA score were associated with shorter overall survival, linking the C1 state to adverse clinical outcome (**Figure 2K; Figure S2D**).

Applying an analogous integrative framework to the tumor epithelial compartment resolved four transcriptional states (E1–E4) with distinct epithelial programs (**Figures 2L–2N**). CellChat predicted more ligand–receptor interactions between C1 and E2 than between C1 and the other epithelial states, nominating preferential communication between these populations (**Figure 2O**). SHAP analysis subsequently identified *SPINK1* as a prominent E2-associated feature, and E2 displayed the highest *SPINK1* expression among the epithelial states (**Figure 2P**). Multiplex imaging further demonstrated the tissue distributions of CK19, SPINK1, and TPM2 and confirmed higher SPINK1 fluorescence intensity in tumor than in adjacent regions (**Figure 2Q**). Across additional BLCA cohorts, C1 and E2 were enriched in tumors and associated with high-grade and recurrent disease (**Figure S2C; Table S11-S14**). E2 spatial enrichment and inferred E2 abundance were also associated with shorter overall survival in TCGA-BLCA, with concordant survival associations observed across three independent cohorts (**Figure 2R; Figure S2D**).

We next asked whether the C1–E2 association was reflected in tissue morphology. An attention-based multiple instance learning model applied to TCGA-BLCA H&E whole-slide images classified tumors with high or low inferred C1 and E2 abundance (**Figure 2S**). Attention maps identified partially overlapping histological regions associated with the two states, and representative tiles illustrated morphological patterns contributing to individual and joint attention (**Figure 2T**). The C1 and E2 classifiers achieved AUCs of 0.619 and 0.771, respectively, exceeding those of the corresponding logistic regression baselines (0.579 and 0.667). respectively, and outperformed mean-pooled feature baselines (**Figure 2U**). Notably, concurrent high C1 and E2 abundance inferred by transcriptomic deconvolution showed only a trend toward shorter survival (P = 0.08), whereas a pathological image-derived C1–E2 co-attention score robustly stratified patient outcomes (P = 0.0053; **Figure 2V**). Thus, coupling MIA-defined spatial states with histopathology captured clinically relevant tissue architecture beyond that represented by expression-derived abundance alone, highlighting the prognostic significance of the spatial C1–E2 niche.

Pathway analysis provided additional molecular context for these states. ESCAPE and PROGENy identified enrichment of MYC targets, G2M checkpoint, mitotic spindle, PI3K/AKT/mTOR, WNT/β-catenin, NOTCH, and HEDGEHOG programs in C1, together with increased inferred TNF, NF-κB, TGF-β, and EGFR pathway activity (**Figure S3A**). E2 was enriched for fatty acid metabolism, hypoxia, PI3K/AKT/mTOR signaling, and protein secretion, with higher inferred TNF, NF-κB, and EGFR activity. Regulon analyses further associated HIF1A activity with E2 and ATF4 activity with C1. These state-specific regulatory programs were supported by scRegulate and spatially localized to the corresponding epithelial and stromal regions by SpaGRN (**Figure S3B**).

Integrated pathway and regulon analyses nominated HIF1A, NF-κB, and FOSB as candidate regulators of protein secretion programs and *SPINK1* expression in E2 tumor cells. In C1 CAFs, inferred EGFR/PI3K/AKT activity was accompanied by ATF4, MYC, and TGF-β associated programs (**Figure S3C)**. Complementary analysis integrating WES data revealed a higher E2 cell fraction in FGFR3-mutant tumors and elevated *SPINK1* expression in the mutation-positive groups analyzed (**Figure S3D**). Collectively, these findings identify spatially associated SPINK1-high E2 epithelial and TPM2-high C1 myCAF-like populations linked to aggressive BLCA and adverse clinical outcomes.

### SPINK1 and TPM2 contribute to tumor and stromal state maintenance during BLCA progression

The spatial association between tumor cells expressing high levels of *SPINK1* and CAFs expressing high levels of *TPM2* prompted us to examine the functional contributions of these molecules in their respective compartments. In J82 and T24 cells, *SPINK1* knockdown and antibody neutralization reduced cell viability and EdU incorporation, whereas recombinant SPINK1 increased both readouts (**Figures 3A** and **S4A**). These complementary perturbations support a contribution of extracellular SPINK1 to BLCA cell proliferation. Recombinant SPINK1 also increased PI3K, AKT, and mTOR phosphorylation, whereas SPINK1 knockdown reduced phosphorylation of these proteins (**Figure 3B**), linking SPINK1 availability with activity of this signaling pathway.

**Figure 3.**
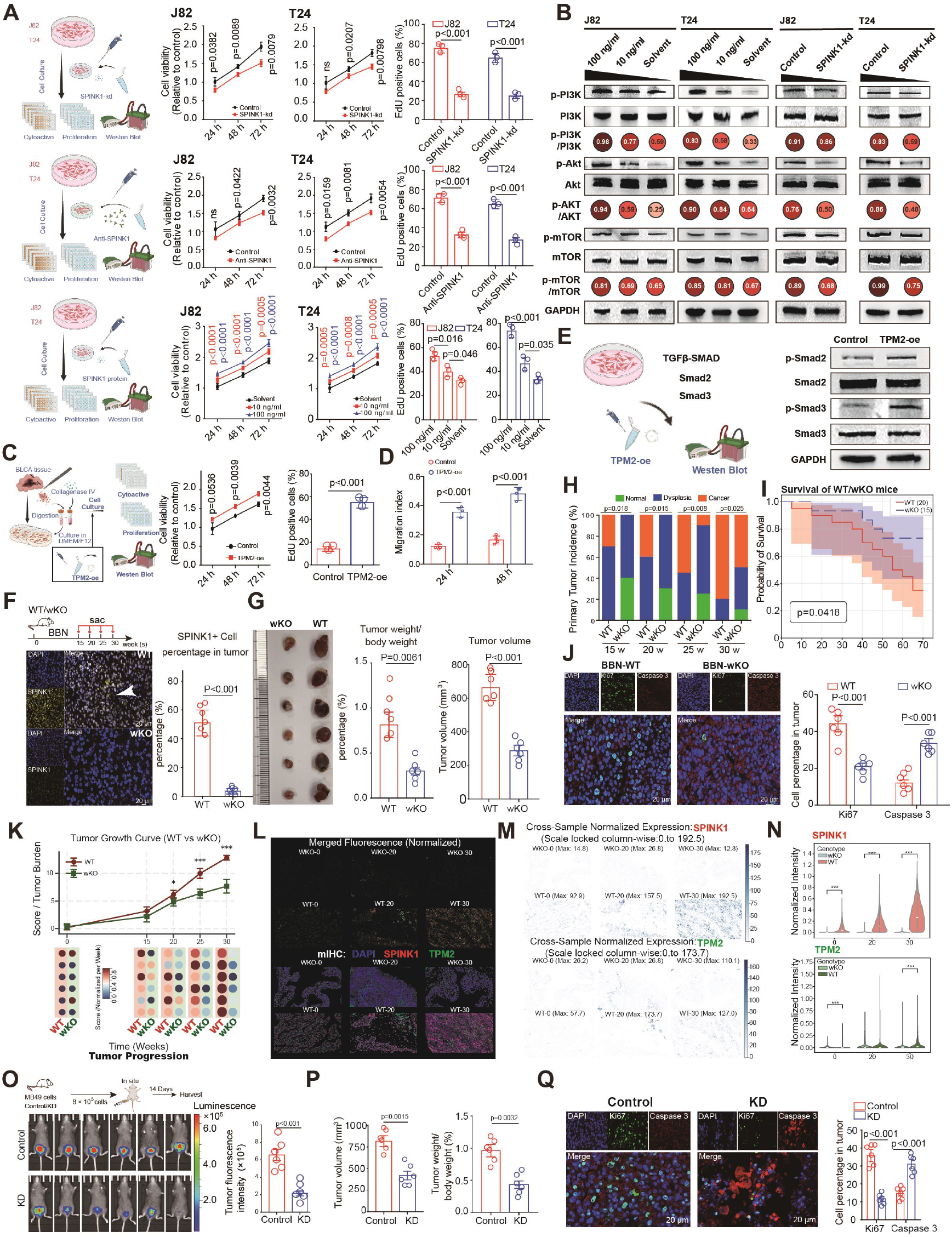
SPINK1 and TPM2 reinforce tumor and stromal fitness during BLCA progression. (A) Viability and EdU incorporation in J82 and T24 cells following *SPINK1* knockdown, neutralization, or recombinant SPINK1 treatment (10 or 100 ng/mL). (B) Immunoblots of phosphorylated and total PI3K, AKT, and mTOR following recombinant SPINK1 treatment or *SPINK1* knockdown. (C) Viability and EdU incorporation in control and *TPM2*-overexpressing patient derived CAFs. (D) CAF migration indices at 24 and 48 h. (E) Immunoblots of phosphorylated and total SMAD2/3 following *TPM2* overexpression. (F) BBN induced carcinogenesis in wild-type (WT) and whole-body Spink1 knockout (wKO) mice, with immunofluorescence and quantification of SPINK1 positive cells. (G) Gross tumor images, tumor/body weight ratios, and tumor volumes in WT and wKO mice. (H) Histopathological distributions of normal mucosa, dysplasia, and carcinoma at 15, 20, 25, and 30 weeks. (I) Kaplan–Meier survival curves following BBN exposure. (J) Immunofluorescence and quantification of Ki67-positive and Caspase-3-positive cells in BBN-induced tumors. (K) Tumor burden scores over time and corresponding progression dot plots. (L) SPINK1/TPM2 fluorescence images at 0, 20, and 30 weeks. DAPI, blue; SPINK1, red; TPM2, green. (M) Spatial SPINK1/TPM2 intensity maps with cross-sample normalization and a common scale for each marker. (N) Distributions of normalized SPINK1 and TPM2 fluorescence intensity across genotypes and time points. (O) Orthotopic implantation of control or *Spink1*-knockdown MB49 cells, with bioluminescence imaging and quantification at day 14. (P) Tumor volumes and tumor weight-to-body weight ratios in the MB49 model. (Q) Immunofluorescence and quantification of Ki67-positive and Caspase-3-positive cells in MB49 tumors. Statistical significance was assessed using two-sided Student’s t-tests for experimental group comparisons, the log-rank test for survival analysis, and the Wilcoxon test for fluorescence-intensity distributions. P values and significance annotations are shown in the panels. Scale bars are indicated in the images.

In CAFs isolated from patient tumors, *TPM2* overexpression increased cell viability and EdU incorporation (**Figures 3C** and **S4A**). It also accelerated scratch wound closure at 24 and 48 h (**Figures 3D** and **S4A**). Immunoblotting revealed increased Smad2 and Smad3 phosphorylation after *TPM2* overexpression (**Figure 3E**), indicating that the proliferative and wound closure phenotypes were accompanied by changes in SMAD signaling. We next tested whether *Spink1* contributed to bladder tumor development *in vivo* using BBN (N-Butyl-N-(4Hydroxybutyl) nitrosamine) exposure in *WT* and *Spink1-wKO* mice. Genotyping confirmed the knockout alleles, and tissue imaging demonstrated reduced SPINK1 staining in *wKO* tumors (**Figures 3F, S4B** and **S4C**). *Spink1-wKO* mice developed smaller tumors and had lower tumor weight to body weight ratios than WT controls (**Figure 3G**). Histopathological assessment showed a lower proportion of carcinoma lesions at the examined time points, consistent with delayed progression (**Figures 3H** and **S4D**; **Table S5**). This reduction in disease severity was accompanied by prolonged survival, lower Ki67⁺ fractions, increased Caspase-3 staining, and lower tumor-burden scores at later time points (**Figures 3I**-**3K**). Multiplex imaging identified SPINK1 and Ki67 co-expression in WT tumors (**Figure S4E**).

Multiplex imaging at baseline and weeks 20 and 30 showed increasing SPINK1 signal in WT tissues during tumorigenesis. *Spink1-wKO* tissues exhibited reduced SPINK1 staining, accompanied by reduced TPM2 intensity and altered organization of the TPM2 positive stromal compartment (**Figures 3L–3N**). Alongside these stromal changes, *Spink1-wKO* mice exhibited a progressive increase. As tumors progressed, *Spink1-wKO* mice exhibited a progressive increase in both the proportion and intratumoral infiltration of CD8⁺ T cells compared with WT controls (**Figures S4F-S4I**). Thus, reduced tumor burden was accompanied by changes in epithelial and stromal marker patterns.

To isolate the contribution of tumor-cell *Spink1* from systemic *Spink1* loss, we orthotopically implanted MB49 cells with or without *Spink1* knockdown (**Figures 3O, S4J**). At day 14 after implantation, knockdown tumors showed lower bioluminescence, tumor volume, and tumor weight to body weight ratios (**Figures 3O** and **3P**; **Figure S4K**). Reduced SPINK1 staining was accompanied by lower Ki67⁺ fractions and increased Caspase-3 staining (**Figures 3Q**, **S4L,** and **S4M**). Together, these complementary models establish a tumor-promoting role for SPINK1 in BLCA and show that systemic *Spink1* loss is accompanied by remodeling of TPM2 positive stromal regions and increased CD8⁺ T-cell infiltration.

### Spatial juxtaposition of SPINK1⁺ tumor cells and TPM2⁺ CAFs defines a structured tumor–stromal niche in BLCA

Building on our previous observation of spatial association between SPINK1⁺ epithelial cells and TPM2⁺ CAFs, we examined the organization of E2 epithelial cells and C1 CAFs within BLCA tissues. RCTD deconvolution of spatial transcriptomic data revealed regional co-enrichment of E2 and C1, while BANKSY identified spatial domains containing both populations (**Figures 4A** and **4B**). E2 and C1 further exhibited highly similar spatial abundance profiles (cosine similarity = 0.89), suggesting that these states preferentially occupy related tissue regions.

**Figure 4.**
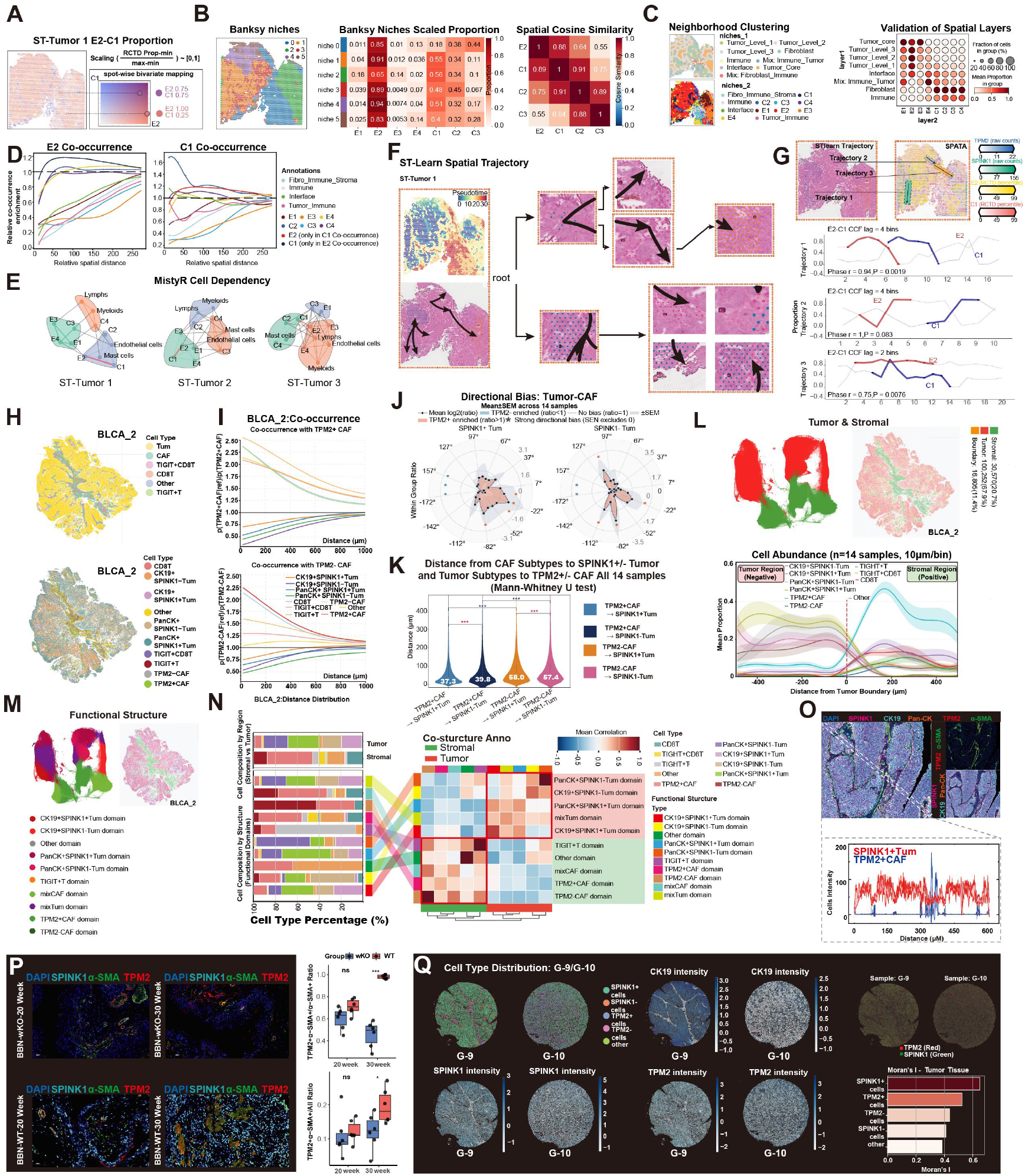
Spatial profiling defines a SPINK1⁺ tumor and TPM2⁺ CAF niche in BLCA. (A) RCTD deconvolution and bivariate mapping show inferred E2 tumor and C1 CAF proportions and their spatial distributions in ST-Tumor 1. (B) BANKSY niches, scaled E1–E4 epithelial and C1–C3 CAF proportions, and spatial cosine similarities among E2 and C1–C3. (C) Hierarchical neighborhood clustering of spatially smoothed cell state proportions annotates tissue compartments; the dot plot summarizes tumor cells and CAF representation across spatial layers. (D) Distance dependent co-occurrence around E2 tumor cells and C1 CAFs in ST-Tumor 1. (E) MISTyR networks summarize spatial predictive relationships among tumor cells, CAF, immune, and endothelial states across three BLCA spatial transcriptomic samples. (F) stLearn pseudotime and inferred trajectories in ST-Tumor 1, with corresponding H&E regions. (G) stLearn and SPATA trajectory comparison, E2 and C1 proportion profiles along three paths, and cross correlation lags. (H) mIF maps in BLCA_2 show major cell classes and tumor and CAF phenotypes resolved using SPINK1, pan-CK, CK19, TPM2, and α-SMA. (I) Spatial co-occurrence of major classes and resolved phenotypes around TPM2⁺ CAFs in BLCA_2. (J) Polar plots compare directional TPM2⁺ and TPM2⁻ CAF distributions relative to SPINK1⁺ or SPINK1⁻ tumor populations, showing mean log₂ratios ± SEM. (K) Nearest neighbor distance distributions from TPM2⁺ or TPM2⁻ CAFs to SPINK1⁺ or SPINK1⁻ tumor cells. (L) Tumor and stromal compartments in BLCA_2 and mean cell proportions across the boundary in 10 μm bins; negative and positive distances indicate tumor and stroma. (M) BANKSY spatial domains in BLCA_2 annotated by constituent tumor, CAF, and immune phenotypes. (N) Compartment and domain compositions, domain grouping, and correlations between composition profiles. (O) Representative mIF images and line scan intensity profiles show SPINK1⁺ tumor and TPM2⁺ CAF positioning at the tissue interface. (P) Representative mIF images of Spink1, Tpm2, and α-SMA in WT and Spink1-wKO BBN tumors at weeks 20 and 30. Boxplots quantify TPM2⁺α-SMA⁺ proportions among α-SMA⁺ and all cells. (Q) Paired tumor (G-9) and adjacent tissue (G-10) TMA cores show cell distributions, CK19, SPINK1, and TPM2 intensities, and SPINK1/TPM2 overlays. Moran’s I summarizes spatial autocorrelation of individual populations in tumor tissue. J–L summarize 14 mIF samples; P values in J were calculated using Mann–Whitney U tests (two sided).

To resolve the tissue architecture underlying this association, we developed a hierarchical neighborhood clustering framework based on spatially smoothed cell state proportions. At the first level, neighborhoods were classified into tumor, fibroblast, immune, interface, and mixed regions, with tumor regions further subdivided according to their spatial organization (**Figure S5**). At the second level, neighborhoods were characterized by the dominant tumor states (E1–E4) and CAFs (C1–C4) states, delineating distinct stromal, epithelial, and mixed niches. This hierarchy localized neighborhoods enriched for C1 and E2 to adjacent stromal and tumor compartments, respectively, providing a tissue-level framework for describing the spatial relationship between these states (**Figure 4C**). Distance-dependent co-occurrence analysis further revealed enrichment of C1 CAFs around E2 epithelial cells and reciprocal enrichment of E2 cells around C1 CAFs across a range of spatial distances (**Figure 4D**), supporting their organization within a shared tumor–stromal neighborhood.

This spatial relationship was reproducible across tumors. MISTy analysis identified recurrent spatial dependencies between E2 epithelial cells and C1 CAFs across three BLCA samples (**Figure 4E**). Spatial trajectory analysis using STlearn further revealed coordinated variation in E2 and C1 abundance across the tissue landscape (**Figure 4F**). Along these trajectories, E2 and C1 abundance peaked at distinct but neighboring positions, consistent with an ordered spatial arrangement of the two states (**Figure 4G**). SPINK1 and TPM2 expression varied correspondingly with the E2 and C1 profiles, respectively (**Figure 4G)**.

To assess these relationships at the protein level, we generated mIF datasets and performed single-cell phenotyping (**Figures S6A** and **S6C**). Spatial mapping revealed cellular arrangements consistent with the transcriptomic observations (**Figure 4H; Figures S6B** and **S6D**). At the tissue-domain level, E2 epithelial and C1 CAF compartments exhibited reciprocal enrichment across intermediate spatial distances. At the cellular level, SPINK1⁺ tumor cells showed reduced short-range co-occurrence with TPM2⁺ CAFs in the BLCA_2 sample, with co-occurrence ratios approaching unity as distance increased (**Figure 4I**), indicating limited direct cellular intermixing. Directional analyses further revealed sector-specific heterogeneity in the relative distribution of TPM2⁺ and TPM2⁻ CAFs around tumor cells (**Figure 4J; Figure S6H**). Nevertheless, TPM2⁺ CAFs were significantly closer to SPINK1⁺ tumor cells than TPM2⁻ CAFs across the 14 samples (37.3 versus 58.0 μm; **Figure 4K**), supporting preferential proximity within a spatially compartmentalized tumor–CAF interface.

Mapping cell abundance across the tumor–stroma boundary localized TPM2⁺ CAFs to stromal regions adjacent to compartments enriched for SPINK1⁺ tumor cells, supporting their organization along the tissue interface (**Figure 4L**). The protein measurements therefore provided complementary evidence for the spatial association of the epithelial and stromal populations identified by transcriptomic analysis.

To further characterize microenvironmental architecture, we applied BANKSY across the mIF cohort and identified 35 spatially coherent domains. These domains were organized into tumor and stromal compartments and 10 recurring cellular neighborhoods (**Figure 4M**; **Figure S6E**). Neighborhood assignments agreed with the broader compartmental structure. Cell composition analysis identified multiple domains jointly enriched for SPINK1⁺ tumor cells and TPM2⁺ CAFs (**Figure 4N; Figure S6F**). Domain composition showed positive correlations within compartments and negative correlations between tumor and stromal domains (**Figure S6G**). Representative mIF images showed SPINK1⁺ tumor cells and TPM2⁺ CAFs in close proximity, approximately 20 – 50 μm apart (**Figure 4O**).

To examine changes during tumorigenesis, we revisited the BBN-induced BLCA model and quantified CAF abundance across disease stages. CAF abundance increased from week 20 to week 30 alongside increasing tumor burden (**Figure 4P**). Immunofluorescence also showed spatial proximity between SPINK1⁺ tumor cells and TPM2⁺ stromal cells in mouse tissues (**Figure S6I**).

In an independent BLCA TMA cohort, we performed analogous unsupervised clustering and cell annotation (**Figures S7A–S7C**). TPM2⁺ cells were more prevalent than TPM2⁻ cells within neighborhoods surrounding SPINK1⁺ cells, and SPINK1⁺ neighborhoods showed greater relative representation of TPM2⁺ cells than SPINK1⁻ neighborhoods (**Figures S7D** and **S7F**). Compared with adjacent tissues, tumor specimens showed higher SPINK1 and TPM2 fluorescence intensities and increased signals for markers associated with metabolism (CD98, SLC16A3) and T cell dysfunction (TIGIT) (**Figure S7E**). Moran’s I analysis identified spatial autocorrelation of SPINK1 and TPM2 signals, indicating clustering of each marker within tissue (**Figure 4Q**). BANKSY independently resolved tumor–stromal compartments and finer cellular neighborhoods consistent with the observed distribution patterns (**Figures S7G–S7J**). This independent cohort supported the spatial association between SPINK1⁺ tumor cells and TPM2⁺ CAFs observed in the discovery samples, defining a reproducible spatial niche in BLCA.

### SPINK1–EGFR– and TGF-β-associated signaling coordinates reciprocal tumor– stromal interactions

Having established the preferential spatial association between SPINK1⁺ tumor cells and TPM2⁺ CAFs, we investigated their functional interactions. We established complementary Transwell co-cultures using BLCA cells overexpressing *SPINK1* or CAFs overexpressing *TPM2* (**Figure 5A**). The two configurations separately assessed the effects of tumor cells on patient derived CAFs and of CAFs on tumor cells. Tumor cells with *SPINK1* overexpression increased CAF viability and EdU incorporation, indicating enhanced fibroblast proliferation (**Figure 5B; Figure S8A**). Conversely, CAFs overexpressing *TPM2* enhanced J82 and T24 viability and proliferation (**Figure 5C; Figure S8B**), together with tumor cell migration and invasion assessed by wound-healing and Transwell assays (**Figures 5D** and **5E**; **Figures S8C** and **S8D**). Voronoi neighborhood analysis across 14 mIF samples revealed preferential connectivity between TPM2⁺ CAFs and SPINK1⁺ tumor cells within spatial adjacency networks (**Figure 5F**). These findings suggest potential reciprocal interactions between stromal and tumor cells.

**Figure 5.**
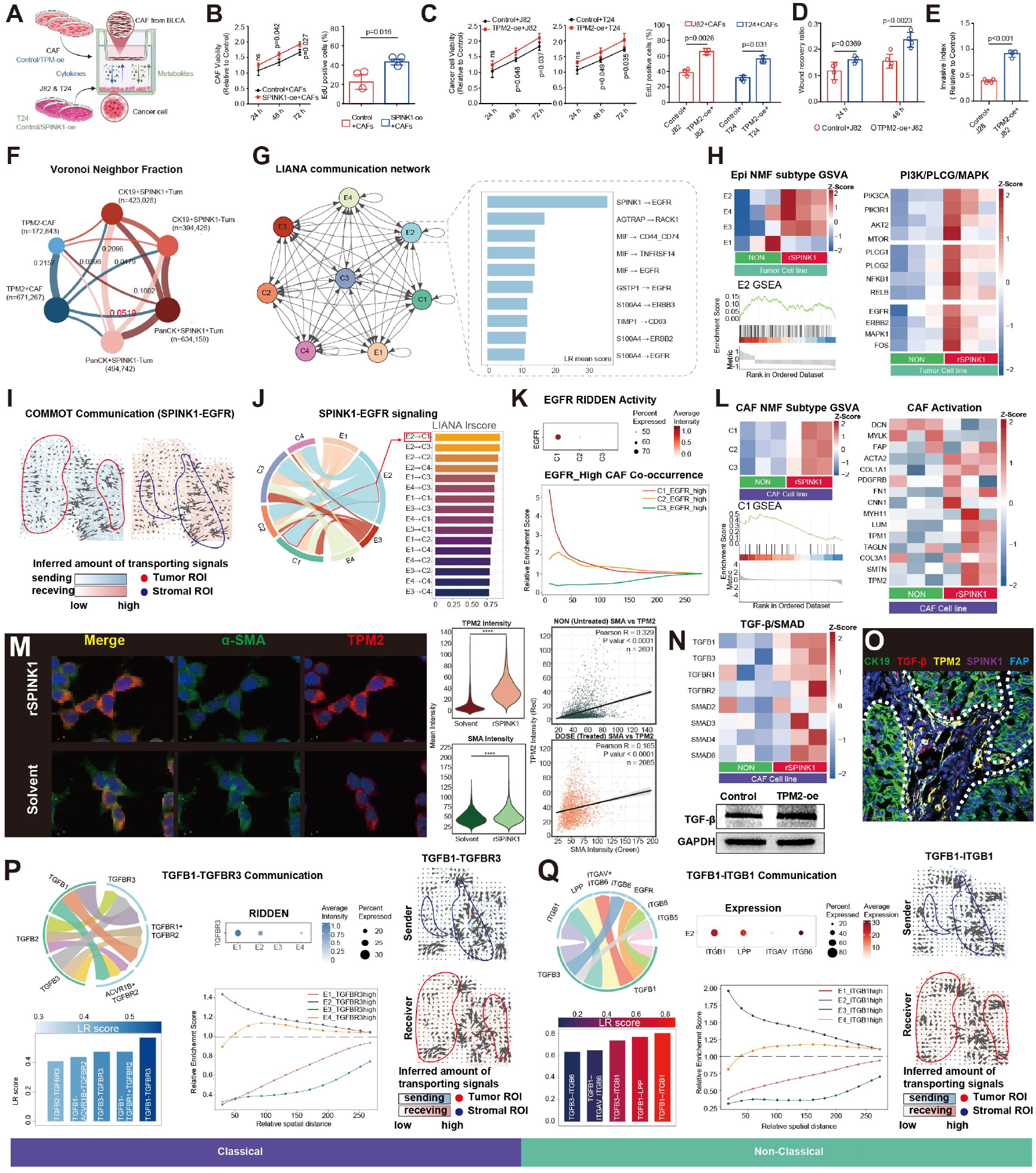
Functional and spatial analyses implicate SPINK1–EGFR and TGF-β signaling in tumor and CAF crosstalk. (A) Transwell coculture system assessing reciprocal effects of *SPINK1* overexpression in BLCA cells and *TPM2* overexpression in patient derived CAFs. (B) Patient derived CAF viability at indicated time points and EdU incorporation following coculture with control tumor cells or tumor cells overexpressing *SPINK1*. (C) J82 and T24 viability over time and EdU incorporation following coculture with control CAFs or CAFs overexpressing *TPM2*. (D) Quantification of wound closure in J82 cells cocultured with control CAFs or CAFs overexpressing *TPM2* at indicated time points. (E) Quantification of J82 cell invasion following coculture with control CAFs or CAFs overexpressing TPM2. (F) Voronoi neighborhood analysis across 14 mIF samples. Edge widths represent mean neighborhood fractions between tumor and CAF phenotypes. (G) Communication network inferred using LIANA among E1–E4 epithelial and C1– C4 CAF states. Ranked ligand–receptor interactions associated with E2 tumor cells highlight SPINK1–EGFR. (H) Transcriptional responses of BLCA cells to recombinant SPINK1, epithelial NMF state GSVA heatmap, GSEA of the E2 transcriptional program, and heatmap of PI3K/PLCγ/MAPK component expression. (I) Spatial SPINK1–EGFR communication inferred using COMMOT between annotated tumor and stromal ROIs. (J) Predicted SPINK1–EGFR communication shown by a chord diagram and LIANA scores ranking epithelial–CAF pairs. (K) EGFR activity inferred using RIDDEN across CAF states and spatial co-occurrence of EGFR-high C1–C3 CAFs relative to tumor cells. (L) Transcriptional responses of patient derived CAFs to rSPINK1, CAF NMF state GSVA heatmap, GSEA of the C1 transcriptional program, and heatmap of genes associated with CAF activation. (M) Representative immunofluorescence images of TPM2 and α-SMA in CAFs treated with vehicle or rSPINK1. Violin plots quantify fluorescence intensities; scatterplots show correlations between TPM2 and α-SMA intensities. (N) Heatmap of TGF-β ligand, receptor, and SMAD component expression following rSPINK1 treatment and TGF-β immunoblotting in control CAFs and CAFs overexpressing TPM2. (O) Representative mIF image showing CK19, TGF-β, TPM2, SPINK1, and FAP at the tumor–stromal interface in human BLCA tissue. Dashed lines indicate tumor– stromal boundaries. (P) Predicted TGFB1–TGFBR3 communication from CAFs to tumor cells: chord diagram and interaction scores, TGFBR3 activity inferred using RIDDEN, spatial enrichment of TGFBR3-high tumor states around C1 CAFs, and COMMOT sender and receiver maps. (Q) Predicted TGFB1–ITGB1 interactions from CAFs to tumor cells: chord diagram and ligand–receptor scores, ITGB1 expression, spatial enrichment of ITGB1-high tumor states around C1 CAFs, and COMMOT sender and receiver maps.

We next investigated molecular signals potentially underlying these interactions. LIANA prioritized SPINK1–EGFR as a candidate autocrine interaction within E2, suggesting a possible contribution to maintenance of the E2 program (**Figure 5G**). To model putative autocrine signaling in SPINK1⁺ bladder cancer cells, we treated T24 cells with recombinant SPINK1 (rSPINK1). Treatment increased enrichment of the E2 transcriptional signature and altered expression of genes associated with PI3K, PLCγ, and MAPK signaling (**Figure 5H**), consistent with our earlier signaling observations (**Figure 3B**).

We then examined whether SPINK1 could also influence the stromal compartment. COMMOT inferred a spatial pattern of SPINK1–EGFR communication linking tumor and stromal regions of interest, with predicted sending activity in tumor regions and receiving activity in adjacent stroma (**Figure 5I**). Although E2 was predicted to communicate with multiple CAF states, C1 was prioritized as a candidate receiver, supported by higher LIANA scores, a greater proportion of EGFR-expressing cells, and elevated receptor activity inferred by RIDDEN (**Figures 5J** and **5K**). EGFR-high C1 cells also showed the strongest co-occurrence at short spatial distances. RNA sequencing of primary CAFs treated with rSPINK1 revealed enrichment of C1 and myCAF transcriptional signatures (**Figure 5L**), supporting induction of C1/myCAF-like transcriptional programs. These observations extended the SPINK1-associated transcriptional response from the tumor compartment to CAFs, complementing the reciprocal co-culture phenotypes.

Additionally, rSPINK1 increased TPM2 and α-SMA expression, consistent with enhanced expression of myCAF markers (**Figure 5M**). The correlation between these markers decreased after treatment, despite the overall increase in their expression. rSPINK1 also altered expression of *TGFB1–3*, *TGFBR1/2*, and downstream SMAD-associated genes, consistent with engagement of a TGF-β-related transcriptional program (**Figure 5N**). Clinical samples mIF localized TGF-β and FAP signals to TPM2⁺ stromal regions adjacent to SPINK1⁺ tumor cells at the tumor–stroma interface (**Figure 5O**), providing tissue context for the CAF phenotype observed in culture.

Ligand–receptor analysis further nominated TGF-β-associated communication from CAFs to tumor cells. TGFB1–TGFBR3 emerged as a prominent candidate interaction, with TGFBR3-high E2 cells showing the strongest enrichment at short spatial distances (**Figure 5P**). A complementary analysis prioritized TGFB1–ITGB1, accompanied by high interaction scores and local enrichment of ITGB1-high E2 cells (**Figure 5Q**). Collectively, these findings support a model in which tumor-derived SPINK1 promotes a TPM2-high spatial CAFs program and reciprocal CAF activity enhances aggressive tumor-cell phenotypes. SPINK1–EGFR-associated signaling provides a candidate molecular link for tumor to stroma communication, whereas CAF-derived TGF-β associated signals including TGFB1–TGFBR3 and TGFB1–ITGB1 converge preferentially on E2 cells, identifying E2 as the predominant tumor-cell recipient of this proposed stromal to tumor feedback.

### Spatial multi-omics delineates E2 and C1 metabolic programs

To characterize the metabolic programs associated with the E2–C1 spatial niche, we first used METAFlux to derive metabolic subsystem profiles from single-cell transcriptomes. CAFs exhibited prominent amino acid, nucleotide, glutathione, and glycolysis/gluconeogenesis programs, with C1 showing high relative scores for amino acid and nitrogen metabolism. E2 and E3 displayed prominent fatty acid oxidation and cholesterol metabolism (**Figure 6A)**. Building on these broad profiles, complementary Compass analysis integrated gene expression with a curated metabolic network to further resolve differences in metabolic reaction potential. E2 exhibited higher scores for glycolysis/gluconeogenesis, fatty acid synthesis and oxidation, glycerophospholipid metabolism, and the pentose phosphate pathway than other epithelial states. C1 was distinguished from other CAF states by differences in CoA metabolism, the urea cycle, glutathione metabolism, and purine, nucleotide, and nitrogen metabolism (**Figure 6A**). Inferred reaction profiles distinguished glycolysis and pyruvate metabolism in E2 from broader glutamate synthesis and utilization in C1 (**Figure S9A**), suggesting distinct organization of shared metabolic programs.

**Figure 6.**
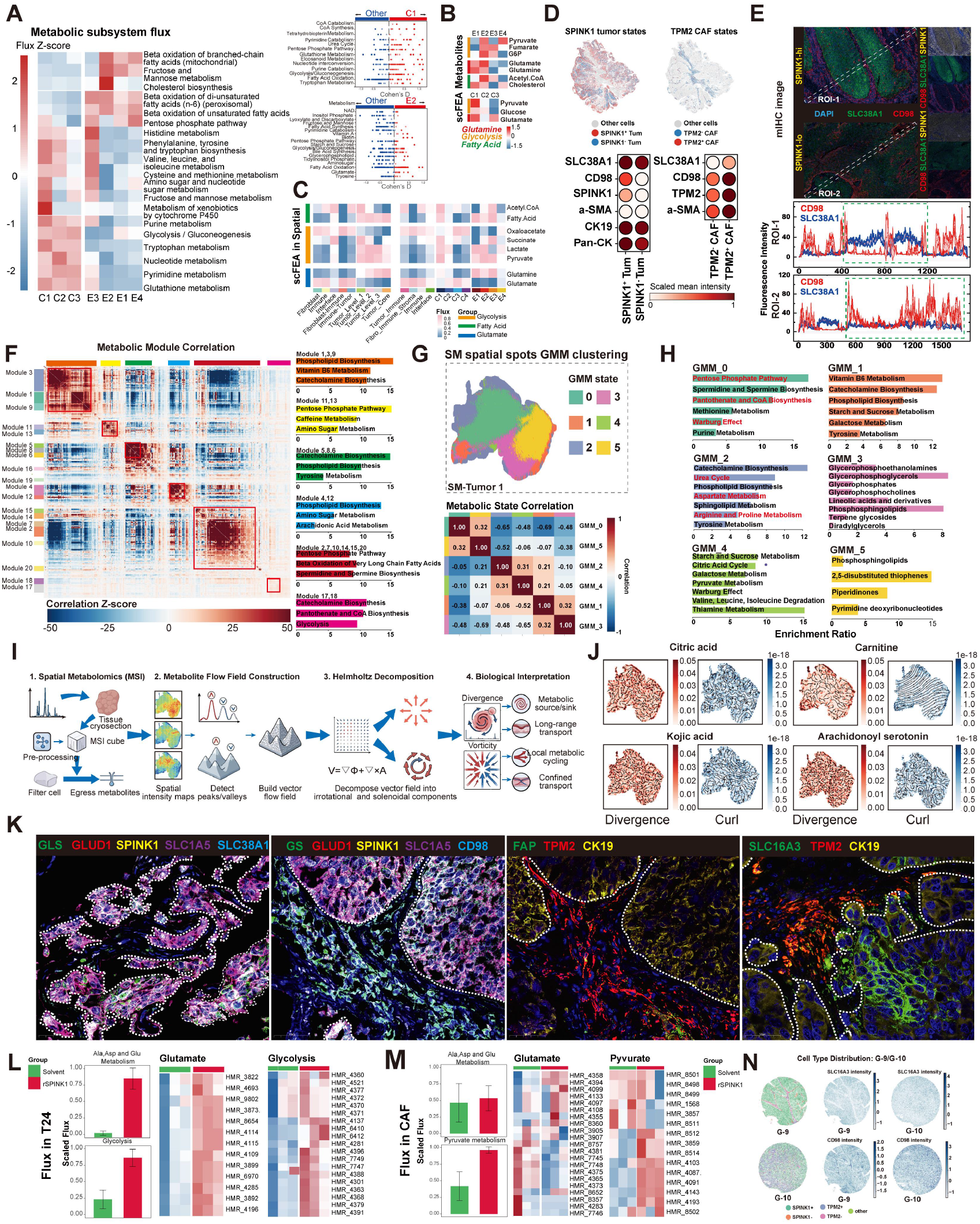
Tumor cells and CAFs exhibit complementary metabolic programs and distinct responses to SPINK1. (A) MetaFlux inferred metabolic subsystem scores across CAF and epithelial states (Z-score normalized). Compass comparisons of C1 versus other CAF states and E2 versus other epithelial states, expressed as Cohen’s d (red, positive toward C1/E2; blue, negative toward other states). (B) scFEA inferred relative metabolite scores across tumor and CAF states. (C) Heatmap of spatial transcriptome derived scFEA scores for acetyl-CoA, fatty acid, tricarboxylic acid cycle, glycolytic, and glutamine/glutamate-associated metabolism across regions assigned to major cell types and NMF states. (D) Spatial maps of cells classified as SPINK1 positive or SPINK1 negative tumor cells and TPM2 positive or TPM2 negative CAFs in the indicated BLCA specimen. The dot plot shows scaled mean intensities of SLC38A1, CD98, SPINK1, TPM2, α-SMA, CK19, and Pan-CK measured by highplex immunofluorescence. (E) Representative SPINK1-high and SPINK1-low regions of interest and fluorescence line-scan profiles of SLC38A1 and CD98 along the indicated transects. (F) Correlation matrix of ion features grouped into Hotspot modules, with pathway enrichment based on their putative metabolite annotations. (G) Spatial distribution of six GMM MALDI-MSI states in SM-Tumor 1 and pairwise correlations among state-averaged ion profiles. (H) Pathway enrichment based on putatively annotated ion features associated with GMM0 to GMM5. (I) Workflow for detecting peaks and valleys in MALDI-MSI ion-intensity maps, constructing intensity derived spatial vector fields, applying Helmholtz decomposition, and calculating divergence and curl descriptors. (J) Divergence and curl maps for ion features putatively annotated as citric acid, carnitine, kojic acid, and arachidonoyl serotonin. (K) Representative multiplex immunofluorescence images showing the compartmentalized distributions of GLS, glutamine synthetase (GS), GLUD1, SLC1A5, SLC38A1, CD98, and SLC16A3 relative to SPINK1, TPM2, FAP, and CK19. (L) Transcriptome-inferred HMR metabolic flux and reaction scores for alanine, aspartate, and glutamate metabolism and glycolysis in T24 cells treated with rSPINK1 or solvent. (M) Transcriptome-inferred HMR pathway and reaction scores for alanine, aspartate, and glutamate metabolism and pyruvate metabolism in patient derived CAFs treated with rSPINK1 or solvent. (N) Cell-type distributions and normalized SLC16A3 and CD98 intensities in paired G-9 tumor and G-10 adjacent tissue microarray cores.

To further characterize the metabolic transformations associated with these programs, we applied scFEA to single-cell transcriptomes. The resulting scores highlighted pyruvate and glutamate metabolism in both populations, with additional features involving glucose-6-phosphate, glutamine, acetyl-CoA, and cholesterol in E2 and glucose in C1 (**Figure 6B**). Extending scFEA to Visium transcriptomes delineated the spatial organization of these inferred metabolic programs. Regions enriched in E2 cells exhibited glycolytic and lipid metabolic features, whereas regions enriched in C1 CAFs showed glutamate metabolism and TCA cycle features, with heterogeneity within both regional compartments (**Figure 6C**). These spatial patterns reinforced the conclusions drawn from the single-cell dataset, supporting distinct transcriptome-inferred metabolic biases toward carbon and lipid metabolism in E2 and amino acid metabolism in C1.

We next examined whether these inferred programs were reflected in the spatial organization of metabolic proteins in BLCA tissues. SLC38A1 signals were enriched in tumor-associated regions, whereas CD98 (SLC3A2) showed prominent expression in TPM2⁺ CAFs (**Figure 6D**). Representative SPINK1-high fields showed more evident local covariation between SLC38A1 and CD98 than SPINK1-low fields (**Figures 6E**), while SLC38A1 tracked CK19 and PanCK, whereas CD98 occupied distinct but adjacent regions (**Figures S9B** and **S9C**).

### Bladder tumors harbor distinct metabolic territories with gradients across their boundaries

To relate transcriptome-inferred metabolic programs to the spatial distribution of metabolite-associated ions, we performed matrix-assisted laser desorption/ionization mass spectrometry imaging (MALDI-MSI). Profiling bladder tumors and adjacent tissues generated pixel-resolved ion maps (**Figures S9D** and **S9E**), with significant differences in aggregated spectra across approximately 650–900 m/z. Differential analyses identified tumor-associated ion features with putative links to tumor-associated metabolic pathways, including ether lipid metabolism, cysteine and methionine metabolism, and the pentose phosphate pathway whereas adjacent tissues displayed distinct metabolic associations (**Figures S9F–S9H**). Among individual features, putative immunosuppressive 5′-methylthioadenosine (MTA) and immunostimulatory kojic acid showed opposing tumor-associated patterns.

Hotspot identified 20 modules of covarying ion features in a Harmony-corrected representation of MALDI-MSI profiles (**Figure 6F; Figure S10A**). Projection of module scores onto tissue coordinates revealed regional variation in these profiles. These modules captured coordinated variation in ion abundance across tumor tissue regions. Gaussian mixture modeling (GMM) of module profiles resolved six states, selected by balancing silhouette performance and the Bayesian information criterion. GMM0–GMM5 separated in UMAP space and occupied restricted territories, with variable abundance and intermixing across tumors (**Figures S10B–S10D**). State-associated ions, observed-to-expected module scores, and correlations further distinguished their profiles (**Figure 6G; Figures S10E** and **S10F**).

Comparative analysis of serial sections revealed spatial correspondence between these metabolic territories and the E2–C1 tumor–stromal niche. GMM0/1 qualitatively overlapped E2-enriched territories, whereas GMM2 partly overlapped C1-enriched regions (**Figure 6H**). GMM0 showed associations with the pentose phosphate pathway, the Warburg effect, pantothenate and CoA biosynthesis, and methionine and purine metabolism, whereas GMM2 emphasized urea cycle, aspartate, arginine and proline, and sphingolipid pathways, suggesting greater representation of nitrogen metabolism, consistent with the transcriptomic findings. GMM1 highlighted vitamin B6, phospholipid, and carbohydrate metabolism; GMM3, glycerophospholipid pathways; GMM4, the TCA cycle, pyruvate and branched-chain amino acid metabolism, and the Warburg effect; and GMM5, phosphosphingolipid and pyrimidine deoxyribonucleotide pathways (**Figure 6H**). Spatial maps further showed regionalized amino acid, carbohydrate, lipid, and nucleotide programs (**Figures S10G-S10H**). These relationships indicate regional correspondence and suggest potential metabolic interactions within and between regions.

Among features annotated as key intracellular metabolites, GMM0/1 were enriched in CDP-diacylglycerol, ceramide, phosphoenolpyruvate, and putative PIP/PIP2 signals, whereas GMM2 showed cardiolipin, phosphatidylcholine, and phosphatidylethanolamine features (**Figure S11**). The phosphoinositide-associated pattern in GMM0/1 was concordant with the PI3K/AKT/mTOR transcriptional program associated with E2, suggesting potential activation of this signaling axis in these regions.

Finally, spatial intensity mapping and vector field analysis delineated regional and interfacial metabolic organization (**Figures 6I** and **6J**). Within GMM0/1 territories partly corresponding to E2 regions, citric acid intensity was higher and its standardized divergence lower in GMM0, whereas acylcarnitine and LPC intensities and carnitine divergence were higher in GMM1. These features provided additional spatial context for the glycolytic programs inferred in E2 (**Figure 6A**). GMM2, partly corresponding to C1 regions, showed higher intensities but lower divergence scores for 2-arachidonoylglycerol, lipoxin A4, and citrulline than GMM0/1. Kojic acid was prominent in GMM2/3, whereas arachidonoyl serotonin was most prominent in GMM3. Compared with GMM0–GMM2 transitions, GMM1–GMM2 transitions exhibited higher N-acetylleucine intensity, higher L-lysinamide divergence scores, and higher curl scores for LPC, LPE, platelet activating factor, and Arg/Lys peptides (**Figure S12A**). Collectively, these patterns suggest compartmentalized carbon, lipid, and nitrogen metabolism, with regional differences in metabolite availability and potential metabolic interactions at the E2–C1 interface.

### SPINK1 stimulation elicits distinct metabolic responses in tumor cells and CAFs

Having identified distinct metabolic programs across the E2–C1 tumor–stromal niche, we next asked whether SPINK1 stimulation altered metabolic features in tumor cells and CAFs. In T24 cells, rSPINK1 increased SREBP1 protein abundance and its nuclear-to-cytoplasmic ratio (**Figures S12B and S12C**). SREBP1 abundance and nuclear localization correlated positively with SPINK1-associated fluorescence, and line-scan analyses further supported increased nuclear accumulation of SREBP1 (**Figures S12D** and **S12E**). Evidence for SREBP1 transcript changes was less conclusive (P = 0.062; **Figure S12C**), placing the clearest response at the level of protein abundance and localization.

rSPINK1 also increased SLC1A5, GLUT, and MCT4 signals in T24 cells, with positive within-condition associations between SPINK1-associated fluorescence and SLC1A5 and coordinated variation in GLUT and MCT4 signals (**Figures S12F–S12I**). Patient derived CAFs showed a distinct transporter response, with increased LAT1/SLC7A5, SNAT2/SLC38A2, and MCT1/SLC16A1 signals (**Figure S12J**). Consistent with these protein-level changes, RNA-seq indicated rSPINK1-responsive programs involving lipid synthesis, glucose handling, glutamine metabolism, and lactate transport in T24 cells, including changes in *SLC1A5, SLC2A1, ACLY, SLC16A3,* and *GLUD1* (**Figure S12K**). In CAFs, rSPINK1 altered expression of genes including *SLC7A5, SLC38A2, SLC16A1, SLC3A2, GLUL,* and *GLS* (**Figure S12K**). These responses involved partially shared metabolic pathways, suggesting potential metabolic complementarity between tumor cells and CAFs. Serial section imaging further localized SLC38A1, SLC1A5, GLS, and GLUD1 preferentially to epithelial/tumor regions, together with tumor enrichment of SLC16A3/MCT4 and stromal enrichment of glutamine synthetase (**Figure 6K**; **Figures S13A–S13C**).

Transcriptome-derived reaction analyses further indicated higher aggregate scores for alanine/aspartate/glutamate metabolism and glycolysis in T24 cells (**Figure 6L**), paralleling selected transcriptional responses to rSPINK1 (**Figures S13D–S13F**), whereas CAFs showed a prominent pyruvate metabolism response (**Figure 6M**). Individual HMR reaction profiles further revealed heterogeneous changes in predicted flux within glutamate metabolism and glycolysis in T24 cells and within glycolysis and pyruvate metabolism in CAFs (**Figure S13G**), with concordant estimates of metabolic potential from MetaFlux (**Figure S13H**). Spatial profiling of representative tumor and adjacent TMA cores further demonstrated compartmentalized distributions of SLC16A3 and CD98 (**Figure 6N**). Across the broader TMA cohort, both proteins showed higher signals in TPM2 positive and SPINK1 positive populations while retaining distinct spatial biases (**Figure S13I**). Together, these SPINK1 responsive changes in SREBP1 and nutrient transporters, together with tumor enrichment of GLS, GLUD1, and MCT4 and stromal enrichment of glutamine synthetase, support metabolic remodeling across the E2–C1 tumor and stromal interface.

### Potential SPINK1-associated metabolic complementarity in the E2–C1 niche

To integrate the metabolic features associated with the E2–C1 niche, we mapped transcriptome-inferred reaction activity, metabolic enzyme and transporter expression, and spatial metabolite-associated ion signals across epithelial and CAF states (**Figure 7A**). Integrated analysis of gene expression, predicted reaction fluxes and spatial metabolite abundance supported a candidate lactate–glutamine circuit connecting E2 tumor cells with C1 CAFs (**Figure 7A**). Within the epithelial compartment, E2 showed higher expression of several glycolytic genes, including PGK1, ENO1 and HK2, than E1, consistent with its HIF1A-associated SPINK1 program (**Figure S3B**). The inferred reaction network connected phosphoenolpyruvate conversion to pyruvate with lactate formation, suggesting that E2 glycolysis could generate a carbon source accessible to neighboring CAFs. The preferential adjacency of E2 and C1 provided a spatial basis for this potential transfer of lactate from tumor cells to the stromal compartment (**Figures 4D** and **4E**). C1 CAFs exhibited relatively higher LDHB expression than the other CAF states, whereas LDHA expression was broadly comparable. Together with SLC16A1 expression, this pattern was compatible with lactate uptake and conversion to pyruvate. The accompanying oxidative profile included relatively prominent SDHB, FH and MDH2 expression, particularly compared with C3, as well as higher SDHA and SUCLG1/2 expression than C3. These enzymes participate in succinate formation and its subsequent oxidation through fumarate and malate to oxaloacetate, supporting the capacity to sustain downstream TCA metabolism. Spatial metabolomics localized elevated succinate signals to portions of the tissue margins and internal interfaces. Among the metabolic territories, GMM1 and GMM5 showed higher succinate intensity distributions, whereas GMM2 showed a lower distribution, indicating heterogeneous regional metabolite pools rather than enrichment confined to a single CAF-associated territory (**Figure 7A**). Coupling lactate utilization to TCA metabolism could also provide α-ketoglutarate (α-KG) for glutamate and glutamine synthesis. GOT1 expression was concentrated in C1, and GDH expression was more evident in C1 than C3, suggesting connections between TCA intermediates and glutamate production. Glutamine synthetase expression was maintained across C1–C3 and lower in C4, supporting glutamine biosynthetic potential within this stromal compartment. C1-derived glutamine could subsequently provide a substrate for E2 cells, where SLC1A5 and SLC38A5 expression suggested glutamine transport capacity (**Figures S12F–S12I**). Inferred reactions converting glutamine to glutamate and subsequently to α-KG thus connected potential stromal glutamine supply with the replenishment of tumor carbon intermediates. E2 cells expressed ACO2, IDH1/2 and ACLY, linking citrate metabolism to acetyl-CoA production, while showing lower SUCLG2 and SDHB expression than E1/E3 and lower FH expression than E1. These selective differences in downstream TCA enzymes were compatible with differential carbon allocation between oxidative metabolism and biosynthesis. IDH1/2-mediated reductive carboxylation of glutamine-derived α-KG to isocitrate and citrate, followed by ACLY-mediated citrate cleavage, therefore represents a candidate route for supplying acetyl-CoA to fatty acid synthesis. Within the proposed circuit, lactate utilization and glutamine synthesis in C1 could consequently support glutamine catabolism and lipogenesis in E2. The lipogenic component of this model was consistent with increased SREBP1 following rSPINK1 treatment (**Figures S12B–S12E**).

**Figure 7.**
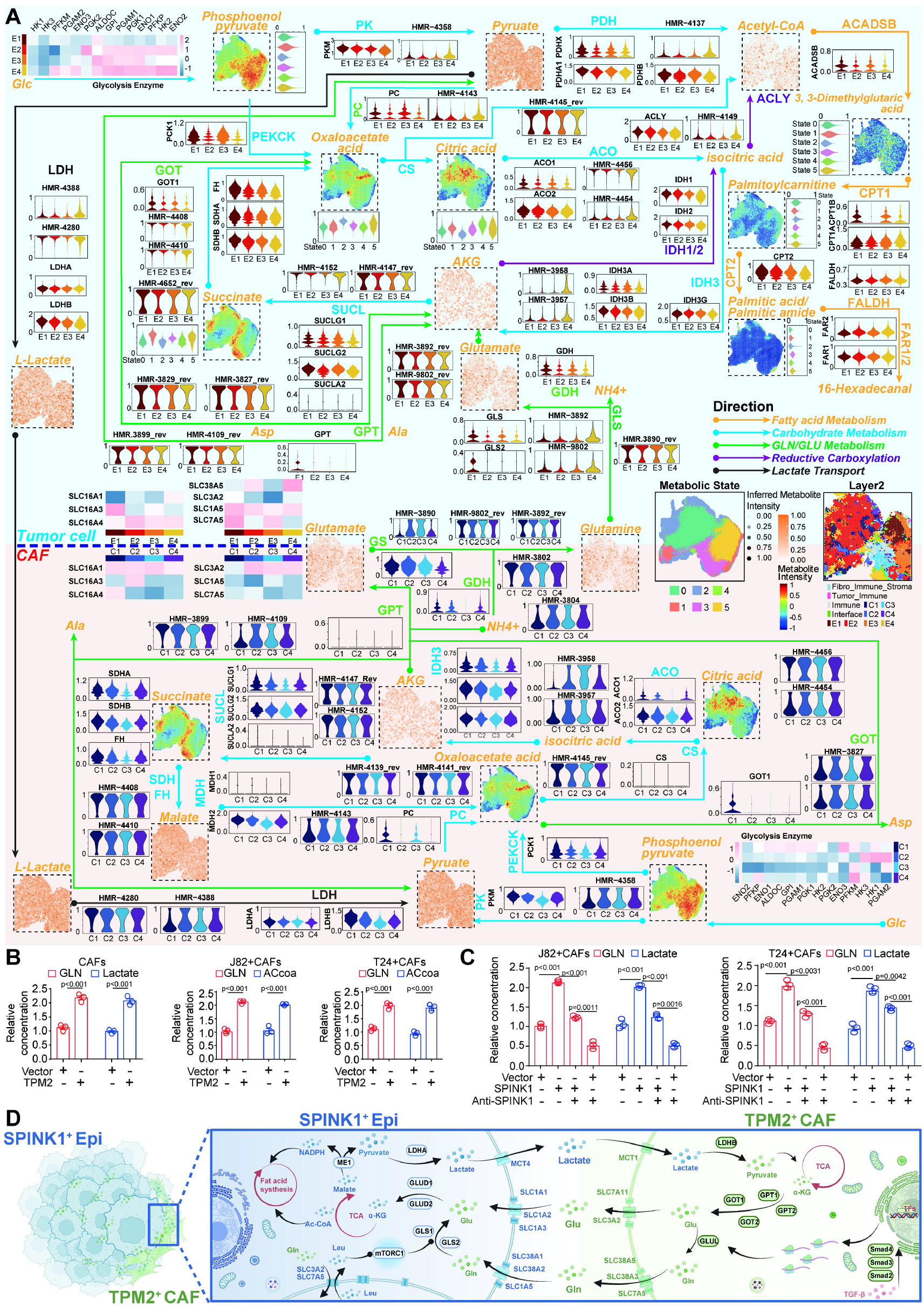
Metabolic programs and metabolite abundance in SPINK1⁺ tumor cells and TPM2⁺ CAFs. (A) Integrated map of glucose, glutamine/glutamate, TCA cycle, lactate, and fatty acid metabolism across epithelial states E1–E4 (top) and CAF states C1–C4 (bottom), combining HMR reaction scores inferred from transcriptomes, metabolic enzyme and transporter expression, and spatial metabolite abundance. (B) Relative intracellular glutamine and lactate levels in control and TPM2-overexpressing CAFs (left). Relative intracellular glutamine and acetyl-CoA levels in J82 cells (middle) and T24 cells (right) following coculture with control or TPM2-overexpressing CAFs. (C) Extracellular GLN and lactate levels in J82–CAF and T24–CAF cocultures under four conditions: vector control, SPINK1, SPINK1 plus neutralizing antibody (anti-SPINK1), and vector plus anti-SPINK1. (D) Working model integrating glycolytic, lipid, and glutamine metabolic programs in SPINK1⁺ tumor cells with oxidative and amino acid metabolic programs in TPM2⁺ CAFs. Metabolic enzymes and transporters are shown within their respective compartments; arrows between compartments indicate hypothesized routes of metabolite exchange.

To assess the metabolic effects of SPINK1 or TPM2 perturbation, we quantified changes in metabolite abundance. *TPM2* overexpression in patient derived CAFs increased relative glutamine and lactate levels compared with control CAFs (**Figure 7B**). In J82–CAF and T24–CAF co-cultures, *TPM2* overexpression increased glutamine and acetyl-CoA abundance relative to vector controls (**Figure 7B**). Culture supernatants from both co-culture systems further showed increased glutamine and lactate abundance following *SPINK1* overexpression, whereas SPINK1 neutralizing antibody attenuated these changes (**Figure 7C**). Overexpression and neutralization experiments provide mutually reinforcing evidence for SPINK1 responsive remodeling of metabolite availability in tumor–CAF cultures.

Integration of these perturbation data with the spatial and transcriptomic analyses supports a model in which the E2–C1 niche comprises metabolically distinct tumor and stromal compartments whose metabolic states are responsive to SPINK1-associated signaling (**Figure 7D**). C1 CAFs showed transcriptome-inferred programs related to lactate processing, oxidative metabolism, glutamine metabolism, and amino acid handling alongside TGF-β–SMAD signaling identified in the preceding signaling analyses. Distinct SLC family transporter profiles suggest potential routes for coordinating lactate and amino acid availability across the interface. SPINK1 perturbation altered extracellular glutamine and lactate abundance, whereas TPM2 overexpression in CAFs increased glutamine and acetyl-CoA abundance in tumor–CAF cocultures. Combined with the spatial enzyme and transporter profiles, these responses support metabolic remodeling at the E2–C1 tumor stromal interface involving both extracellular nutrient availability and biosynthetic metabolite pools. SPINK1 associated tumor stromal signaling contributes to metabolic remodeling within the E2–C1 niche.

### *Spink1* loss remodels the immune microenvironment and enhances tumor control during PD-1 blockade

Stromal organization can restrict antitumor immunity in urothelial cancer, with fibroblast TGFβ signaling and stromal transcriptional programs associated with limited T cell penetration and reduced responses to checkpoint blockade [9, 12]. The metabolic interactions proposed for the E2–C1 niche therefore raised the possibility that this niche also shapes the functional environment encountered by infiltrating T cells. CytoSig analysis identified positive inferred responses to TGFβ1, TGFβ3, IL10 and CXCL12 in E2 regions, accompanied by a weak IFNG response, whereas C1 regions exhibited an elevated IL6 response relative to the tissue reference (**Figures S14A–S14D**). These findings suggested a local environment conducive to fibroblast recruitment, fibrogenesis and immune regulation. TCellSI analysis identified regional cytotoxicity– and exhaustion-associated signature scores in mixed-cell Visium spots across E2– and CAF-enriched territories, with relatively higher exhaustion scores in C1 than in C2 and C4 regions (**Figures S14E** and **S14F**). Regional module scoring further characterized the metabolic and immune features of the E2–C1 niche. Single-cell ligand–receptor analysis identified stronger predicted T-cell-recruiting chemokine programs in C2–C4 than in SPINK1-associated C1 CAFs. Spatial transcriptomics further showed that C1-enriched regions preferentially expressed chemokines associated with myeloid and neutrophil recruitment, whereas C2–C4 regions retained T-cell-trafficking programs (**Figures 8A** and **8B**). Compared with other CAF-associated regions, C1 regions exhibited prominent branched chain amino acid, serine and glycine, glutamine metabolism and amino acid transport programs, accompanied by higher immune checkpoint scores and lower CD8 effector marker scores (**Figure 8C**). Across E2 tumor territories and adjacent C1 stroma, spatial mapping identified glycolysis and TGFβ programs alongside negative regulation of T cell activation, proliferation and effector processes. PD-1 coinhibition and PD-L1/PD-1 pathway signatures further connected this metabolic and stromal environment to local checkpoint regulation (**Figures S14G– S14J**). C1 regions also contained relatively high inferred abundances of SPP1⁺GPNMB⁺ macrophages, IL1R2⁺ neutrophils and TNFRSF9⁺ regulatory T cells (**Figures S14K–S14L**). Effector CD8⁺ T cells showed heterogeneous overlap with C1 and E2 territories (**Figures S14M–S14N**). These observations suggest that C2/C3-associated chemokine programs may facilitate lymphocyte recruitment, whereas the checkpoint-enriched C1 compartment and the exhaustion-associated E2 compartment may constrain effector function after recruitment.

**Figure 8.**
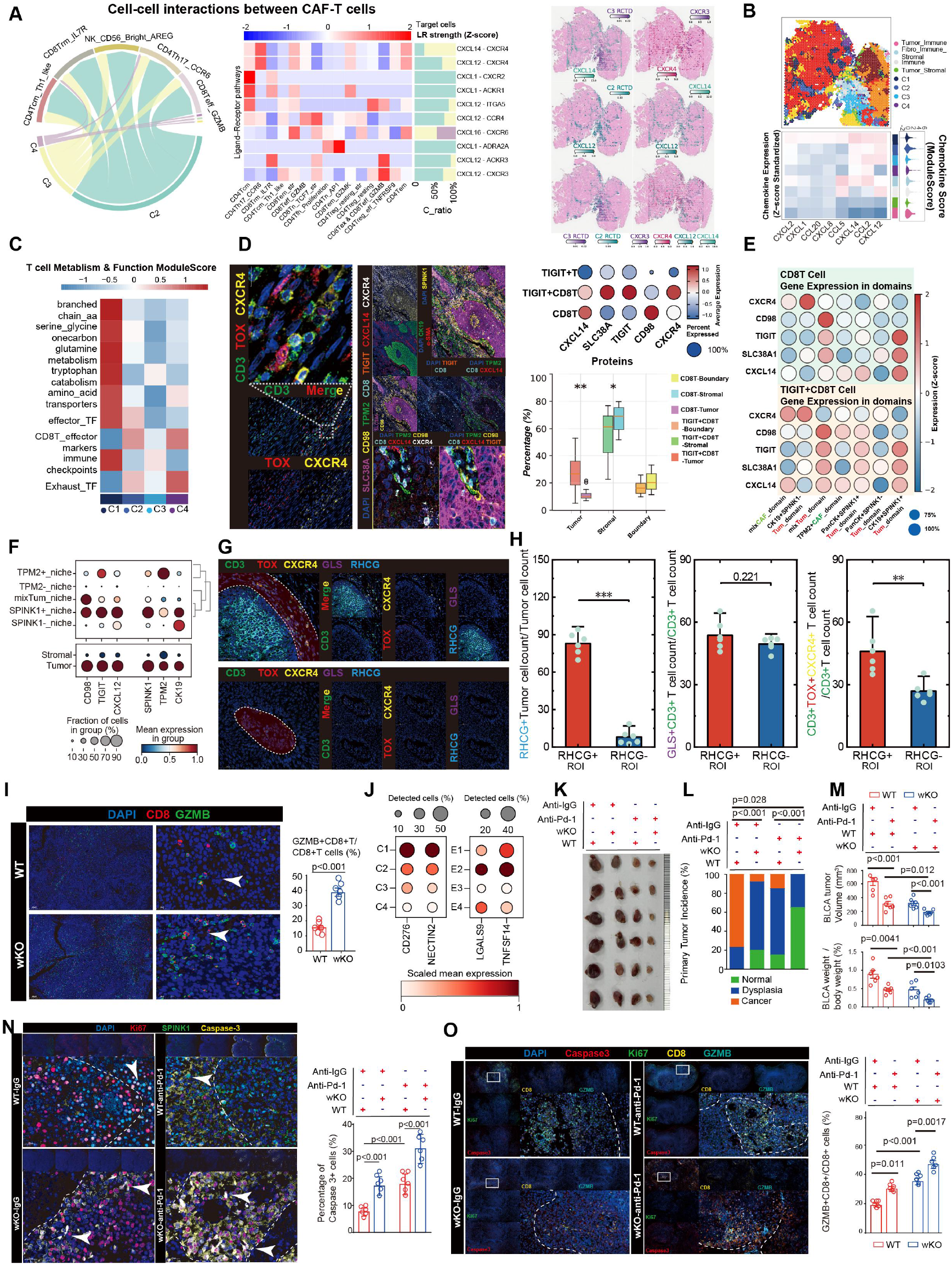
Spatial immune profiling and tumor control following *Spink1* loss and PD-1 blockade in BLCA. (A) Predicted CAF–T cell interactions and ligand–receptor scores shown by a chord diagram and heatmap, with spatial expression of CXCL14, CXCL12, CXCR3, and CXCR4. (B) Spatial domains and chemokine expression profiles in BLCA tissue. (C) T cell functional module scores across C1–C4 CAF and E1–E4 tumor spatial layers. (D) Multiplex immunofluorescence (mIF) images showing CD3, TOX, CXCR4, and CXCL14, with T cell phenotyping and distributions across tumor, stromal, and boundary regions. (E) Protein abundance and positivity in CD8⁺ and TIGIT⁺CD8⁺ T cells across spatial domains. (F) Protein profiles across tissue microarray (TMA) niches and tumor and stromal regions. (G) Representative mIF images showing spatial relationships among RHCG, CD3, TOX, CXCR4, and GLS. (H) RHCG⁺ tumor cell fractions and proportions of GLS⁺ and TOX⁺CXCR4⁺ cells among CD3⁺ T cells in RHCG⁺ and RHCG⁻ regions. (I) CD8 and Gzmb staining and Gzmb⁺CD8⁺ fractions among CD8⁺ T cells in *WT* and whole body *Spink1-wKO* mice following BBN exposure. (J) Expression of CD276 and NECTIN2 across CAF states C1–C4 and of LGALS9 and TNFSF14 across epithelial states E1–E4. Dot size indicates the percentage of cells with detected expression, and color denotes scaled mean expression. (K) Representative bladders from *WT* and *Spink1-wKO* mice receiving control IgG or anti-PD-1. (L) Distributions of normal mucosa, dysplasia, and carcinoma across treatment groups. (M) Quantification of tumor volumes and tumor weight/body weight ratios. (N) Ki67, SPINK1, and Caspase-3 staining, with quantification of Caspase-3⁺ cells. (O) Representative mIF images of Caspase-3, Ki67, CD8, and Gzmb, with Gzmb⁺CD8⁺ fractions among CD8⁺ T cells across treatment groups.

Multiplex imaging linked this spatial organization to T cell phenotypes. TOX and CXCR4 were co-expressed in stromal CD3⁺ T cells, and highly multiplexed imaging identified CXCL14-rich interfaces containing SPINK1⁺ tumor cells and TPM2⁺ CAFs. CXCR4⁺CD8⁺ T cells occupied tumor and interface regions containing SLC38A1 and CD98 signals, placing these cells within the amino acid transport landscape associated with the niche (**Figure 8D**). Compared with the overall CD8⁺ population, TIGIT⁺CD8⁺ T cells showed stronger CXCR4 and SLC38A1 signals and greater proportional representation in the tumor compartment, whereas CXCL14 signals were higher in the overall CD8⁺ population (**Figure 8D**). CXCR4 and CXCL14 profiles also varied between mixed CAF and TPM2⁺ CAF domains (**Figure 8E**). In the expanded TMA cohort, TIGIT signals were prominent in SPINK1⁺ tumor and TPM2⁺ niches, accompanied by regional variation in CXCL12 and CD98 (**Figure 8F**). These findings associated chemokine-related T cell localization with an exhaustion-associated marker phenotype in the tumor–stromal niche; recruitment dynamics and functional impairment were not directly measured.

The proposed glutamine exchange between C1 CAFs and E2 tumor cells provided a potential metabolic link to this immune phenotype. Glutamine catabolism generates ammonia, which can promote T cell exhaustion and induce effector CD8⁺ T cell death through lysosomal and mitochondrial damage [7, 13]. Spatial transcriptomics identified elevated expression of the ammonia transport protein RHCG in E2 related regions (**Figure S14O**). Multiplex imaging showed a greater proportion of TOX ⁺ CXCR4 ⁺ cells among CD3⁺ T cells in RHCG-positive tumor regions, whereas the GLS-positive T cell fraction did not differ significantly between RHCG-positive and negative regions (P = 0.221; **Figures 8G** and **8H**). These observations associated tumor ammonia transport machinery with local T cell dysfunction and nominated ammonia exposure as a potential consequence of E2–C1 metabolic interactions.

Checkpoint-associated molecules provided an additional potential constraint on recruited T cells. C1 CAFs showed prominent CD276 and NECTIN2 expression, whereas E2 tumor cells exhibited relatively high scaled expression of LGALS9 and TNFSF14 (**Figure 8J**). E2 cells also showed relatively high CD274 expression, encoding PD-L1, providing a molecular basis for potential PD-1 engagement within the tumor compartment (**Figure S14P**). Together with the spatial PD-1 pathway signatures, these findings support a putative T cell trap, C2/C3-associated chemokines may facilitate recruitment, while T cells entering the E2 – C1 niche encounter a PD-L1-expressing tumor compartment, checkpoint-associated stromal programs (**Figure S14Q**) and potentially adverse ammonia exposure. This model links lymphocyte recruitment to subsequent functional restriction through both checkpoint signaling and metabolic stress.

In BBN-induced bladder tumors, *Spink1* loss was accompanied by increased CD8⁺ T cell infiltration and a higher GZMB ⁺ fraction among intratumoral CD8 ⁺ T cells (**Figures S4F–S4I** and **8I**). Combining *Spink1* loss with PD-1 blockade produced the lowest tumor volume and tumor weight relative to body weight, together with less advanced histological lesions (**Figures 8K–8M, S14R**). Relative to the respective IgG controls, PD-1 blockade reduced mean endpoint tumor weight by 46.7% in *WT* mice and 61.2% in *wKO* mice (**Figures S14S**). The prevalence of dysplasia or cancer decreased from 80% to 35% in *wKO* mice (P = 0.019), compared with 100% to 85% in *WT* mice (**Figure S14T**). No carcinoma was observed in the combined intervention group, in which 65% of mice showed histologically normal tissue (**Figure S14U**).

Improved tumor control was accompanied by changes in both tumor cell markers and the intratumoral CD8⁺ T cell phenotype. Combined Spink1 loss and PD-1 blockade reduced Ki67 staining and produced the highest proportion of Caspase-3⁺ tumor cells, exceeding that observed with PD-1 blockade in WT mice (P < 0.001; **Figure 8N**). PD-1 blockade also further increased the GZMB⁺ fraction among intratumoral CD8⁺ T cells in *wKO* mice (P = 0.0017), with the combined group exhibiting the highest fraction across all four conditions (**Figure 8O**). Thus, the reduction in tumor burden coincided with lower proliferation-associated staining, increased Caspase-3 positivity and a more pronounced cytotoxic T cell phenotype. These findings support SPINK1 as a candidate cotarget for improving antitumor immunity and tumor control during PD-1 blockade.

## Discussion

Our study identifies a spatially organized BLCA niche in which SPINK1-high E2 tumor cells and TPM2-high C1 CAFs exhibit complementary metabolic programs and are associated with local immune dysfunction. These findings support a model in which nutrient cooperation between tumor cells and fibroblasts may sustain tumor anabolism while creating conditions that restrict recruited T cells. Molecular classifications have established substantial biological heterogeneity in BLCA [14]; the E2–C1 niche provides a tissue context for connecting this heterogeneity to metabolic organization and immune function.

Previous studies have described cellular neighborhoods and transcriptionally defined ecotypes [15, 16], while spatial analyses have linked CAF states to their surrounding communities [17]. A common integration strategy maps predefined single-cell identities onto tissue, potentially leaving functional variation associated with location unresolved within existing annotations [18]. Our workflow instead selects informative genes from spatial transcriptomes and combines MIA with NMF to define single-cell epithelial and stromal states through shared spatial gene programs. Spatial context therefore becomes a criterion for defining cellular states. This perspective also informs the clinical observations: bulk-derived E2/C1 fractions did not consistently stratify TCGA survival, whereas AI pathology predictions from TCGA H&E images identified significant survival differences. These findings suggest that tissue morphology contains clinically relevant information incompletely captured by aggregate cell fractions.

SPINK1 and TPM2 perturbation connects this spatial organization to functional interactions. SPINK1 has been implicated in EGFR-associated growth signaling in prostate cancer [19]. Here, SPINK1 depletion restricted tumor growth, whereas recombinant SPINK1 promoted proliferative signaling and induced myofibroblastic programs in CAFs, supporting potential autocrine and paracrine functions. TPM2 overexpression promoted CAF proliferation and motility and increased SMAD2/3 phosphorylation. Together with predicted SPINK1–EGFR and TGF-β-associated communication, these observations suggest that local signaling may coordinate epithelial and stromal behavior, consistent with extracellular regulation of CAF states through mechanisms such as IL1–TGF-β antagonism in pancreatic cancer [20].

Such regulation could also shape nutrient demand and availability. Stromal alanine secretion and glutamine provision support tumor metabolism in other cancers [5, 21]. For potentially exchangeable metabolites, local abundance reflects production, uptake and redistribution across cellular boundaries. Spatial metabolomics was therefore interpreted through both regional intensity and the gradients connecting abundance peaks and troughs. Intensity-derived fields and Helmholtz decomposition characterized divergent and rotational structure across territories and interfaces, describing potential exchange environments rather than directly measuring transport [22]. Putative phosphoinositide features, more closely associated with cellular membranes, complemented the PI3K/AKT transcriptional program and linked the metabolic landscape to the signaling context discussed above. This approach provides a spatial basis for interpreting metabolic relationships between neighboring populations.

The integrated data support a model of metabolic symbiosis linking E2 glycolysis and lipogenesis to C1 oxidative metabolism. In this model, E2-derived lactate supplies C1 with pyruvate for TCA metabolism, connecting α-ketoglutarate (αKG) generation to glutamate and glutamine synthesis. Glutamine provision to E2 replenishes glutamate and αKG, which could support reductive carboxylation to citrate and ACLY-dependent acetyl-CoA production for fatty acid synthesis [23]. E2 IDH1/2 and ACLY profiles, together with relatively prominent citrate/αKG-associated features and less uniform enrichment of subsequent TCA steps, support this interpretation; spatial succinate-associated signals localized to CAF-associated margins. The HIF1A-associated SPINK1 program and SREBP1 response to recombinant SPINK1 further connect this state to glycolytic and lipogenic regulation. The pericyte-like component of C1 suggests a possible perivascular association. If C1 occupies relatively oxygenated sites, lactate oxidation could reduce glucose demand and competition with E2, potentially involving feedback on glycolysis. Such glucose sparing has precedent in tumor cells [24]. Close E2–C1 apposition could facilitate cooperation by shortening diffusion distances.

Glutamine support for tumor anabolism also raises the possibility that its nitrogenous products influence local immunity. Lactate and glutamine metabolism affect lymphocyte function [25, 26], and fibroblast TGF-β programs have been linked to immune restriction in urothelial cancer [9]. C2/C3-associated chemokines may recruit T cells into an E2–C1 environment containing metabolic and checkpoint constraints. E2 glutamine catabolism could increase ammonia generation, with RHCG representing a candidate route for release into surrounding tissue. The association of RHCG-positive regions with TOX⁺CXCR4⁺ T cells nominates ammonia handling as a potential connection to immune dysfunction. Studies linking microenvironmental ammonia to exhaustion and intracellular ammonia accumulation to effector T cell injury provide mechanistic precedent [7, 13]. Together with E2 PD-L1, regional PD-1 signatures and C1 CD276/NECTIN2 expression, these observations support a putative T cell trap in which recruitment may coexist with subsequent functional restriction.

The association of fibroblast-rich, immune-excluded communities with nonresponding bladder tumors further motivates intervention in this tissue context [27]. *Spink1* loss increased CD8⁺ infiltration and the GZMB⁺ fraction among intratumoral CD8⁺ T cells, while combination with PD-1 blockade produced the lowest tumor burden, reduced Ki67 staining and increased the Caspase-3⁺ tumor cell fraction. These observations link improved tumor control to changes in malignant and immune compartments and support SPINK1 as a candidate cotarget for immunotherapy.

## Limitations

Several questions remain unresolved, including niche formation, perivascular oxygenation, directional metabolite transfer and the contribution of ammonia to T cell dysfunction. Whole-body deletion combines tumor and host effects, and reduced tumor burden may itself alter immunity. Matched prognostic validation, compartment-specific perturbation, metabolic tracing and pharmacological intervention in established tumors are needed. The E2–C1 niche provides a testable model linking local metabolic cooperation to restricted antitumor immunity, with SPINK1 as a candidate intervention point.

## RESOURCE AVAILABILITY

### Lead contact

Further information and requests for resources and reagents should be directed to, and will be fulfilled by, the lead contact, Lufeng Zheng.

### Data and code availability

Publicly available single-cell RNA sequencing datasets reanalyzed in this study are available from the Gene Expression Omnibus (GEO) under accession numbers GSE129845, GSE186520, GSE190888, GSE217956, GSE222315, GSE267718, GSE269877, and GSE277524. Spatial transcriptomic data are available under GEO accession number GSE285715. TCGA-BLCA transcriptomic data, clinical annotations, and histopathology images are accessible through the Genomic Data Commons Data Portal. Whole-exome sequencing data from the source cohort are deposited in dbGaP under accession number phs003742.v1.p1, with access subject to authorization and the applicable data-use agreements. Additional bulk transcriptomic validation datasets are available from the GEO under accession numbers GSE133624, GSE188715, GSE59483, GSE120736, GSE37815, GSE69795, and GSE70691. RNA sequencing, spatial metabolomics, and imaging data generated in this study will be made available through the GEO under accession numbers GSE285715 or DOIs 10.1073/pnas.2505125123 and 10.1038/s41417-025-00947-z [28, 29].

Custom analysis code developed for this study will be made publicly available at github.com/zlab-cpu/BLCA upon publication.

Any additional information required to reanalyze the data reported in this paper is available from the lead contact upon request.

## Materials availability

### STAR METHODS

#### Human BLCA cohorts and clinical specimens

This study assembled complementary human BLCA cohorts encompassing multiple molecular modalities and clinical contexts. The integrated single-cell RNA-sequencing cohort comprised 82 tissue samples, including 66 bladder tumors and 16 adjacent tissues. Available clinical annotations included tumor grade for 25 cases, pathological T stage for 21 cases, 23 primary tumors, and 17 recurrent tumors. Samples were also obtained across different treatment settings, including BCG-naive (n = 14), BCG-treated (n = 14), epirubicin-naive (n = 4), and epirubicin-treated (n = 4) groups. Matched whole-exome sequencing data from paired tumor and peripheral blood mononuclear cell specimens were available for 23 patients. Spatial transcriptomic and spatial metabolomic profiling was performed on three tumor and three adjacent tissue sections; the spatial transcriptomic specimens represented paired tumor and adjacent tissues from three patients. Multiplex immunofluorescence-based spatial proteomic profiling was conducted on 14 human BLCA specimens using multiple antibody panels. An independent tissue microarray cohort comprising 96 tissue samples from 63 patients was used for extended multiplex immunofluorescence validation. Clinical and pathological annotations were used, where available, for comparisons according to tissue type, tumor grade, pathological stage, primary versus recurrent disease, and treatment status.

#### Human bladder tissue collection and ethics statement

All animal experiments described in this study were reviewed and approved by the Animal Care and Use Committee of Fudan University (Approval No.: 202206033S). Following approval by the Human Research Ethics Committee of Huashan Hospital, Fudan University (KY2011-009), paired samples of BLCA and adjacent tissues were obtained from patients who had provided written informed consent. The clinicopathological information of the included patients was described in **Supplementary Table S1** [28, 29]. Human BLCA tissues arrays, which contains 96 samples from 63 patients, were purchased from Shanghai OUTDO Biotech Co., Ltd (Cat #JS W-11-01).

#### Clinical metadata collection and cohort harmonization

Clinical metadata were compiled from the annotations provided for each cohort and harmonized before cross-cohort analysis. Tissue origin was categorized as tumor or adjacent; tumor grade as high or low; disease status as primary or recurrent; and treatment exposure as BCG-naive or BCG-treated and epirubicin-naive or epirubicin-treated. Pathological T stage was retained according to the reported T-category designation. Analyses involving individual clinical variables were restricted to samples with the corresponding annotations.

#### Cell culture, lentivirus infection, and reagents

Mouse bladder carcinoma cell line MB49 was obtained from Procell (Wuhan, China). Human BLCA cell lines (T24, J82) were obtained from ATCC. These cells were maintained in RPMI1640 medium (Thermo Fisher Scientific, Waltham, MA, USA) supplemented with 10% fetal bovine serum (FBS; Cat #OM780027, OmnimAbs, Shanghai, China), 1% penicillin, and 1% streptomycin (Cat #E607064, Sangon Biotech, Shanghai, China). Primary CAFs were isolated from freshly resected BLCA specimens obtained from patients undergoing surgical resection at Huashan Hospital, Fudan University (KY2011-009). Briefly, tissue samples were collected in sterile RPMI 1640 medium (Cat #11875085, Thermo Fisher Scientific) supplemented with 1% penicillin-streptomycin and transported to the laboratory on ice within 30 min of resection. Tumour tissues were washed three times with ice-cold phosphate-buffered saline (PBS) containing 1% penicillin-streptomycin to remove blood and debris. Specimens were then minced into small pieces of approximately 1-2 mm³ using sterile scalpels and subjected to enzymatic digestion in DMEM/F12 medium (Cat #12634010, Thermo Fisher Scientific) containing 1 mg/mL collagenase IV (Cat #C4-BIOC, Sigma-Aldrich) and 0.1% DNase I (Cat #10104159001, Sigma-Aldrich) at 37°C for 1–2 h with intermittent agitation. The resulting cell suspension was filtered through a 100 µm cell strainer to remove undigested tissue fragments and centrifuged at 300 × g for 5 min at room temperature. The cell pellet was resuspended in DMEM/F12 medium supplemented with 10% FBS and 1% penicillin-streptomycin, and plated in T25 culture flasks. After 2–4 h of incubation at 37°C in a humidified atmosphere containing 5% CO₂, non-adherent cells, including epithelial tumour cells and immune cells, were removed by gentle washing with PBS. The remaining adherent fibroblasts were maintained in complete DMEM/F12 medium with 10% FBS and 1% penicillin-streptomycin at 37°C and 5% CO₂, with medium changes every 2–3 days until reaching 80–90% confluence. To obtain purified CAF populations, cells were subjected to differential trypsinization (0.25% trypsin-EDTA, Cat #25200072, Thermo Fisher Scientific) for 1–2 min, allowing preferential detachment of contaminating epithelial cells, followed by repeated passaging. CAF identity was confirmed by positive immunofluorescence staining for fibroblast activation protein (FAP), vimentin, and α-smooth muscle actin (α-SMA), and by negative staining for pan-cytokeratin (Pan-CK) and CD45. All CAF lines were used between passages 3 and 6 for subsequent functional assays to minimize phenotypic drift. Cell culture media were supplemented with 1% non-essential amino acids (Cat #11130051) and 1 mM sodium pyruvate (Cat #11360070) (both from Thermo Fisher Scientific) where indicated for specific metabolic assays. For SPINK1 modulation, the construction and packaging of all plasmids were carried out by OBiO (Shanghai, China). Lentiviral vectors encoding SPINK1 shRNA (shSPINK1) or overexpression constructs (SPINK1-OE) were added into cells using Polybrene (8 μg/mL, Sigma). The efficiency of infection was validated by western blotting. Recombinant SPINK1 protein (Cat #HY-P71053) was purchased from MedChemExpress (Monmouth Junction, NJ, USA). SPINK1 neutralizing antibody (clone 839304) (MAB74961) was purchased from R&D Systems and used at a concentration of 5 μg/mL.

#### Mouse strains and animal experiments

All experimental animals were maintained under specific pathogen-free (SPF) conditions with controlled 12-h light/dark cycles. *Wild-type (WT)* C57BL/6 (Strain ID: N000013) and *Spink1-knockout (KO)* mice (Strain ID: T044045) were commercially acquired from GemPharmatech Co., Ltd (Nanjing, China).

Carcinogenesis Protocol: Carcinogen-induced BLCA was initiated in male mice (≥5 weeks old) through continuous administration of 0.05% N-butyl-N-(4-hydroxybutyl) nitrosamine (BBN) in drinking water over 16 weeks, as previously established [30].

Immunotherapy Intervention: Following BBN exposure, wild-type (WT) and *Spink1*-*knockout (KO)* mice were divided into four groups: WT + IgG, WT + anti-PD-1, KO + IgG, and KO + anti-PD-1. Mice received intraperitoneal injections of either anti-PD-1 monoclonal antibody (200 μg/dose; clone RMP1-14, Bio X Cell, Cat# BE0146) or isotype control IgG (clone 2A3, Bio X Cell, Cat# BE0089) three times weekly.

Orthotopic BLCA Model: MB49 murine bladder carcinoma cells were genetically engineered to express firefly luciferase through lentiviral transduction (puromycin selection, 2 µg/mL for 72 h). For orthotopic implantation, female C57BL/6 mice underwent 8-h fasting before transurethral instillation of 8×10^^5^ luciferase-expressing MB49 cells suspended in 100 µL PBS.

Surgical Procedure: Anesthetized mice received bladder preconditioning with 50 µL poly-L-lysine (20-min dwell time) prior to tumor cell inoculation. Urethral catheterization was performed under aseptic conditions using lubricated soft catheters, with bladder evacuation achieved through manual compression.

Tumor Monitoring & Analysis: Quantitative tumor monitoring was performed using bioluminescence imaging (IVIS Spectrum, PerkinElmer) following intraperitoneal luciferin injection (150 mg/kg). Terminal measurements at day 14 included tumor volume (calculated as length×width²/2) and bladder index (tumor-to-body weight ratio). All specimens were harvested for histopathological evaluation post-euthanasia.

#### Tumor cell & CAF co-culture system

J82 and T24 BLCA cells were treated with recombinant human SPINK1 protein (rhSPINK1, 100 ng/mL, Cat #ab152041, Abcam) or anti-SPINK1 neutralizing antibody (clone 839304) (MAB74961) for 24 h. Culture medium was then removed and cells were washed twice with PBS. Treated cells (2×10⁴ cells/well) were seeded in serum-free medium in the upper chamber of Transwell inserts (Cat #14341, LABSELECT, Beijing, China). Patient-derived CAFs (5×10⁴ cells/well) were seeded in complete medium in the lower chamber. After 48 h of co-culture, culture supernatants from the lower chamber were collected and centrifuged at 1,000 g for 5 min to remove cell debris. Extracellular glutamine and lactate concentrations in the culture medium were measured using the Glutamine Assay Kit (Abcam, Cat #ab197011) and Lactate Assay Kit (Sigma-Aldrich, Cat #MAK064) according to the manufacturer’s instructions. Absorbance was measured at 450 nm (glutamine) and 570 nm (lactate) using a microplate reader (BioTek Synergy H1).

#### Cell viability assay and Transwell co-culture

Cell viability assays were performed to evaluate the effects of SPINK1 and TPM2 on BLCA cells and primary CAFs, respectively. Briefly, BLCA cells with SPINK1 overexpression (SPINK1-OE) or knockdown (SPINK1-KD), together with their corresponding controls, were cultured independently to assess the effects of SPINK1 modulation on BLCA cell viability. Similarly, primary CAFs overexpressing TPM2 (TPM2-OE) or the corresponding vector control were cultured independently to determine the effects of TPM2 on CAF viability. To further investigate reciprocal tumor–stromal interactions, BLCA cells and primary CAFs were co-cultured using a Transwell coculture system. To evaluate the effects of TPM2 overexpression in CAFs on BLCA cell viability, TPM2-overexpressing CAFs or vector-control CAFs were seeded in the upper inserts, while BLCA cells were cultured in the lower chambers. Conversely, to assess the effects of tumor-derived SPINK1 on CAF viability, SPINK1-overexpressing BLCA cells or their corresponding vector controls were seeded in the upper inserts, with primary CAFs cultured in the lower chambers.

Following the indicated culture or co-culture periods, cell viability was measured using CCK-8 colorimetric assay (Cat #C6005, NCM Biotech, China) following standardized protocols. For co-culture experiments, viability was assessed specifically in the recipient cells cultured in the lower chambers. Briefly, 5×10³ cells/well were plated in 96-well microplates and maintained under standard culture conditions. At 24, 48, and 72 h, cellular metabolic activity was determined by incubating cultures with 10 μL CCK-8 reagent diluted in 100 μL PBS during the terminal 2-h period. Absorbance values at 450 nm were recorded using a BioTek Synergy H1 multimode microplate reader (Winooski, VT, USA), with triplicate biological replicates performed for each experimental condition.

#### EdU incorporation assay

Cells were seeded in 24-well plates (5×104/well) and treated as indicated. After 48 h, 10 μM EdU (Cat #ST067, Beyotime, Beijing, China) was added for 2 h. Cells were fixed with 4% paraformaldehyde, permeabilized with 0.3% Triton X-100, and stained using the BeyoClick™ EdU Cell Proliferation Kit with AF488 (Cat #C0071S, Beyotime). Nuclei were counterstained with DAPI (Cat #D9542, Sigma-Aldrich). EdU-positive cells were quantified from five random fields per sample using ImageJ.

#### Wound-healing assay

When cell confluence reached 90% in 6-well plates, a wound was created on the cell surface, and cell debris was washed away with PBS twice. To reduce the contribution of proliferation to wound closure, cells were then cultured in a serum-starved medium (≤2%) containing the compounds or solvent. The migration rate was calculated as follows: Migration rate = (D_0_ – D_t_)/D_0_×100%, where D_t_ represents the distance between the wound edges at different time points, and D_0_ represents the initial wound distance at 0 h.

#### Transwell invasion assays

For invasion assays, transwell membranes were pre-coated with Matrigel (200 μg/mL, Cat # 356234, BD Biosciences) for 4 h. Cells (2×105) were seeded in serum-free medium. After 48 h, invaded cells were processed and quantified as above. Three independent experiments were performed in triplicate.

#### Western blotting

Total proteins were extracted using RIPA lysis buffer (Cat #P0013B, Beyotime) containing protease/phosphatase inhibitors (Cat #PPC1010, Sigma-Aldrich). Protein concentration was determined via BCA assay kit (Cat #P0009, Beyotime). Lysates (30 μg/lane) were separated on 10% SDS-PAGE gels and transferred to PVDF membranes (Cat #03010040001, Sigma-Aldrich). After blocking with 5% non-fat milk, membranes were incubated with antibodies against SPINK1 (1:1000, Abcam #ab183034), EGFR (1:1000, CST #4267), p-EGFR (Tyr1068, 1:1000, CST #3777), p-PI3K (1:1000, CST #17366), PI3K (1:1000, CST #4257), AKT (1:1000, CST #9272), p-AKT (Ser473, 1:1000, CST #4060), or GAPDH (1:5000, CST #2118), TGF-β (1:1500, CST #3711), EGF (1:2000, Abcam #ab184265), p-Smad2 (1:1500, CST #3108), Smad2 (1:1500, CST #5339), p-Smad3 (1:1500, CST #9520), Smad3 (1:1500, CST #9513), p-mTOR (1:1500, CST #5536), mTOR (1:1500, CST #2983). HRP-conjugated secondary antibodies (1:5,000, CST) and ECL substrate (Cat # P0018JS, Beyotime) were used for detection. Band intensities were quantified using ImageJ.

#### Immunofluorescence staining

Formalin-fixed paraffin-embedded (FFPE) tissues were sectioned at 4 μm thickness. After deparaffinization and antigen retrieval (citrate buffer, pH 6.0), sections were blocked with 5% BSA and incubated overnight with primary antibodies: anti-SPINK1 (1:200, Abcam #ab231208), anti-TPM2 (1:100, Proteintech #66325-1-Ig), anti-α-SMA (1:200, CST #19245), or anti-Ki67 (1:100, CST #9129). Alexa Fluor 488/594-conjugated secondary antibodies (1:500, Invitrogen) were applied for 1 hr. Nuclei were counterstained with DAPI (Sigma-Aldrich). Images were captured using a confocal microscope (Zeiss LSM 980) and analyzed with ZEN 3.4 software.

#### Intracellular metabolite quantification

Patient-derived CAFs were transfected with TPM2 overexpression plasmid (TPM2-OE) or empty vector control using Lipofectamine 3000 (Cat #L3000001, Thermo Fisher Scientific) according to the manufacturer’s instructions. After 48 h of transfection, cells were washed twice with ice-cold PBS and lysed in metabolite extraction buffer. Cell lysates were centrifuged at 12,000 g for 10 min at 4°C, and supernatants were collected for metabolite analysis. Intracellular glutamine and lactate levels were measured using the Glutamine Assay Kit (Abcam, Cat #ab197011) and Lactate Assay Kit (Sigma-Aldrich, Cat# MAK064). Absorbance was measured at 450 nm (glutamine) and 570 nm (lactate) using a microplate reader (BioTek Synergy H1). Additionally, patient-derived CAFs with or without TPM2 overexpression were co-cultured with BLCA cells. After 48 h of co-culture, glutamine and acetyl-CoA levels were measured in BLCA cells using Glutamine Assay Kit (Abcam, Cat #ab197011) and Acetyl-CoA Assay Kit (Sigma-Aldrich, Cat #MAK566). Metabolite levels were normalized to total protein concentration determined by BCA assay and expressed as fold change relative to vector control. Statistical significance was determined by two-tailed Student’s t-test.

#### Extracellular metabolite measurement

J82 and T24 BLCA cells with or without SPINK1 overexpression were treated with anti-SPINK1 neutralizing antibody (clone 839304) (MAB74961) for 24 h. Culture medium was then removed and cells were washed twice with PBS. Treated cells (2×10⁴ cells/well) were seeded in serum-free medium in the upper chamber of Transwell inserts (Cat #14341, LABSELECT). Patient-derived CAFs (5×10⁴ cells/well) were seeded in complete medium in the lower chamber. After 48 h of co-culture, culture supernatants from the lower chamber were collected and centrifuged at 1,000 g for 5 min to remove cell debris. Extracellular glutamine and lactate concentrations in the culture medium were measured using the Glutamine Assay Kit (Abcam, Cat# ab197011) and Lactate Assay Kit (Sigma-Aldrich, Cat# MAK064). Absorbance was measured at 450 nm (glutamine) and 570 nm (lactate) using a microplate reader (BioTek Synergy H1).

#### Bulk RNA sequencing

T24 BLCA cells and patient-derived CAFs were treated with recombinant human SPINK1 protein (rhSPINK1, 10 ng/mL, Cat #ab152041, Abcam) or vehicle control for 72 h. Total RNA was extracted using TRIzol reagent (Cat #15596026CN, Thermo Fisher Scientific) according to the manufacturer’s instructions. RNA quality and concentration were assessed using a NanoDrop spectrophotometer and Agilent 2100 Bioanalyzer. Samples with RNA integrity number (RIN) ≥ 7 were used for library preparation. RNA-seq libraries were constructed using the NEBNext Ultra RNA Library Prep Kit (New England Biolabs) and sequenced on an Illumina NovaSeq 6000 platform with 150 bp paired-end reads.

#### Human BLCA staging and histological grading

Human bladder tumour specimens were evaluated on haematoxylin-and-eosin-stained sections. Histological type, histological grade, and anatomical stage were recorded as separate pathological variables. For urothelial tumours, grade was classified as low grade or high grade according to the WHO/ISUP system represented in AJCC-7; urothelial carcinoma in situ was treated as a high-grade lesion [31, 32]. Anatomical extent was classified using the AJCC-7 TNM system [31]. The T category reflected the deepest histologically confirmed level of invasion. Regional lymph-node and distant-metastasis categories were derived from the available pathological, operative, and radiological records, with the clinical or pathological prefix retained as applicable.

For the prespecified patient-level analysis, the highest verified T category and highest histological grade were retained when a patient had multifocal or temporally distinct tumours; specimen-level findings remained available for sensitivity or descriptive analyses. The presence of muscularis propria in each specimen was documented, and pT2a or pT2b was assigned only when the depth of muscularis propria involvement could be assessed reliably. Cases with insufficient information were coded TX or NX, rather than being assigned to a lower category. Histological grade was not inferred from the AJCC stage group. The AJCC Seventh Edition anatomical stage grouping, TNM definitions, and histological grading criteria used for pathological annotation are summarized in **Tables S2–S4**, respectively.

#### Mouse urinary bladder histopathological assessment

At the prespecified experimental endpoints, mouse urinary bladders were collected, gently emptied, fixed under standardized partial inflation, consistently oriented, and paraffin embedded. Serial sections obtained at standardized anatomical levels were stained with haematoxylin and eosin. Slides were coded to conceal genotype and treatment allocation during assessment. Each lesion was first assigned an exact morphological diagnosis using terminology aligned with INHAND recommendations for the rodent urinary tract [33, 34]. Model-specific descriptors, including dysplasia-like atypical proliferation and carcinoma-in-situ-like flat high-grade atypia, were retained when needed to describe the observed spectrum [35–37].

The assessment recorded growth pattern, cytological atypia, focality, lesion extent, histological differentiation, and the deepest anatomical layer involved. A study-defined histopathological progression score was then assigned for animal-level analysis. The highest score across all adequate sections from an animal was used as the primary ordinal endpoint, while the exact lesion diagnoses were retained. Carcinoma incidence was calculated only from unequivocal carcinoma diagnoses. Each animal, rather than each section, microscopic field, or separate lesion, was the independent experimental unit. Reporting follow the applicable ARRIVE 2.0 items [38].

#### Statistical analysis of mouse pathology

Ordinal scores were reported as medians and interquartile ranges, together with the number and proportion of animals in every score category. Carcinoma incidence and muscle-invasive carcinoma incidence are also presented as n/N (%). For two independent groups, scores were compared using an exact Mann–Whitney U test. For three or more groups, a Kruskal–Wallis test with multiplicity-adjusted post hoc comparisons is appropriate. Proportional-odds ordinal regression may be used for covariate-adjusted analyses after checking the proportional-odds assumption. Incidences were compared using Fisher’s exact test. The analysis plan, missing-data handling, and multiplicity procedure was specified before unblinding.

#### Single-cell RNA-seq quality control and preprocessing

Publicly available scRNA-seq datasets generated over an extended period, including several early-generation datasets, were integrated. To accommodate differences in sequencing depth and data quality, cells with fewer than 200 total counts or with mitochondrial transcripts accounting for more than 35% of total counts were excluded. Genes detected in fewer than three cells were removed. Gene identifiers across datasets and sequencing batches were mapped to current gene symbols before integration. Expression matrices were subsequently merged using an outer join to preserve the union of genes across all datasets.

#### Single-cell RNA-seq normalization and batch integration

Following quality control, the merged count matrix was normalized for library size and log-transformed. Highly variable genes were identified using the variance-stabilizing procedure implemented in Seurat v3 [39]. The normalized expression matrix was subsequently centered and scaled to unit variance using the scale function, followed by principal-component analysis. To mitigate inter-sample technical variation, Harmony was applied to the principal-component embedding with sample identity specified as the batch covariate [40]. The Harmony-corrected components were used to construct a nearest-neighbor graph and generate a UMAP embedding. Unsupervised graph-based clustering was performed at a resolution of 0.6.

#### Single-cell clustering and cell-type annotation

The corresponding cell clusters were extracted, and then re-clustered using the unsupervised clustering. The functional gene signatures were inputted for “AddModuleScore” scoring method. Functional subtypes were then annotated based on the scoring of each feature.

#### Tumor cnv analysis

We employed inferCNV method [41] to complete the tumor malignancy evaluation. Here, we evaluated the malignancy of epithelial in tumor and normal samples using CNV score.

#### Cell-state perturbation analysis

scDist [42] provides a statistically rigorous and computationally efficient approach for detecting transcriptomic differences based on a mixed-effects model. In brief, scDist estimates the distance between condition means in high-dimensional gene expression space for each cell type to identify the most perturbated one.

#### Cell trajectory analysis

Using slingshot method [43] to identify the cellular trajectories of cell clusters. The slingshot was used to perform the pseudotime trajectory analysis and describe putative state relationships and genes associated with pseudotemporal ordering; these analyses do not establish lineage or temporal progression.

#### Non-negative matrix factorization of epithelial and CAF states

To classify the subtypes of cells, the NMF clustering method was applied, based on the integrated gene list of MIA common genes using package “NMF” [44]. The configuration was set as follows: the “snmf/r” method with default parameters, and different subgroups were obtained depending on the scRNA expression matrix.

#### Pathway activity and gene-set enrichment analysis

To identify the varied molecular pathways in samples, the R package Escape (ver 2.1.4) was used for pathway score computation. Gene sets were derived from the Hallmark library of the Molecular Signature Database and from previous publications.

To explore the activity of different NMF subclusters in various molecular pathways, the “progeny” package [45] was utilized to estimate the intensities of 11 different signaling pathways.

#### Transcription factor and regulon activity analysis

To identify the potential TF regulatory networks in different types of subclusters in scRNA data, we performed SCENIC (ver 1.3.1) analysis [46]. The GENIE3 was used for identification of TF targets via constructing co-expression network, then the enrichment of TF-motif and discovery of direct TF target were completed by RcisTarget. The reference genome dataset was acquired from cisTarget Database (https://resources.aertslab.org/cistarget/). Eventually, the AUCell score of regulon activity in each cell was calculated.

#### Metabolic pathway activity scoring

We used scMetabolism [47] to calculate transcriptome-based metabolic pathway activity scores from the single-cell expression matrix. The analysis included 85 KEGG and 82 Reactome pathways. Differential pathway activity between cell states was assessed using limma.

#### Single-cell metabolic flux inference

The different metabolic flux between the NMF subtypes were also investigated. We performed single cell flux estimation analysis (scFEA, ver 1.1.2) [48] to specifically explore the metabolites of NMF in single-cell level. scFEA generates a graph neural network based on inputted single-cell matrix, module gene file and the stoichiometry matrix of relationship between compounds and modules, then characterize the metabolic flux activities in different cell clusters. Using “limma” package, the differential expressed metabolic pathways between these groups were identified.

#### Bulk RNA-seq read processing and expression quantification

Raw sequencing reads were quality-filtered using FastQC and trimmed with Trimmomatic. Clean reads were aligned to the human reference genome (GRCh38) using HISAT2. Gene expression levels were quantified as fragments per kilobase of transcript per million mapped reads (FPKM) using StringTie.

#### Bulk RNA-seq deconvolution

To evaluate the cell composition of TCGA-BLCA bulk-seq samples, we used the BayesPrism (https://github.com/Danko-Lab/BayesPrism) package [49], which models a prior from cell-type-specific expression profiles from single-cell RNA-seq data to jointly estimate the posterior distribution of cell type composition and cell-type-specific gene expression from bulk RNA-seq expression data of tumor or non-tumor samples.

#### Whole-exome sequencing data processing and variant calling

Whole-exome sequencing data for 27 samples from 23 patients, including tumor and peripheral blood mononuclear cell specimens were obtained from dbGaP (accession phs003742.v1.p1) and subjected to secondary analysis. Sequencing and initial variant calling were completed in the source study; no additional library preparation or sequencing was conducted in the present study. The source data were generated using SureSelect V5 capture and 150 bp paired end sequencing, with a mean coverage of 187× and a range of 151× to 263×. Sequencing reads were aligned to the GRCh38 reference genome using STAR. Somatic single nucleotide variants were identified using Mutect2, SomaticSniper v1.0.4, VarScan v2.4.3, Strelka v2.9.10, and Platypus v0.8.1. Small insertions and deletions were detected using Mutect2, VarScan, Strelka, and Platypus. Variant annotation was performed using SnpEff.

Variants were retained when tumor coverage was greater than 10, tumor variant allele frequency was at least 0.04, normal coverage was greater than 7, normal variant allele frequency was no greater than 0.01, and the tumor alternate allele count was greater than 4. Only variants supported by at least two callers were included. Common variants recorded in dbSNP build 142 or gnomAD v2.1.1 and ExAC were excluded. Insertions and deletions located within ENCODE blacklist regions or regions of low mappability were also removed. The filtered somatic mutation profiles were analyzed using maftools v2.12.0 and integrated with epithelial state abundance inferred at the sample level from the corresponding single cell transcriptomic data.

#### Survival analysis

MIA [50] was further performed between NMF cell types and ST samples. After extracting the overlapped features, the survival analysis was conducted to evaluate prognostic value of NMF cell type proportion, NMF cell type featured DEGs, and NMF celltype MIA overlapped features.

#### Attention based multiple-instance learning

We implemented an attention-based multiple-instance learning (MIL) framework [51] to associate histopathologic features with C1 and E2 molecular phenotypes. Image tiles were grouped at the patient level, with individual tiles treated as instances within each patient bag. When multiple samples were available for the same patient, patient-level C1 and E2 labels were assigned by majority voting. Separate binary classifiers were trained to distinguish high-C1 from low-C1 and high-E2 from low-E2 patients based on matched transcriptomic data.

Tile-level features were extracted using an ImageNet-pretrained ResNet50. Tiles were resized to 224 × 224 pixels, normalized using ImageNet channel-wise statistics, and represented by the 2,048-dimensional output of the global average pooling layer.

For MIL aggregation, each 2,048-dimensional tile feature vector was projected into a 512-dimensional latent representation using a fully connected layer followed by ReLU activation and dropout (0.25). Tile-level relevance was estimated using a gated attention mechanism comprising parallel tanh– and sigmoid-activated transformations, followed by element-wise multiplication and linear projection to a scalar attention score.

Attention scores were normalized across tiles within each patient bag using a temperature-scaled softmax function with a temperature parameter of 0.5. The resulting attention weights were used to compute a weighted sum of tile embeddings, generating a patient-level representation that was subsequently passed to a linear classifier for binary prediction. The attention module used a 256-dimensional hidden representation. Patients were divided into training and validation cohorts using an 80:20 stratified split with a fixed random seed of 42. To constrain bag size, up to 256 tiles per patient were used for the C1 classifier and up to 64 tiles per patient for the E2 classifier. During training, tile subsets were sampled from a patient-specific shuffled tile pool, whereas a fixed tile subset was used during validation to ensure deterministic evaluation. Patient bags were processed individually with a batch size of one.

Models were trained for 15 epochs using the AdamW optimizer with a learning rate of 1 × 10−4 and weight decay of 1 × 10−4. Class imbalance was addressed using positive-class-weighted binary cross-entropy loss. Model performance was evaluated using the area under the receiver operating characteristic curve (AUROC) and average precision (AP), and the model checkpoint achieving the highest validation AUROC was retained for downstream analyses. Tile-level attention weights from the selected models were subsequently used to rank histopathologic regions according to their model-assigned relevance for C1 or E2 classification.

#### Spatial transcriptomics library preparation

Spatial transcriptomic data from three paired tumor and adjacent tissue specimens were reanalyzed from GSE285715. The cDNA libraries were constructed and then sequenced on 10 X Visium Spatial platform.

#### Spatial transcriptomics preprocessing and quality control

Space Ranger was used to perform quality control, reads alignment and gene mapping. After processing the ST data, the spots matrices were analyzed using Seurat package [39] with following criteria: (1) ruling out ribosomal and mitochondrial genes; (2) ruling out genes expressed in less than 10 spots; (3) screening spots with criteria nFeature_Spatial_filt>300 & nCount_Spatial_filt>500.

#### Spatial transcriptomic normalization and clustering

SCTransform function [39] was applied to spot normalization and PCA was used for dimensional reduction and spot clustering.

#### RCTD-based spatial deconvolution

Robust Cell Type Decomposition (RCTD) method [52] was used to estimate cell proportions to each spot on ST slices, based on single-cell reference data. We strictly adhered to the standard RCTD process in our workflow, as the RCTD object was created in default parameters. Then the run.RCTD function with the full model was performed to generate the deconvolution results.

#### Spatial tissue-compartment annotation

To integrate scRNA and ST data, we adopted MIA method [50]. After unsupervised clustering of spatial transcriptomic spots, the spatial regions and different types of cells were further integrated via MIA. MIA is proceeded by first delineating sets of cell-type specific and tissue region-specific genes and then determining whether their overlap is higher (enrichment) or lower (depletion) than expected by chance. In scRNA and ST data, the gene sets were set by identifying positive characteristic genes using “FindMarkers” function, with a threshold of logFC=0.5 and p<=0.05. After extracting gene sets in both scRNA and ST data, MIA next computes the overlap between each pair of cell type-specific and region-specific gene sets and performs a hypergeometric test to assess significant enrichment or depletion.

CellTrek is a computational framework that can directly map single cells back to their spatial coordinates in tissue sections based on scRNA-seq and ST data [53]. This method provides a new paradigm that is distinct from ST deconvolution, enabling a more flexible and direct investigation of single cell data with spatial topography.

#### Multiplex immunofluorescence staining and image acquisition

FFPE sections (5 μm) from clinical specimens and murine tissues were deparaffinized, rehydrated, and subjected to multiplex immunofluorescence staining using the Opal 7-Color Manual IHC Kit (Akoya Biosciences, Cat# NEL811001KT). Antigen retrieval was performed by microwave treatment in citrate buffer (pH 6.0) or Tris-EDTA buffer (pH 9.0). Endogenous peroxidase was blocked with 3% H₂O₂ for 10 min. Sections were incubated with primary antibodies overnight at 4°C, followed by HRP-conjugated secondary antibodies (30 min, room temperature) and TSA fluorophore deposition (Opal 520, 570, 620, 650, 690). Antibody stripping was performed between cycles by microwave heating in AR6 buffer (Akoya Biosciences). Nuclei were counterstained with DAPI and slides were mounted with ProLong Diamond Antifade Mountant (Thermo Fisher Scientific). Stained slides were scanned using a Vectra Polaris automated quantitative pathology imaging system (Akoya Biosciences) at 40× magnification.

#### Multiplex immunofluorescence image preprocessing

Multispectral images were unmixed using inForm software (v2.6, Akoya Biosciences), imported into QuPath (v0.4.3) for ROI annotation, and exported for downstream analysis.

#### Single-cell image segmentation & Fluorescence-intensity extraction

Single-cell segmentation was performed using Cellpose 3.0 with the nuclei model applied to the DAPI channel and the cyto2 model applied to pan-cytokeratin. Segmentation masks were generated with a diameter parameter of 10 μm and flow threshold of 0.4. Mean fluorescence intensity (MFI) for each marker in each cell was extracted by overlaying segmentation masks onto corresponding fluorescence channels. Regions of interest (ROIs) were manually defined in FIJI (ImageJ v1.53) based on tissue architecture and pathological features. Cell-type-specific intensity was calculated as the combined mean fluorescence intensity (MFI) of α-SMA and TPM2 for CAFs, and the combined MFI of pan-CK and SPINK1 for tumor cells. For certain analyses, individual channel intensities were directly extracted and quantified. For each ROI, the spatial distribution and intensity profiles of CAFs and tumor cells were visualized using Origin 2025b. Cell density and intensity values were normalized to ROI area and compared across different tissue regions.

#### Cell phenotype assignment

Single-cell intensity matrices and spatial coordinates were imported into Python and converted to AnnData objects using scanpy (v1.9.3). Intensity values were log-transformed (log1p) and z-score normalized. Dimensionality reduction was performed using UMAP, and unsupervised clustering was conducted using the Leiden algorithm (resolution = 0.6). Cell phenotypes were assigned based on cluster-specific marker profiles.

#### Spatial neighborhood enrichment and co-occurrence analysis

Spatial analysis was performed using squidpy (v1.2.3). Neighborhood enrichment analysis, co-occurrence analysis, and spatial autocorrelation (Moran’s I) were computed to assess spatial organization and cell-cell interactions.

The extent of spatial clustering of spots within different types of cells was assessed by the GetSpatNet function in “STUtility” (ver 1.1.1) R package. A network was constructed for the estimation of the spot degree for every included spot, and thereafter the network’s average degree was further calculated. The cell connectivity was identified as: for a specific type of cells, if there are same cells exist in one adjacent spot, the cell connectivity was 1 degree. If there are multiple adjacent spots with the existence of the same cell, the connectivity was the number of adjacent spots. In our research, the maximum number of directly adjacent spots was 6 at a distance of 200 μm (10X Genomic), and the threshold proportion of cell was selected according to the general proportion of cell in each spatial transcriptomic slice.

#### Cell connectivity score

Cell connectivity score indicates the cumulative C1 proportion in 6 direct adjacent spots of the selected spots with E2 > upper quartile threshold in each spot.

#### Bayesian spatial connectivity test

The Bayesian spatial connectivity test involves two key components designed to assess the spatial co-localization and connectivity between E2 and C1, in BLCA ST dataset. These components are evaluated by comparing the observed spatial patterns to what would be expected under a random distribution: 1. probability of co-localization between E2 and C1 in the same spot; 2. probability of C1 colocalization around E2 threshold points

#### Cell distance analysis

Cell centroid coordinates and phenotype annotations were obtained from multiplex immunofluorescence data acquired from 14 BLCA specimens. Within each specimen, Euclidean distances between cell centroids were calculated in the native tissue coordinate system and converted to micrometres according to the spatial calibration of the images. For each cell, the distance to the nearest cell belonging to the specified comparison phenotype was recorded. Four tumor and stromal phenotype combinations were evaluated according to SPINK1 expression in tumor cells and TPM2 expression in CAFs: SPINK1 positive tumor cells with TPM2 positive CAFs, SPINK1 negative tumor cells with TPM2 positive CAFs, SPINK1 positive tumor cells with TPM2 negative CAFs, and SPINK1 negative tumor cells with TPM2 negative CAFs. Distance distributions were summarized across the 14 specimens and compared using the Mann Whitney U test.

#### Tumor stroma boundary analysis

Single cell coordinates and corresponding phenotype annotations derived from 14 multiplex immunofluorescence specimens were used for spatial boundary analysis. For each specimen, tumor and stromal compartments were delineated from the spatial cell map, and the interface separating these compartments was defined as the tumor boundary. A signed distance from each cell centroid to the boundary was calculated, with negative and positive values assigned to the tumor and stromal compartments, respectively. Cells located within 500 μm on either side of the boundary were stratified into consecutive 10 μm intervals. Within each interval, the relative abundance of each annotated cell population was quantified for individual specimens and subsequently aggregated across the cohort to generate boundary centered cell distribution profiles.

#### Directional spatial distribution analysis

Single cell spatial coordinates and phenotype annotations from 14 multiplex immunofluorescence specimens were analyzed to characterize the angular organization of CAF subsets around tumor cell populations. CK19^+^SPINK1^+^Tum, CK19^+^SPINK1^−^Tum, PanCK^+^SPINK1^+^Tum, and PanCK^+^SPINK1^−^Tum populations were designated as reference populations. For each reference population, the relative angular positions of surrounding TPM2^+^ and TPM2^−^ CAFs were determined across the complete 0° to 360° range and partitioned into 12 angular sectors. Sector specific proportions of each CAF subtype were calculated independently for each specimen and summarized across the cohort as the mean ± SEM. The expected distribution under spatial angular uniformity was used as the reference. Directional bias within each sector was quantified as log2[proportion (TPM2^+^ CAF)/proportion (TPM2^−^ CAF)]. Positive and negative values indicated preferential enrichment of TPM2^+^ and TPM2^−^ CAFs, respectively. Sectors in which the SEM did not encompass zero were classified as exhibiting pronounced directional bias.

#### Spaceshapley analysis

In bioinformatics, the Spatial Shapley method is widely applied to uncover the contributions of various features in complex spatial data. This approach combines the classical cooperative game theory of Shapley values with spatial data analysis, quantifying each feature’s contribution to model predictions and providing deep insights into feature interactions and local effects in spatial dimensions.

#### BANKSY-based spatial-domain identification

Spatial domains were identified using BANKSY [54] by integrating molecular features with local spatial context. For Visium data, log normalized expression of 2,000 highly variable genes and standardized RCTD cell state proportions were combined with their 15 nearest spatial neighborhood profiles using (lambda=0.2). Principal component analysis with 50 components, a 20 nearest neighbor graph, and Leiden clustering at resolution 1.5 were applied, yielding six spatial niches. For mIF data, normalized single cell protein profiles were integrated with neighborhood averaged profiles within each specimen. In the TMA cohort, 15 spatial neighbors, (lambda=0.2), cosine distance, and Leiden clustering at resolution 0.5 were used. BANKSY identified 35 and 16 domains in the discovery and TMA cohorts, respectively. Domains were annotated by predominant cell composition and grouped into tumor, stromal, and recurrent functional niches. Domain similarity was evaluated using Pearson correlation and hierarchical clustering.

#### MISTy-based spatial dependency analysis

The implementation of MISTy in mistyR [55] provides an approach to assess the spatial dependency among different types of cells in slices. RCTD conducted the cell proportion estimation in each spatial spot, which was adopted into a multi-faceted model constructed by MISTy, including a discrete spatial context: the intrinsic perspective that evaluates correlations among the deconvolution estimations, within a specific scale. The MISTy model identifies significant structural patterns in the different spatial contexts by associating the available cell-type proportions in each view for each target cell-type proportion in the intraview.

#### Spatial trajectory analysis

To discover the internal evolvement process of tumor cells, the pseudo-time analysis was adopted to spatial transcriptomic data based on “stLearn” package (ver 0.4.10) [56]. stLearn harnesses PAGA trajectory analysis founded on tissue-wide SME normalized gene expression data to uncover connections within subclusters, and the development trajectory of carcinoma in situ in BLCA. Sctour package was also employed.

#### Ligand–receptor and cell–cell communication analysis

The “CellChat” package (ver 1.6.1) provided a method to infer probability of cell-cell communications between specific spatial clusters in ST samples, based on prior knowledge of communications between signaling ligand-receptor pairs and cofactors [57]. Meanwhile, we also employed LIANA to conduct consensus communication analysis among cell clusters. LIANA is a scalable framework that adapts and extends existing methods and knowledge to study cell-cell communication in single-cell, spatially-resolved, and multi-modal omics data [58].

#### Spatial cellular communication flux analysis

COMMOT constructs a collection of cell-cell communication networks through various predefined ligand-receptor pairs from aggregated ligand-receptor interaction databases by solving a global optimization problem that accounts for potential higher-order interactions between the multiple ligand and receptor species [59]. COMMOT analyze different L-R pairs, taking into consideration the spatial distances between specific cells from paired spatial data while also inferring spatial signaling directionality.

#### Receptor activity inference

RIDDEN (Receptor actIvity Data Driven inferENce) integrates prior knowledge of ligand-receptor interactions from the OmniPath database with a large-scale collection of perturbation gene expression profiles from The Library of Integrated Network-Based Cellular Signatures (LINCS) to infer receptor activities from transcriptomic profiles [60]. RIDDEN predicts the activation or suppression state of receptors based on the downstream gene expression, which is adopted to explore the EGFR activity in spatial spots.

#### Metabolite-mediated cell–cell communication analysis

The BLCA scRNA data were analyzed by MEBOCOST, an algorithm predicting metabolite-mediated cell-cell communication for which the metabolites, such as lipids, are secreted by one cell and travel to interact with the sensor protein of another cell [61]. The sensor proteins on receiver cells might be cell surface receptors, transporters across the cell membrane, or nuclear receptors. The sender and receiver cells are inferred based on the expression levels of enzymes producing or catabolizing the metabolites in the sender cells and the expression levels of the sensor proteins in the receiver cells. MEBOCOST is based on the Compass, an algorithm that characterize the metabolic state of cells based on scRNA and flux balance analysis (FBA) [62]. We also employed Compass to characterize cellular metabolic states based on E2 and C1, compared with other epithelial or CAF subtypes.

#### MALDI mass spectrometry imaging

Frozen tissue specimens were transferred from −80°C storage to a precooled CM1950 cryostat (Leica), trimmed to expose the embedded tissue, and cryosectioned at a thickness of 10 μm. Tissue sections were transferred onto ITO-coated conductive glass slides (Bruker; catalog no. 8237001) using a precooled brush, briefly warmed to facilitate adhesion, and vacuum-dried for 30 min. A 9-aminoacridine matrix (9-AA; Sigma-Aldrich) was applied using an HTX-TM Sprayer (HTX Technologies) at a flow rate of 120 μL/min, a nitrogen pressure of 10 psi, a spraying duration of 5 min, and a nozzle speed of 1,200 mm/min. The matrix-coated slides were mounted on the target plate of a timsTOF fleX mass spectrometer (Bruker Daltonics). Tissue regions of interest were defined using flexImaging software (Bruker), and MALDI mass spectrometry imaging data were acquired in negative-ion mode over an m/z range of 150–1,200 at a spatial resolution of 50 μm. The resulting raw imaging data were retained for subsequent spectral preprocessing and metabolite annotation.

#### MALDI-MSI spectral preprocessing and normalization

For ion image reconstructions, the Cardinal package was used to process and visualize the abundance of metabolites in tissue after background subtraction. The data obtained by scanning not only contained the mass and charge data, but also contained the spatial site information. All MS images were normalized using total ion count normalization (TIC) in each pixel.

#### Mass-peak detection and metabolite annotation

Raw negative-ion-mode MALDI-MSI data acquired over an m/z range of 150–1,200 were processed using SCiLS Lab 2024 (Bruker Daltonics). Spectral preprocessing included baseline subtraction, peak alignment, spectral smoothing, and signal normalization. The aligned m/z features and their corresponding pixel-wise ion intensities were retained for downstream analyses, and the mass spectrometry imaging data were converted to imzML format. Metabolite annotation was performed using an in-house R package by matching the observed m/z features against the BiotreeDB mass spectral database. The annotated metabolites were subsequently assigned to chemical classes according to the Human Metabolome Database (HMDB) taxonomy. Because tandem mass spectrometry matching and validation against authentic standards were not documented, all metabolite assignments were considered putative.

#### Hotspot-based spatial metabolite module identification

Z score standardized intensity profiles from tumor derived MALDI MSI pixels were analyzed using Hotspot [63] with the normal model. A Harmony [40] corrected principal component representation was generated with specimen identity as the batch variable, and an unweighted 30 nearest neighbor graph was constructed in this space. Local autocorrelation was evaluated for each MSI feature, and features with a false discovery rate below 0.05 were retained for local correlation analysis. Modules were identified using a minimum size of 20 features, core features only (core_only = True), and a module false discovery rate threshold of 0.001. This procedure yielded 20 modules, for which pixel level scores were calculated. Relationships among modules were evaluated using Pearson correlation and Ward hierarchical clustering. Representative features were ranked according to their correlation with the corresponding module score, and putative pathway associations were assigned using the available metabolite annotations.

#### Gaussian mixture modeling of spatial metabolic states

Pixel level scores for the 20 Hotspot modules were used as input. Missing values were set to zero, modules with variance below (1 times 10^-6) were excluded, and the remaining variables were Z score standardized. Candidate Gaussian mixture models containing 2–10 components were fitted with three initializations and a random seed of 42. Model selection jointly considered the Bayesian information criterion and the Euclidean silhouette score, with the latter estimated from up to 15,000 randomly sampled pixels. A six-component solution was selected. The final full covariance model was fitted with ten initializations and a random seed of 42. Each pixel was assigned to GMM_0–GMM_5 according to its maximum posterior probability, which was retained as the assignment confidence. State level module profiles and Pearson correlations between states were calculated from mean module scores. State associated MSI features were identified using one versus rest Wilcoxon tests with an adjusted (P<0.05) and a log2 fold change greater than zero. Associations between modules and GMM states were additionally summarized using observed to expected scores.

#### Spatial metabolic vector-field analysis

MSI features annotated as potentially exchangeable were subjected to vector field analysis. For each feature, the spatial intensity surface was smoothed using a Gaussian kernel with (sigma=1). Local vectors were derived from intensity differences between neighboring pixels, normalized by interpixel distance, and oriented along the corresponding connecting directions. The resulting vector components were linearly interpolated onto a regular spatial grid. Divergence and curl were calculated from the spatial derivatives of the interpolated field, and Helmholtz decomposition was performed using a Poisson solver based on the discrete cosine transform to separate gradient and rotational components. These quantities were subsequently mapped back to the original pixels. Comparative summaries were generated for the interiors of the six GMM states and all 15 pairwise transition zones using Z score normalized intensity, divergence, and curl. These descriptors characterize the geometry of static MSI intensity distributions and were not interpreted as direct measurements of metabolite movement, production, consumption, transport direction, or intercellular transfer.

#### Multi-omics integration of tumor and stromal metabolic programs

Integration was performed at the levels of metabolic pathways and spatial territories. Previously derived HMR reaction scores for the E1–E4 epithelial states and C1–C4 CAF states were combined with the expression profiles of key metabolic enzymes and transporters, spatial transcriptomic state maps, normalized MALDI MSI metabolite abundance, Hotspot module scores, and GMM state assignments. Features were organized into glycolysis, pyruvate metabolism, the tricarboxylic acid cycle, glutamate and glutamine metabolism, and lipid metabolism. Because spatial transcriptomic and MSI measurements were obtained from serial tissue sections, correspondence between E2 or C1 enriched regions and MSI defined metabolic territories was evaluated qualitatively through regional overlap and pathway concordance, without imposing one to one state matching. HMR scores were interpreted as transcriptome inferred metabolic potential, whereas MSI measurements represented spatial metabolite abundance. The integrated analysis therefore assessed cross modal concordance but did not establish enzyme activity, intracellular metabolic flux, metabolite origin, transport direction, or direct metabolite transfer between tumor cells and CAFs.

## Statistical analyses

Experimental data were derived from triplicate biological replicates and presented as mean ± SD. Statistical evaluations were conducted in GraphPad Prism 9.0 (La Jolla, CA, USA) for parametric (Student’s t-test) or non-parametric (Mann-Whitney U test) comparisons between two groups, while multigroup analyses employed either one-way ANOVA with Bonferroni post hoc tests (for normally distributed data) or Kruskal-Wallis tests. Parametric assumptions (normality and equal variance) were validated prior to ANOVA implementation. Survival curves were generated using R v4.2.2 (R Foundation for Statistical Computing) with the Survival package for Cox proportional-hazards modeling and Survminer package for visualization. A two-tailed α-level of 0.05 defined statistical significance across all analyses.

## Authors’ contributions

L.F.Z., F.D.Y., Q.Y.Y., and Y.F.W. designed the research. Q.Y.Y., Y.F.W., Y.D.D.J., and X.D.H. performed the data analyses. Q.Y.Y., Y.F.W., and L.F.Z. refined the conceptual framework. H.L.L., Y.Lu, and X.D.H. performed the research. Q.Y.Y., Y.F.W., and X.D.H. wrote the original paper. L.F.Z, F.D.Y, Y.Li, H.H., W.C., and J.Y.Z. reviewed this paper.

## Acknowledgments

Not applicable.

