## Supplementary_data for "Spatial organization of a SPINK1-associated tumor-stromal ecosystem links metabolic remodeling to immune restriction in bladder cancer"

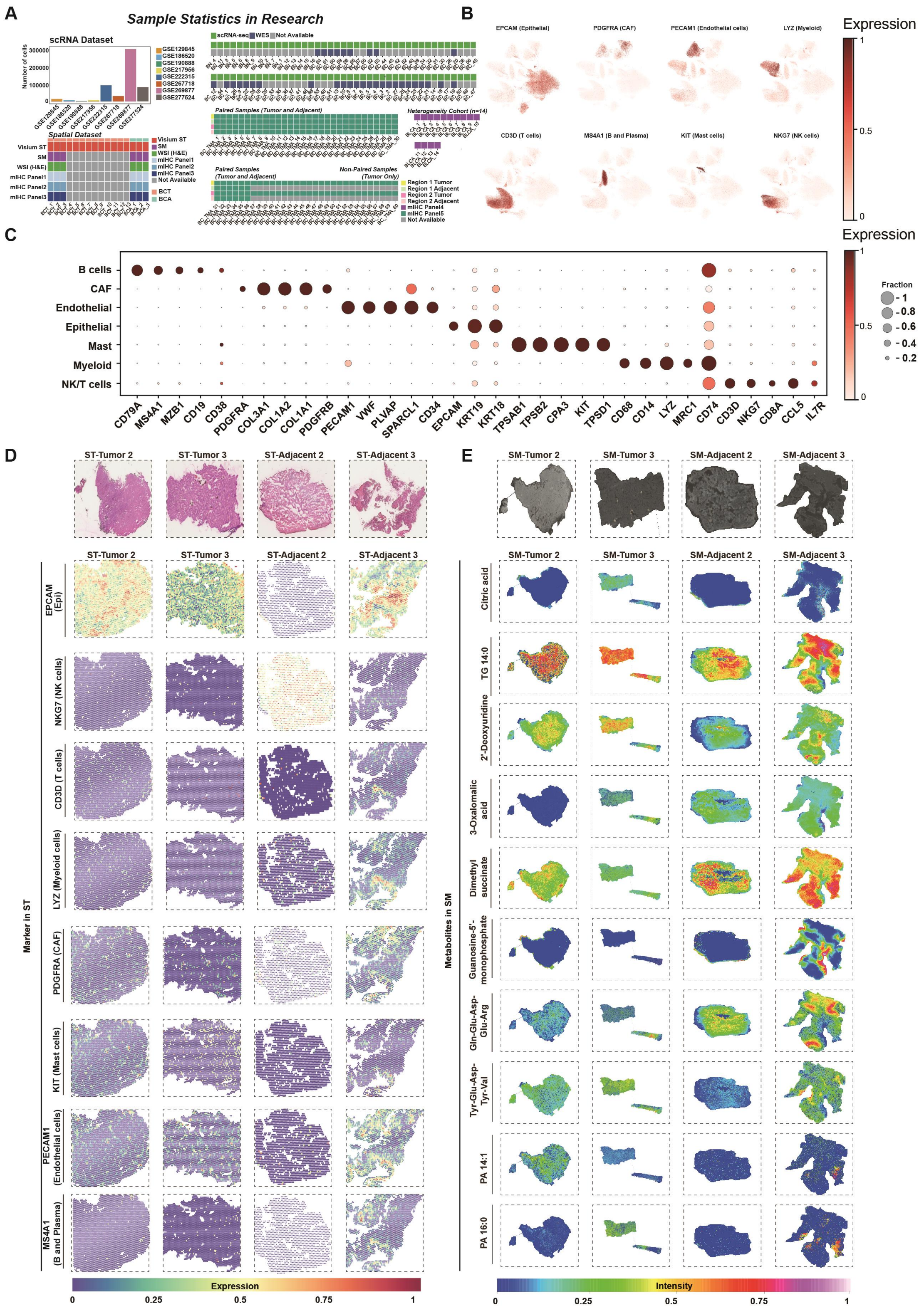

**Figure S1. Cohort overview and multimodal mapping of cellular and metabolic markers.**

(A) Overview of cohort composition, clinical annotations, and sample availability across scRNA-seq, spatial transcriptomics, and spatial metabolomics datasets. Public datasets comprise 82 scRNA-seq samples, 1,238 bulk-transcriptomic samples, 34 WES specimens and 57 Visium profiles. In-house cohorts include six spatial specimens from three patients, 14 mIF specimens and 96 TMA cores from 63 patients. Bars show patient and sample counts; matrices indicate dataset, patient and assay coverage. White dots mark the 23-specimen WES analysis subset. Set, dataset; Pt, patient/group; NR, no corresponding record or unresolved identity.

(B) UMAP feature plots depicting the normalized expression of representative lineage markers within the integrated scRNA-seq atlas.

(C) Dot plot illustrating canonical marker genes used for the annotation of B cells, CAFs, endothelial cells, epithelial cells, mast cells, myeloid cells, and NK/T cells. Dot size represents the proportion of marker-positive cells, whereas color intensity indicates the mean scaled expression level.

(D) Representative H&E images alongside spatial expression maps of the indicated markers across ST sections. Columns correspond to individual specimens, and rows correspond to markers; color scales denote normalized expression levels.

(E) Representative optical images and spatial ion-intensity maps of selected signature metabolites across matched MALDI-MSI sections. Columns correspond to individual specimens, and rows correspond to metabolites; color scales indicate normalized metabolite intensities.

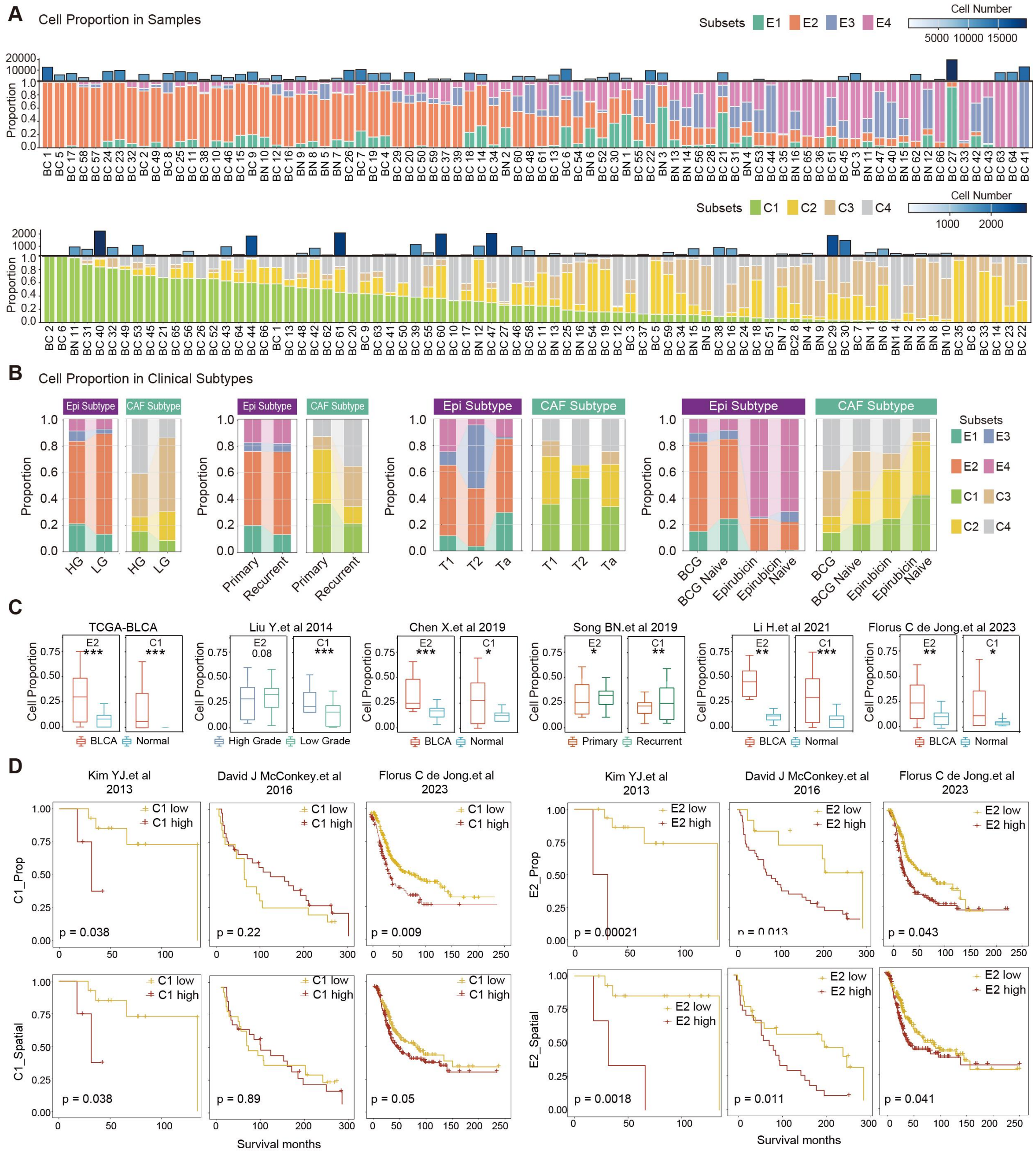

**Figure S2. Clinical distribution and prognostic relevance of NMF-defined epithelial and CAF states in bladder cancer.**

(A) Relative proportions of NMF-defined epithelial states E1–E4 and CAF states C1–C4 across individual bladder cancer (BC) and normal bladder (BN) samples. Bars above the stacked plots indicate the corresponding cell numbers.

(B) Distribution of epithelial and CAF states according to tumor grade, disease status, pathological stage, and treatment category.

(C) Deconvolution-based estimates of E2 and C1 abundance in the TCGA-BLCA, Chen et al. (2019), Li et al. (2021), Liu et al. (2014), Song et al. (2019), and de Jong et al. (2023) bulk transcriptomic cohorts. Comparisons include tumor versus normal tissue, high- versus low-grade tumors, and primary versus recurrent tumors, as indicated.

(D) Kaplan–Meier analyses of patients stratified by high or low C1_Prop, C1_Spatial, E2_Prop, or E2_Spatial scores in the Kim et al. (2013), McConkey et al. (2016), and de Jong et al. (2023) cohorts. P values were determined by log-rank tests and are shown in the panels.

Statistical significance: ns; *P < 0.05; **P < 0.01; ***P < 0.001; ****P < 0.000. Details in STAR Methods.

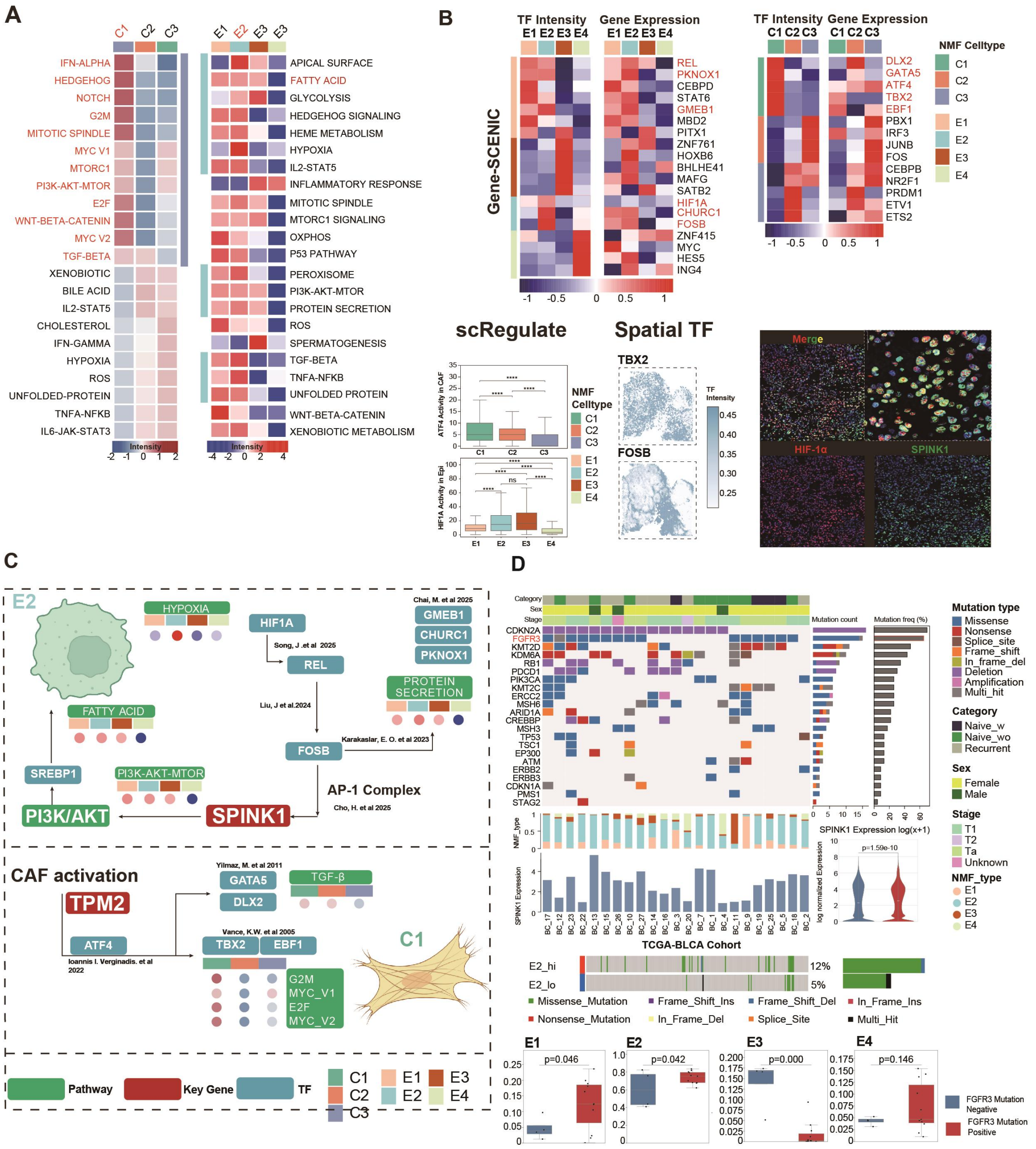


**Figure S3. Functional, regulatory, and genomic features of the E2 epithelial and C1 CAF states.**

(A) Heatmaps of normalized pathway activity across C1–C3 CAF and E1–E4 epithelial states. C1 was enriched in inflammatory, developmental, cell-cycle, and PI3K/AKT/mTOR programs, whereas E2 exhibited prominent hypoxia, fatty-acid metabolism, PI3K/AKT/mTOR signaling, and protein-secretion programs.

(B) Gene-SCENIC analysis of transcription-factor (TF) activity and corresponding gene expression across epithelial and CAF states. scRegulate analyses compare ATF4 activity among C1–C3 and HIF1A activity among E1–E4. Spatial maps show inferred TBX2 and FOSB activities, and multiplex immunofluorescence demonstrates the spatial association of HIF-1α with SPINK1.

(C) Integrated regulatory models of E2 tumor cells and C1 CAFs. The E2 model connects SPINK1 with HIF1A–REL–FOSB/AP-1 and PI3K/AKT–SREBP1 signaling, whereas the C1 model links TPM2 with ATF4, GATA5–DLX2, TBX2–EBF1, and TGF-β-associated CAF activation. Arrows denote literature-supported or computationally inferred relationships.

(D) Somatic mutation landscape of WES-matched samples and associations between FGFR3 status, SPINK1 expression, and epithelial-state composition. SPINK1 expression was compared using a two-sided Wilcoxon rank-sum test. FGFR3 alterations occurred in 12% of E2-high and 5% of E2-low TCGA-BLCA tumors. E1–E4 proportions were compared by one-sided t tests, with exact P values shown.

ns, not significant; ****P < 0.0001.

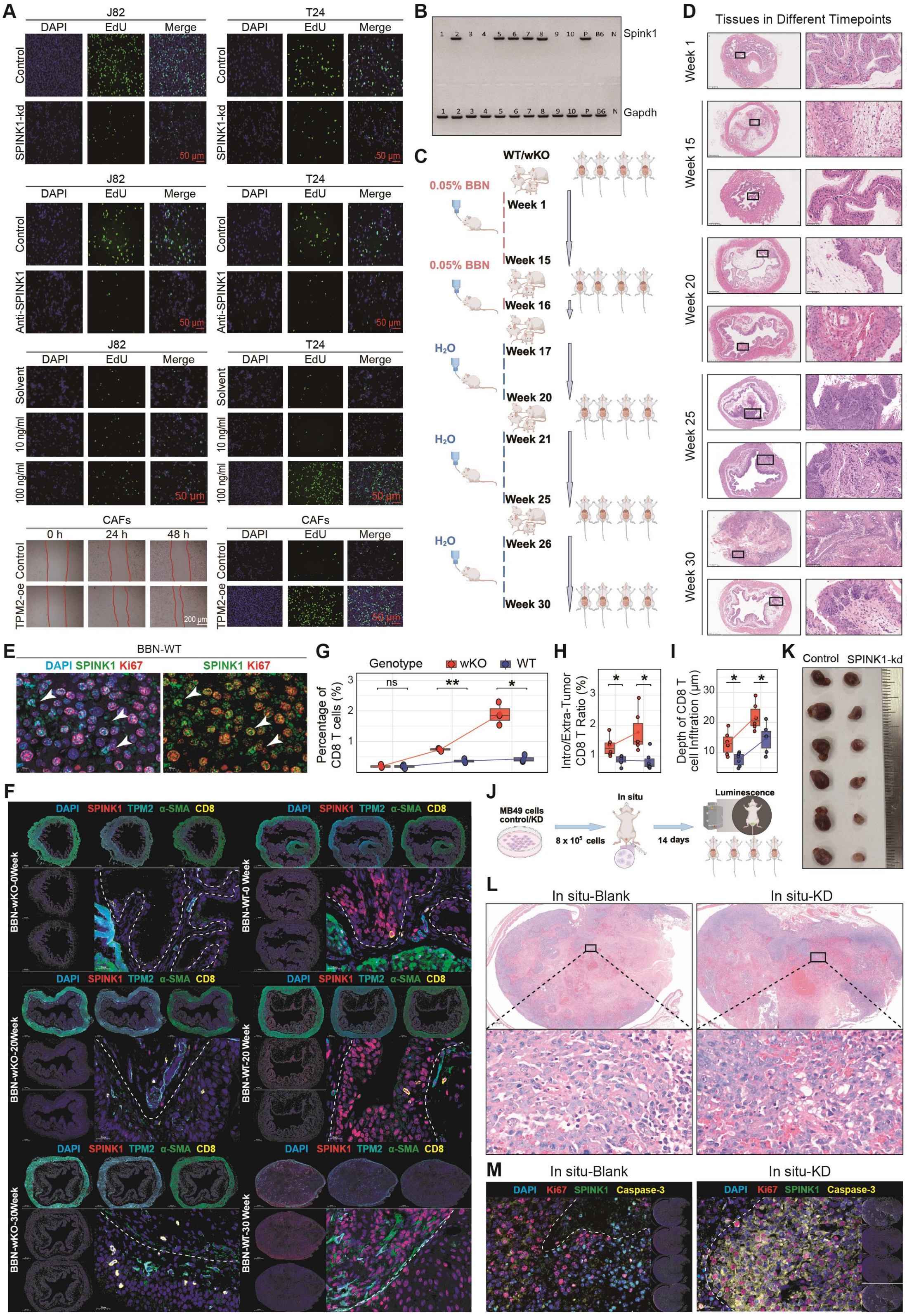

**Figure S4. SPINK1 and TPM2 perturbation modulates cell proliferation, bladder tumor progression, and CD8$^+$ T-cell infiltration.**

(A) Representative EdU incorporation assays in J82 and T24 bladder cancer cells following *SPINK1* knockdown (SPINK1-kd), SPINK1-neutralizing antibody treatment, or recombinant SPINK1 supplementation (10 or 100 ng/mL). Solvent-treated cells served as controls. In parallel, EdU staining and scratch-wound assays were performed in patient-derived CAFs following *TPM2* overexpression (TPM2-oe), with images acquired at 0, 24, and 48 h. Nuclei are shown in blue (DAPI) and proliferating cells in green (EdU). Red lines indicate wound boundaries. Scale bars: 50 μm (EdU) and 200 μm (wound assay).

(B) PCR-based genotyping of tail genomic DNA from the *Spink1* knockout colony. Agarose gel electrophoresis distinguishes *wild-type (WT)* and *Spink1-wKO* alleles via allele-specific amplification.

(C-D) Schematic of BBN-induced bladder tumorigenesis in *WT* and *Spink1-wKO* mice. Mice were exposed to 0.05% BBN followed by regular water and analyzed at defined time points. Representative H&E-stained bladder sections at weeks 1, 15, 20, 25, and 30 are shown at low and high magnification; boxed regions indicate magnified areas.

(E) Representative multiplex immunofluorescence of BBN-induced bladder tumors in WT mice. DAPI, SPINK1, and Ki67 are shown in blue, green, and red, respectively. White arrowheads indicate tumor cells co-expressing SPINK1 and Ki67.

(F) Multiplex immunofluorescence of bladder tissues from *WT* and *Spink1-wKO* mice at baseline and at 20 and 30 weeks post-BBN. DAPI, SPINK1, TPM2, α-SMA, and CD8 are shown in blue, red, cyan, green, and yellow, respectively. Whole-section and magnified views illustrate progressive organization of epithelial/tumor, CAF-enriched stromal, and CD8$^+$ T-cell compartments. White dashed lines demarcate epithelial–tumor boundaries.

(G) Quantification of CD8$^+$ T-cell proportion among all segmented cells in bladder tissues at 0, 20, and 30 weeks post-BBN. *Spink1-wKO* mice exhibited increased CD8$^+$ T-cell abundance at 20 and 30 weeks. Two-sided t tests were used for comparisons.

(H) Ratio of CD8$^+$ T-cell abundance in epithelial–tumor versus stromal compartments at 20 and 30 weeks post-BBN. Elevated ratios in *Spink1-wKO* mice indicate preferential intratumoral accumulation. Two-sided t tests were applied.

(I) CD8$^+$ T-cell infiltration depth, defined as the distance from each CD8$^+$ cell to the epithelial–tumor boundary, at 20 and 30 weeks. *Spink1-wKO* tumors exhibited deeper CD8$^+$ T-cell penetration than WT tumors. Two-sided t tests were used.

(J) Experimental design of the orthotopic MB49 bladder cancer model. Control or *Spink1*-knockdown MB49 cells ($8 \times 10^5$ per mouse) were instilled into the bladder, and tumor formation was assessed by bioluminescence imaging at day 14.

(K) Gross morphology of excised bladders bearing orthotopic tumors derived from control or *Spink1*-knockdown MB49 cells.

(L) Representative H&E-stained sections of orthotopic MB49 tumors. Insets indicate magnified regions.

(M) Multiplex immunofluorescence of orthotopic tumors. DAPI, Ki67, SPINK1, and Caspase-3 are shown in blue, red, green, and yellow, respectively. *Spink1* knockdown resulted in near-complete loss of SPINK1 expression and increased Caspase-3 signal relative to controls. White dashed lines delineate tumor regions.

For (G–I), each point represents an individual mouse. Statistical significance was assessed using two-sided t tests (ns; *P < 0.05; **P < 0.01; ***P < 0.001; ****P < 0.000).

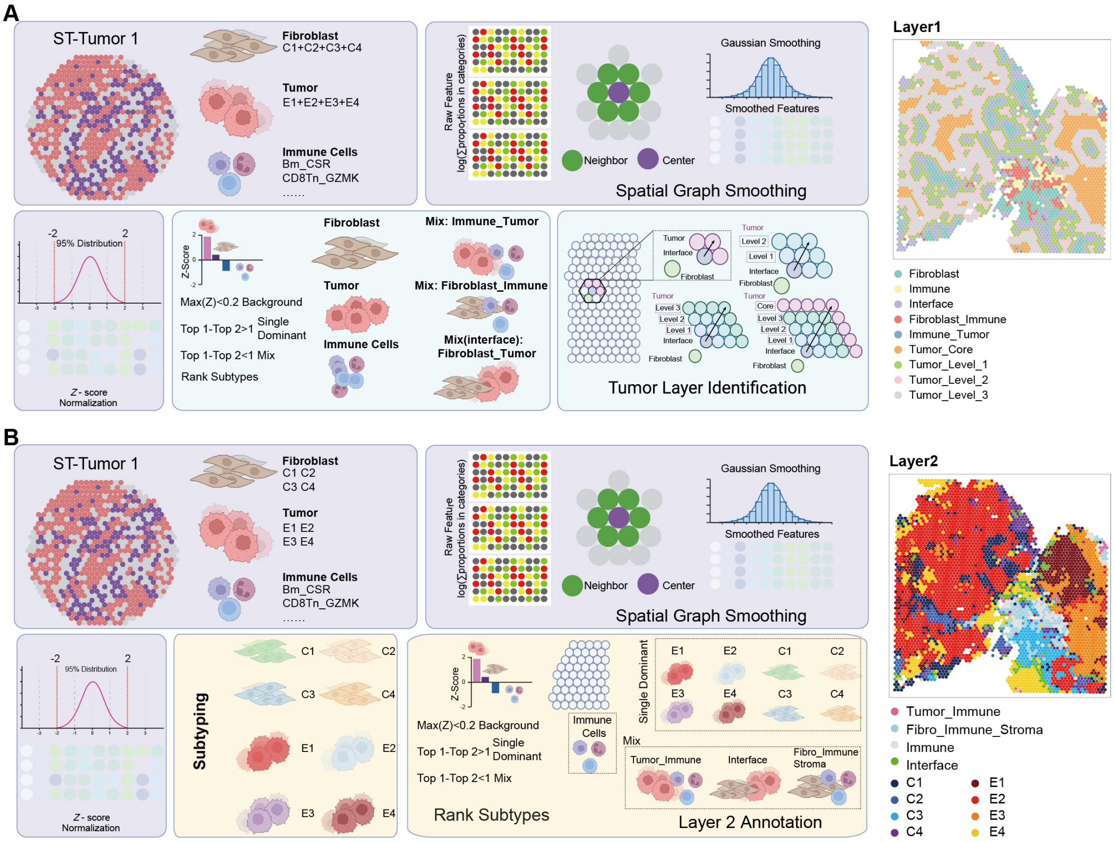


**Figure S5. Hierarchical RCTD-based annotation of tissue compartments and dominant cellular states in Visium spatial transcriptomic data.**

(A) Workflow for delineating tumor, stromal, immune, and tumor–stroma interface regions using RCTD-inferred cell-type proportions. For each Visium spot, RCTD-derived proportions were first aggregated into three major compartments: fibroblasts (CAF states C1–C4), tumor epithelial cells (E1–E4), and immune cells. Compartmental proportions were log-transformed and mapped onto the spatial graph. To reduce local noise while preserving spatial structure, values were Gaussian-smoothed using each spot and its neighbors, followed by Z-score normalization across all spots (central ~95% range: Z = −2 to 2).

Spots with a maximum compartment Z score < 0.2 were defined as background. Among remaining spots, a Z-score difference > 1 between the top two compartments indicated a single-dominant state and was assigned as fibroblast, tumor, or immune. A difference ≤ 1 indicated mixed composition. Mixed tumor_immune and fibroblast_immune spots were annotated as Immune_Tumor and Fibroblast_Immune, respectively. Spots with comparable tumor and fibroblast signals were defined as Fibroblast_Tumor and used to delineate the tumor–stroma interface.

Tumor spatial depth was then inferred based on graph proximity to the interface. Tumor spots directly adjacent to the interface were defined as Tumor_Level_1, with successive inward layers annotated as Tumor_Level_2 and Tumor_Level_3. Tumor regions beyond these layers were classified as Tumor_Core. The final Layer 1 annotation includes fibroblast, immune, interface, fibroblast–immune, immune–tumor, tumor core, and three tumor depth levels. The representative

ST-Tumor 1 map illustrates the resulting spatial organization. This framework preserves transitional states and avoids binary tumor–stroma classification.

(B) Workflow for assigning dominant fine-grained cellular identities at the spot level. Unlike compartment-level aggregation in (A), RCTD proportions were retained for individual epithelial states (E1–E4), CAF states (C1–C4), and immune subpopulations. Each subtype proportion was log-transformed, spatially smoothed, and Z-score normalized using the same procedure as in (A). Within each spot, subtype scores were ranked. Spots with a maximum Z score < 0.2 were labeled as background. For informative spots, a Z-score difference > 1 between the top two subtypes defined a single-dominant identity, assigned to the highest-ranking epithelial or CAF state (E1–E4 or C1–C4). Spots without a clear dominant subtype were retained as mixed states. Tumor–immune mixtures were annotated as Tumor_Immune, fibroblast–immune mixtures within stroma as Fibro_Immune_Stroma, tumor–fibroblast mixtures as Interface, and immune-dominant spots lacking epithelial or CAF dominance as Immune. The representative Layer 2 map shows the spatial distribution of dominant epithelial, CAF, immune, and mixed states. As each Visium spot may contain multiple cells, these annotations represent RCTD-inferred dominant compositional states rather than single-cell identities.

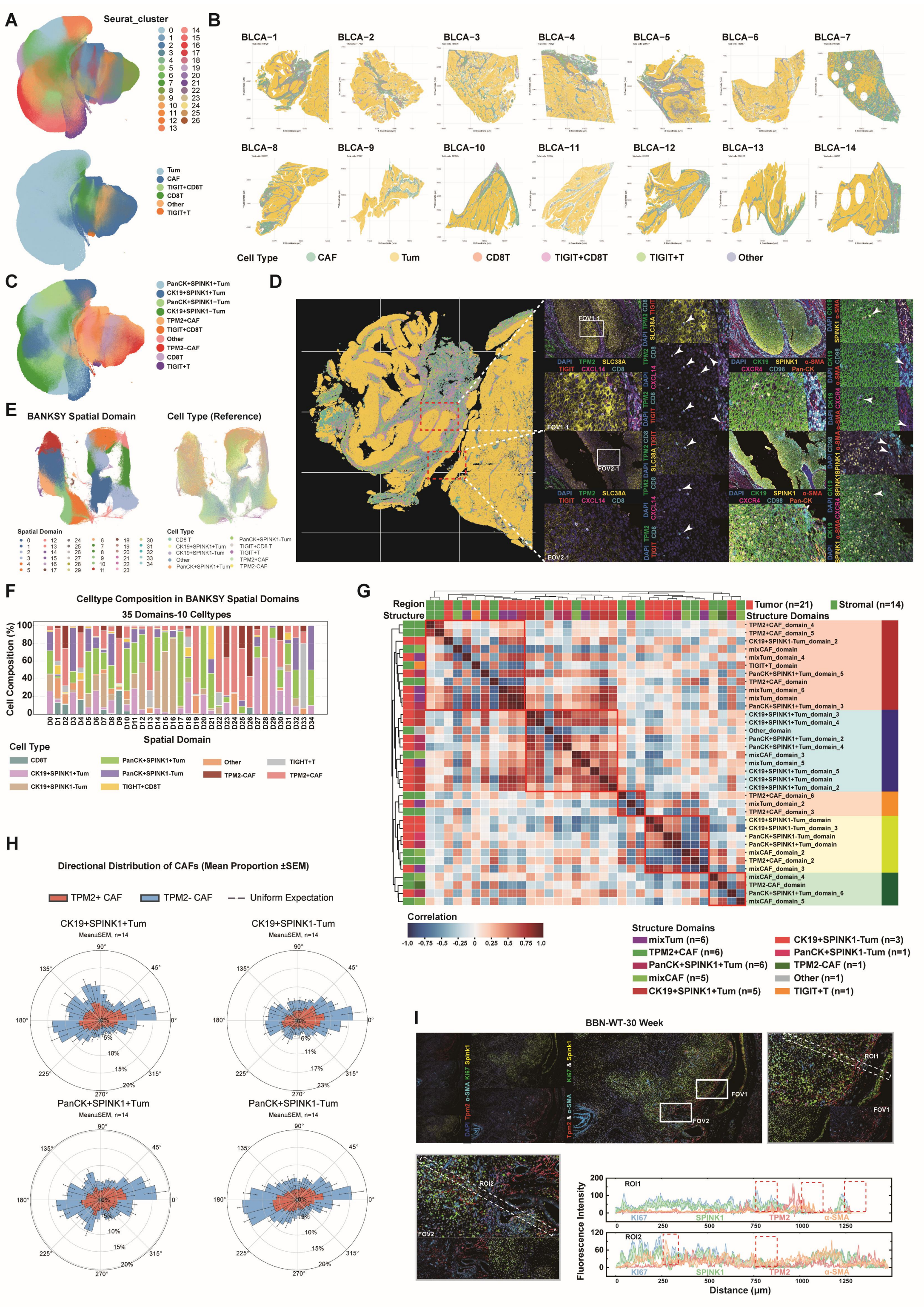

**Figure S6. High-plex spatial proteomics resolves the organization of SPINK1+ tumor cells and TPM2+ CAFs in human and BBN-induced bladder tumors.**

(A) UMAP representation of single-cell protein-abundance profiles quantified by high-plex multiplex immunofluorescence (mIF) across the bladder cancer cohort. Each point represents one segmented cell. Cells are colored by Leiden cluster (clusters 0–26) or by broad cell-type annotation, including tumor cells (Tum), cancer-associated fibroblasts (CAFs), CD8 T cells (CD8T), TIGIT-positive CD8 T cells (TIGIT+CD8T), other TIGIT-positive T cells (TIGIT+T),

and other cell types.
(B) Spatial distribution of the broadly annotated cell populations across 14 human bladder cancer specimens (BLCA-1–BLCA-14). Individual cells are plotted at their native tissue coordinates and colored according to the cell-type annotations defined in (A), illustrating inter-sample heterogeneity in the abundance and spatial organization of tumor, stromal, and immune compartments.
(C) Refined cell-state annotations projected onto the integrated UMAP. Tumor cells were subdivided according to PanCK, CK19, and SPINK1 expression into PanCK+SPINK1+Tum, CK19+SPINK1+Tum, PanCK+SPINK1−Tum, and CK19+SPINK1−Tum populations. CAFs were classified as TPM2+CAF or TPM2−CAF, while immune populations comprised CD8T, TIGIT+CD8T, and TIGIT+T cells. Cells not assigned to these populations were designated "Other."
(D) Detailed spatial organization and protein-expression patterns in the representative BLCA-1 specimen. Left, spatial map of the refined cell identities defined in (C), with each segmented cell displayed at its native tissue coordinate. Boxes identify representative ROIs selected for high-resolution examination. Right, representative multiplex images and enlarged fields of view (FOV1-1 and FOV2-1) showing the spatial distributions of DAPI, TPM2, SLC38A1, TIGIT, CXCL14, and CD8, together with a complementary antibody panel comprising DAPI, CK19, SPINK1, α-SMA, CXCR4, CD98, and PanCK. Channel-resolved and merged images illustrate the localization of tumor, CAF, and immune markers and their organization at tumor–stromal interfaces. White arrowheads indicate representative cells or interfaces displaying the indicated marker patterns.
(E) Identification of spatially organized tissue domains using BANKSY. Cells in a common two-dimensional embedding are colored according to the BANKSY-derived spatial-domain assignment or the refined reference cell identity. BANKSY domains were annotated according to their predominant cellular composition, including CK19+SPINK1+Tum, CK19+SPINK1−Tum, PanCK+SPINK1+Tum, PanCK+SPINK1−Tum, TPM2+CAF, TPM2−CAF, mixed-tumor (mixTum), mixed-CAF (mixCAF), TIGIT+T, and other domains. Comparison with the reference annotations demonstrates that the BANKSY domains capture both dominant cell identities and multicellular spatial organization.
(F) Cellular composition of the 35 BANKSY-derived spatial domains (D0–D34). Each stacked bar represents one spatial domain and shows the percentage contribution of the ten refined cell populations. The marked differences in cellular composition distinguish tumor-dominant, CAF-dominant, immune-enriched, and mixed multicellular domains.
(G) Pairwise correlation structure among the 35 spatial domains. Rows and columns represent individual BANKSY domains, which were hierarchically clustered according to their domain-level cellular profiles. The color scale denotes correlation coefficients ranging from −1 to 1, with blue and red indicating negative and positive correlations, respectively. Annotation tracks indicate the anatomical region assignment, tumor (n = 21) or stromal (n = 14), and the corresponding structural-domain class. Outlined blocks identify groups of domains with internally similar cellular organization. The resulting classes comprise mixTum (n = 6), TPM2+CAF (n = 6), PanCK+SPINK1+Tum (n = 6), mixCAF (n = 5), CK19+SPINK1+Tum (n = 5), CK19+SPINK1−Tum (n = 3), PanCK+SPINK1−Tum (n = 1), TPM2−CAF (n = 1), Other (n = 1), and TIGIT+T (n = 1) domains.

(H) Directional distribution of TPM2+ and TPM2− CAFs around distinct tumor-cell populations. Polar histograms show the angular distributions of TPM2+ CAFs (red) and TPM2− CAFs (blue) surrounding CK19+SPINK1+Tum, CK19+SPINK1−Tum, PanCK+SPINK1+Tum, and PanCK+SPINK1−Tum populations. Angular sectors span 0°–360°, and radial values represent the mean proportion of CAFs detected in each direction. Bars and error bars denote mean ± SEM across 14 specimens, and the dashed circle indicates the expectation under a uniform angular distribution. TPM2+ CAFs exhibited a comparatively even angular distribution around SPINK1-positive tumor populations, consistent with a more circumferential organization of TPM2+ CAFs around SPINK1-positive tumor regions. This pattern supports a spatial association but does not by itself establish directed recruitment or a causal interaction.

(I) Spatial validation in a wild-type mouse bladder tumor collected 30 weeks after induction with N-butyl-N-(4-hydroxybutyl) nitrosamine (BBN-WT, 30 weeks). Representative low-magnification and merged mIF images show DAPI, TPM2, α-SMA, Ki67, and SPINK1. White boxes identify FOV1 and FOV2, which are shown at higher magnification as ROI1 and ROI2. Dashed lines indicate the trajectories used for fluorescence line-scan analysis. Corresponding plots show Ki67, SPINK1, TPM2, and α-SMA fluorescence intensities as a function of distance along each trajectory. The images and line profiles demonstrate the localization of SPINK1 and Ki67 within tumor regions, the association of TPM2 and α-SMA with CAF-rich stromal regions, and the close spatial apposition of these tumor and stromal compartments in the BBN-induced model.

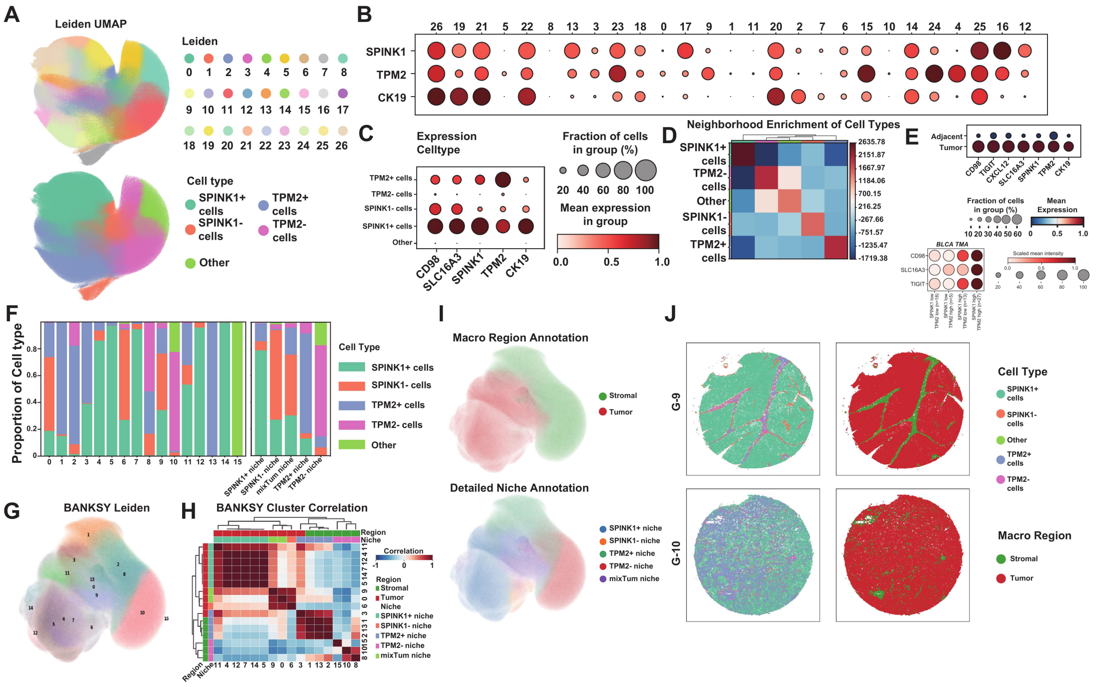


**Figure S7. High-plex spatial proteomic profiling of bladder cancer TMAs delineates SPINK1- and TPM2-associated cellular niches and their functional interactions.**

(A) Unsupervised identification and annotation of protein-defined cellular populations in bladder cancer TMAs. Following cell segmentation of multiplex immunofluorescence images, per-cell protein abundance matrices were normalized, embedded using UMAP, and subjected to Leiden clustering. Each point represents a single cell. The upper UMAP depicts 27 Leiden clusters (0–26), whereas the lower UMAP presents corresponding phenotypic annotations, including SPINK1-positive, SPINK1-negative, TPM2-positive, TPM2-negative, and other cellular states.

(B) Marker expression patterns across Leiden clusters. Dot plots illustrate the distribution of SPINK1, TPM2, and CK19 across clusters 0–26. Dot size denotes the proportion of marker-positive cells, and color intensity reflects mean normalized expression levels. Cluster annotation was guided by epithelial identity (CK19) and relative expression of SPINK1 and TPM2.

(C) Protein abundance across annotated cellular classes. Dot plots summarize CD98, SLC16A3, SPINK1, TPM2, and CK19 across TPM2-positive, TPM2-negative, SPINK1-positive, SPINK1-negative, and other cell populations. Dot size indicates the fraction of positive cells, and color represents mean normalized abundance.

(D) Spatial neighborhood enrichment among cellular classes. The heat map depicts pairwise spatial proximity enrichment or depletion relative to a randomized null model. Red indicates significant enrichment, whereas blue indicates depletion. Hierarchical clustering organizes cell classes according to similarity in spatial interaction profiles. Diagonal enrichment reflects preferential intra-class spatial aggregation.

(E) Protein abundance across TMA compartments and SPINK1/TPM2-defined groups. Dot plots compare seven proteins between tumor and adjacent non-tumor tissues, and CD98, SLC16A3, and TIGIT across four SPINK1/TPM2 high/low tumor groups using all cells. Dot size indicates the fraction of cells with intensity >0; color denotes mean abundance, scaled per protein in the four-group comparison.

(F) Spatially informed clustering using BANKSY. BANKSY integrates protein expression with local spatial context prior to Leiden clustering. The UMAP projection reveals 16 spatial domains (0–15), capturing both molecular and tissue architectural features.

(G) Cellular composition of BANKSY-derived spatial domains and ecological niches. Left, stacked bar plots depict the proportional composition of SPINK1-positive, SPINK1-negative, TPM2-positive, TPM2-negative, and other cells across 16 spatial domains. Right, aggregation of domains into five ecological niches based on dominant cellular composition and spatial context.

(H) Correlation structure of BANKSY spatial domains. The heat map shows pairwise correlations among 16 domains, with red indicating positive and blue indicating negative correlations. Hierarchical clustering identifies groups of domains with shared spatial-proteomic signatures. Annotations denote tumor and stromal macro-regions and corresponding niche identities.

(I) Projection of spatial domains onto the BANKSY UMAP. The upper panel distinguishes tumor and stromal macro-regions. The lower panel illustrates refined ecological niches, highlighting transitions between SPINK1- and TPM2-associated states within the spatial manifold.

(J) Spatial mapping of cellular populations in paired TMAs derived from the same patient (G9: tumor; G10: adjacent non-tumor tissue). Left, cells are displayed in native spatial coordinates and colored by phenotypic class. Right, macro-region classification delineates tumor and stromal compartments, revealing spatial reorganization across tissue contexts.

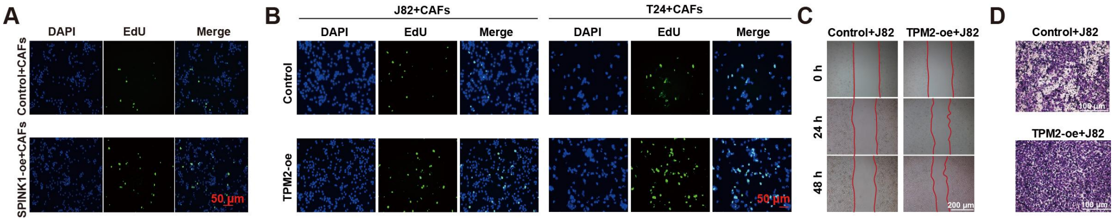


**Figure S8. Effects of SPINK1/TPM2 on proliferation and migration in CAF co-cultures.**

(A) EdU incorporation assay in patient-derived cancer-associated fibroblasts (CAFs) co-cultured with control or *SPINK1*-overexpressing T24 cells. DAPI (blue) labels nuclei, and EdU (green) marks proliferating cells. Scale bar, 50 μm.
(B) EdU incorporation assays in J82 and T24 cells co-cultured with CAFs transduced with control or *TPM2* overexpression constructs. DAPI (blue) and EdU (green) indicate nuclei and proliferating tumor cells, respectively. Scale bar, 50 μm.
(C) Scratch-wound assay of J82 cells co-cultured with control or *TPM2*-overexpressing CAFs. Representative images at 0, 24, and 48 h illustrate progressive wound closure. Red lines delineate wound boundaries. Scale bar, 200 μm.
(D) Matrigel invasion assay of J82 cells co-cultured with control or *TPM2*-overexpressing CAFs. Migrated cells on the lower surface of the membrane were quantified to assess migratory capacity. Scale bar, 100 μm.

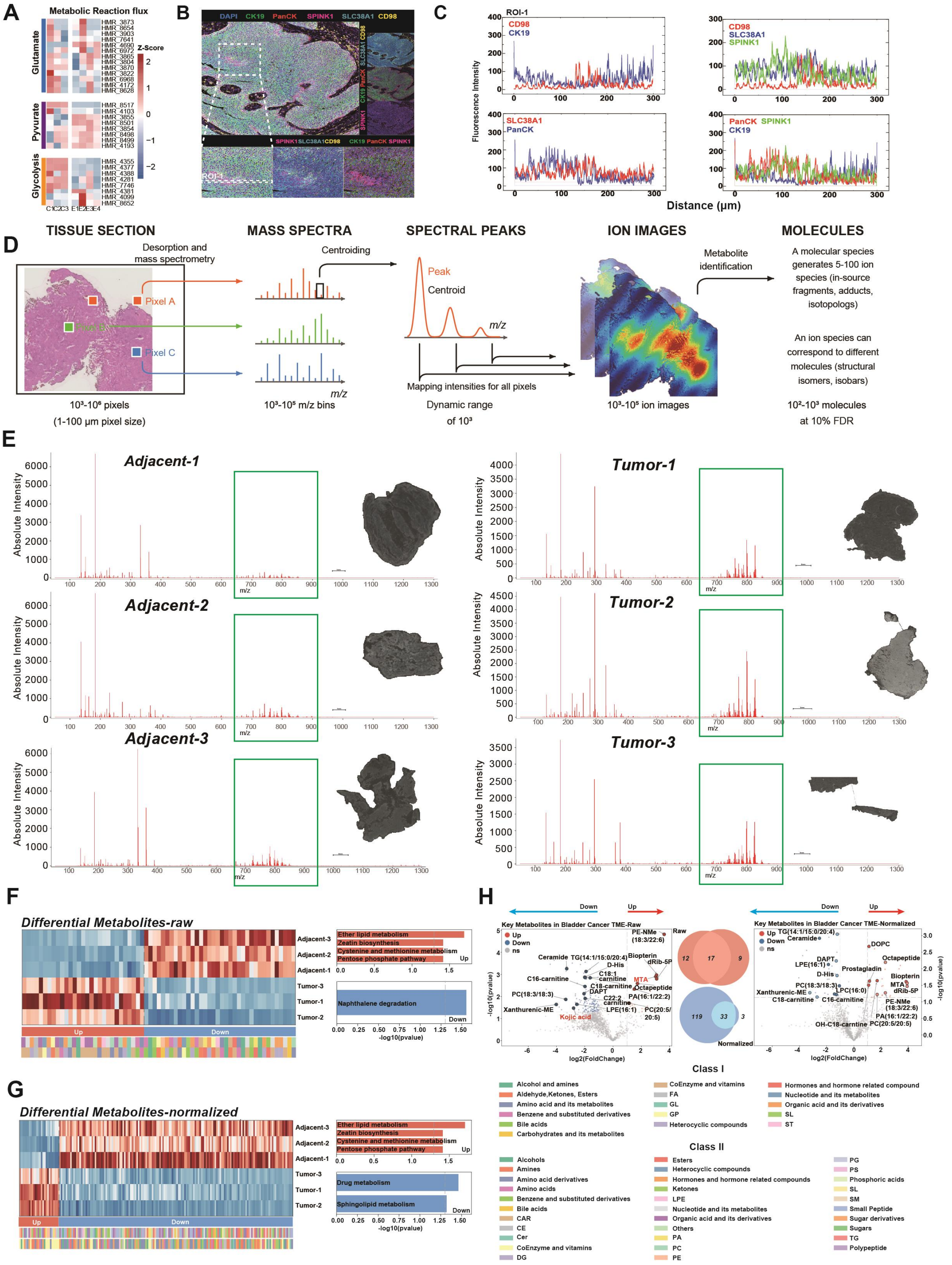


**Figure S9. MALDI-MSI profiling identifies concordant metabolic alterations between bladder tumors and adjacent tissues.**

(A) Metabolic reaction flux. Inferred reaction flux profiles comparing E2 and C1.

(B) Representative multiplex immunofluorescence of a bladder cancer tissue section stained for DAPI (blue), CK19 (green), pan-cytokeratin (PanCK; red), SPINK1 (magenta), SLC38A1 (cyan), and CD98 (yellow). The white dashed box delineates ROI1; enlarged merged and selected-channel views are shown at right and below.

(C) Fluorescence line-scan profiles across ROI1 showing the spatial distributions of CD98 and

CK19; CD98, SLC38A1, and SPINK1; SLC38A1 and PanCK; and PanCK, SPINK1, and CK19. Fluorescence intensity is plotted against distance along the indicated tissue trajectory, illustrating the spatial correspondence between SPINK1-positive tumor regions and metabolic transporter abundance.

(D) Schematic of the MALDI-MSI workflow. Tissue sections are matrix-coated and analyzed by laser desorption/ionization to acquire pixel-wise mass spectra. Spectral centroiding yields discrete m/z features, which are spatially reconstructed into ion images. Putative metabolite annotations are assigned to detected ions, with potential ambiguity arising from in-source fragments, adducts, isotopologues, isomers, and isobars.

(E) Representative MALDI-MSI overview images of three tumor-adjacent tissues (Adjacent-1–3) and three bladder tumors (Tumor-1–3). Scale bars are indicated. Aggregated total-ion mass spectra across all pixels for each specimen. Ion intensity is plotted against m/z. The m/z 650–900 range (green box) contains multiple metabolite-associated features.

(F) Differential metabolic profiling using raw MALDI-MSI intensities. Heatmap shows relative abundance of differential features across specimens and ions. Red and blue indicate higher and lower abundance, respectively. Features are ordered by tumor-associated increase or decrease. Class I and Class II annotations denote chemical superclasses and subclasses. Enrichment analysis of tumor-upregulated features highlights ether-lipid metabolism, purine biosynthesis, cysteine and methionine metabolism, and the pentose phosphate pathway; tumor-downregulated features are associated with naphthalene degradation. Significance is reported as −log10P.

(G) Differential analysis after intensity normalization. Heatmap organization and annotations are as in (F). Tumor-upregulated features are enriched in ether-lipid metabolism, purine biosynthesis, cysteine and methionine metabolism, and the pentose phosphate pathway, whereas tumor-downregulated features are associated with drug metabolism and sphingolipid metabolism. Dashed lines indicate significance thresholds.

(H) Volcano plots of differential features from raw and normalized datasets. x-axis: log2(fold change, tumor vs. adjacent); y-axis: −log10(P). Selected metabolites are labeled. Venn diagrams summarize overlap between analyses, identifying 17 consistently increased and 33 consistently decreased features. Class I and Class II legends indicate chemical classification levels.

All analyses include three tumor and three adjacent samples. Each heatmap row represents one specimen; each volcano point represents one ion feature. Statistical testing, normalization, enrichment analysis, multiple-testing correction, and annotation were performed as described in STAR Methods. As individual m/z features may correspond to multiple chemical entities, annotations remain putative unless validated by tandem MS and authentic standards.

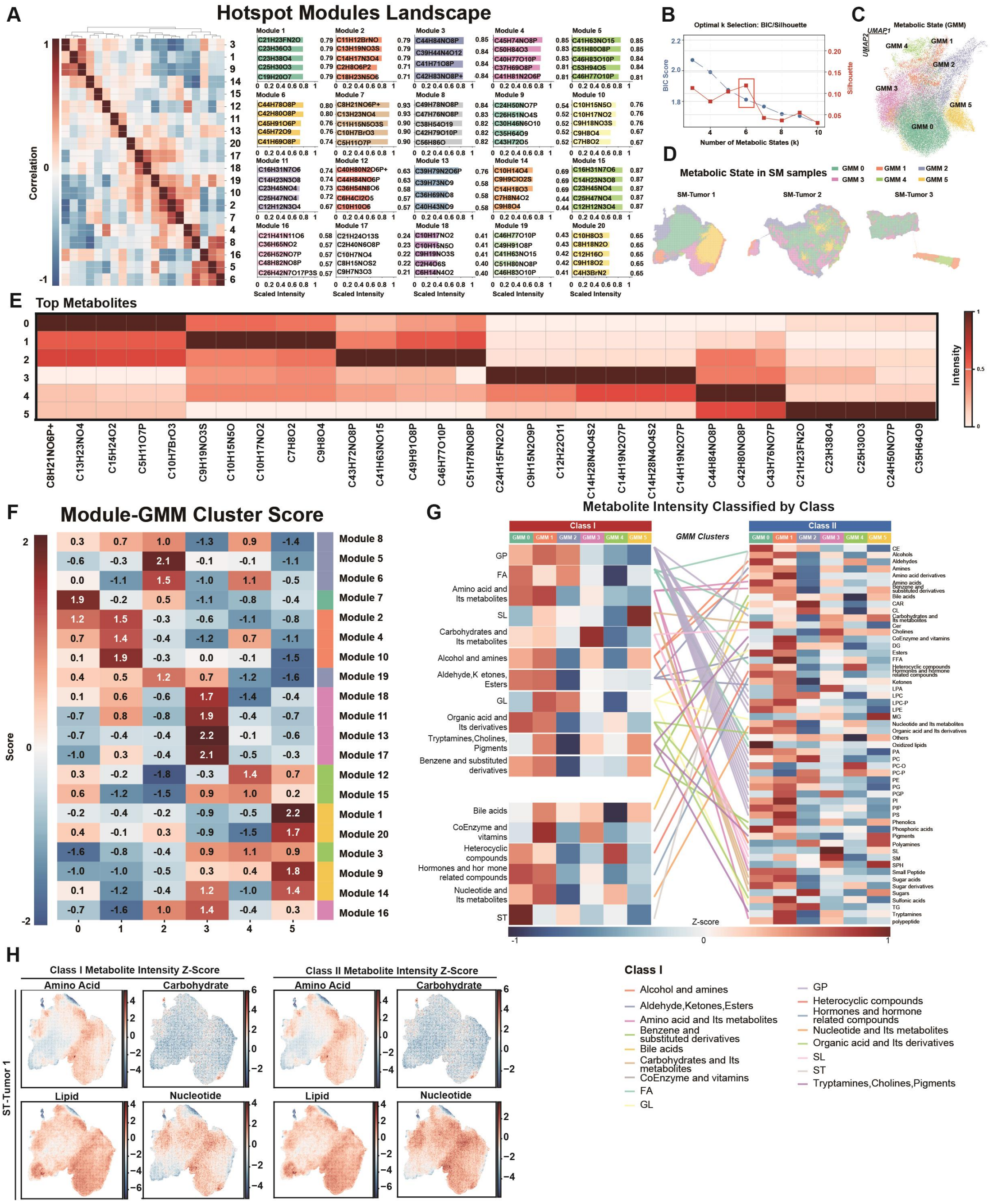


**Figure S10. Hotspot–GMM modeling delineates spatial metabolic states and SPINK1-responsive metabolic programs in bladder cancer.**

(A) Hotspot analysis identified spatially coordinated metabolite modules. A hierarchically clustered correlation matrix of 20 modules revealed distinct positive and negative inter-module relationships. Representative ion profiles within each module are shown, with scaled intensities and within-module association coefficients.

(B) Model selection for Gaussian mixture modeling (GMM) was guided by Bayesian information criterion and silhouette scores across candidate cluster numbers. Integrating model fit and cluster separability supported a six-state solution (k = 6).

(C) UMAP embedding of MALDI-MSI pixels colored by the six GMM-derived metabolic states (GMM 0–5), representing distinct spatial metabolic phenotypes.

(D) Spatial mapping of GMM state assignments in three bladder tumor specimens (SM-Tumor

1–3), demonstrating sample-specific organization of metabolic states.

(E) Heatmap of representative marker metabolites across GMM states, showing state-specific enrichment patterns based on normalized ion abundances.

(F) Association between Hotspot-derived modules and GMM states, quantified using R/OE scores. Positive and negative values indicate preferential enrichment or depletion of modules within specific metabolic states, respectively.

(G) Metabolite-class composition across six GMM-defined metabolic states. Heatmaps show Z-score–normalized abundances of Class I metabolite categories and corresponding Class II subclasses for GMM 0–5. Links indicate hierarchical relationships. Red and blue denote relative enrichment and depletion, respectively.

(H) Spatial distributions of aggregate Class I metabolite scores in representative bladder tumor sections. Maps display Z-score–normalized intensities of amino acid-, carbohydrate-, lipid-, and nucleotide-associated metabolites. Red and blue indicate higher and lower relative abundance.

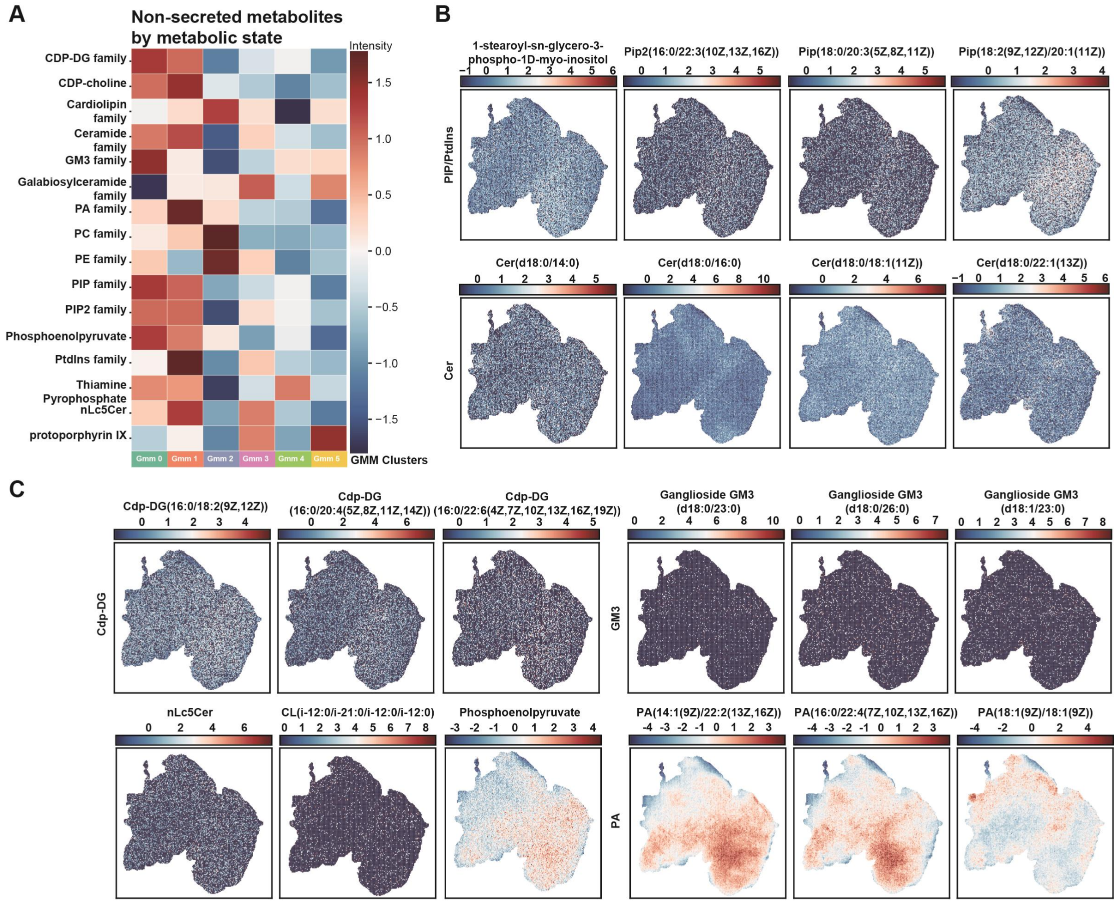


**Figure S11. Spatial metabolite-class mapping reveals PI3K-associated lipid organization and restricted metabolic exchange in bladder cancer.**

(A) Heatmap of non-secreted or poorly exchangeable metabolite families across GMM 0–5, including PI3K/phosphatidylinositol-axis lipids (PIP, PIP2, PtdIns), CDP-DG, CDP-choline, cardiolipin, ceramide, GM3, glycosphingolipids, PA, PC, and PE, as well as selected intracellular metabolites. Values represent normalized intensities per metabolic state.

(B) Spatial ion-intensity maps of representative phosphatidylinositol- and membrane-signaling lipids. The upper panel shows 1-stearoyl-sn-glycero-3-phospho-1D-myo-inositol and representative PIP/PIP2 species; the lower panel shows ceramide species with distinct acyl chains.

Color scales indicate relative ion intensity.

(C) Spatial distributions of representative non-secreted or poorly exchangeable metabolites, including CDP-DG, GM3, nLc5Cer, cardiolipin, phosphoenolpyruvate, and phosphatidic acid species. These patterns define metabolically distinct intratumoral regions not resolved by bulk averages.

CDP-DG, cytidine diphosphate-diacylglycerol; CL, cardiolipin; FA, fatty acyls; GL, glycerolipids; GMM, Gaussian mixture model; GP, glycerophospholipids; MALDI-MSI, matrix-assisted laser desorption/ionization mass spectrometry imaging; PA, phosphatidic acid; PC, phosphatidylcholine; PE, phosphatidylethanolamine; PIP, phosphatidylinositol phosphate; PIP2, phosphatidylinositol bisphosphate; PtdIns, phosphatidylinositol; SL, sphingolipids; ST, sterol lipids.

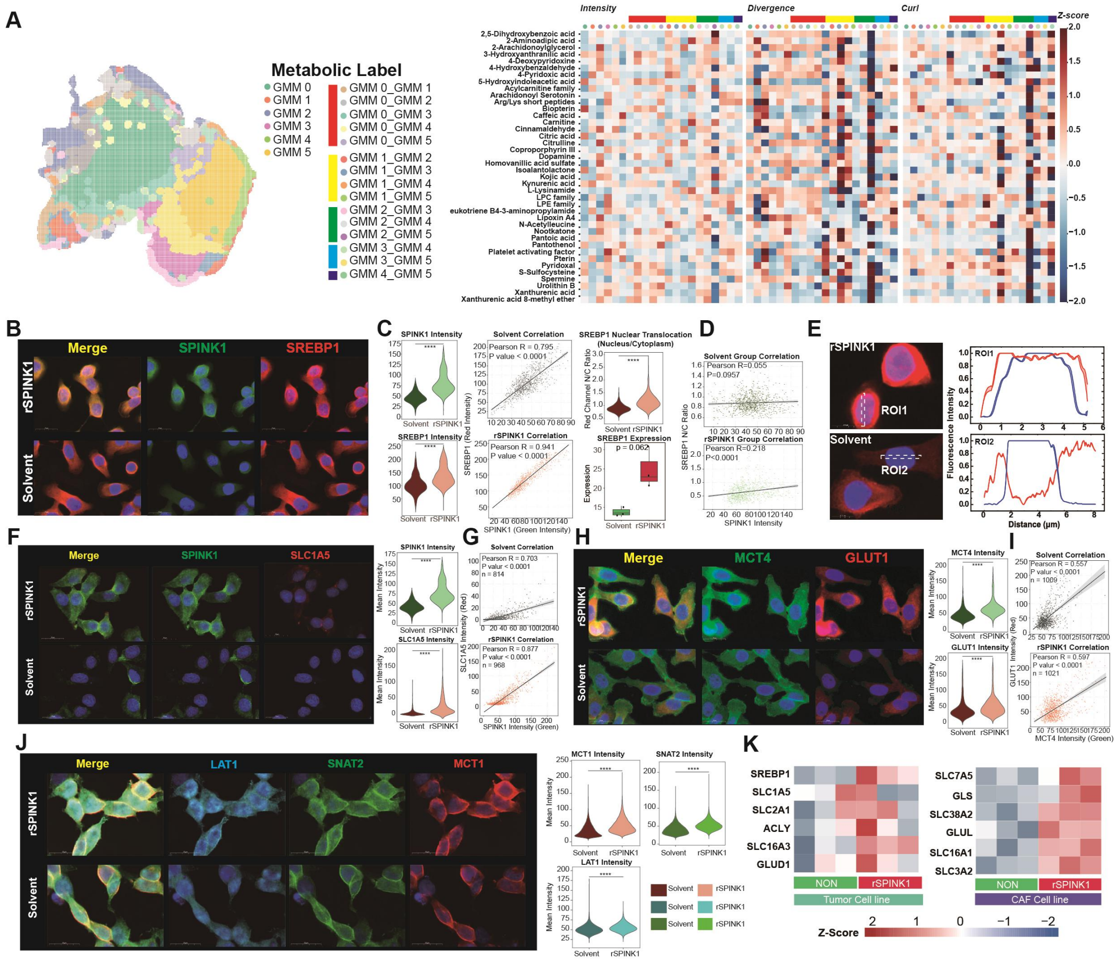


**Figure S12. E2 tumor cells and C1 CAFs exhibit distinct metabolic programs associated with SPINK1-induced SREBP1 nuclear translocation and transporter remodeling.**

(A) Spatial organization and metabolic characterization of GMM-defined regions and their interfaces. Left, spatial map of six core metabolic regions (GMM0–GMM5) and 15 pairwise boundary-associated transition zones, labeled GMMi_GMMj. Right, heatmaps showing Z-score-normalized ion intensity, divergence, and curl for selected metabolite-associated features across core regions and transition zones. Divergence and curl were calculated from intensity-derived spatial vector fields. Rows represent features, and columns represent region categories; colored annotations correspond to the spatial map.

(B) Representative confocal immunofluorescence images of T24 bladder cancer cells treated with recombinant SPINK1 (rSPINK1) or solvent control. SPINK1 is shown in green, SREBP1 in red,

and nuclei stained with DAPI in blue.
(C) Image-based single-cell quantification following cell segmentation. Violin plots show SPINK1 and SREBP1 fluorescence intensities in solvent- and rSPINK1-treated T24 cells. Scatterplots show the relationship between SPINK1 and SREBP1 intensities within each condition, with linear regression lines and Pearson correlation statistics. SREBP1 nuclear translocation was quantified as the nuclear-to-cytoplasmic (N/C) fluorescence ratio. The box plot shows SREBP1 transcript abundance determined by RNA-seq; statistical significance was evaluated using a two-sided t test ($P = 0.062$).
(D) Pearson correlation analysis between single-cell SPINK1 fluorescence intensity and the SREBP1 N/C ratio in solvent- and rSPINK1-treated T24 cells. Regression lines, Pearson correlation coefficients, and P values are indicated.
(E) Representative high-magnification images showing the subcellular distribution of SREBP1 in rSPINK1- and solvent-treated T24 cells. Dashed lines indicate ROI1 and ROI2 used for line-scan analysis. Corresponding profiles show normalized SREBP1 (red) and DAPI (blue) fluorescence intensities as a function of distance, illustrating the redistribution of SREBP1 between cytoplasmic and nuclear compartments.
(F) Representative confocal images of SPINK1 (green) and the glutamine transporter SLC1A5 (red) in rSPINK1- and solvent-treated T24 cells. Nuclei are shown in blue.
(G) Single-cell quantification of SPINK1 and SLC1A5 fluorescence intensities. Scatterplots show their relationship in solvent- and rSPINK1-treated cells, with Pearson correlation coefficients, P values, and analyzed cell numbers indicated.
(H) Representative confocal images of MCT4 (green) and GLUT (red) in rSPINK1- and solvent-treated T24 cells. Nuclei are counterstained with DAPI (blue).
(I) Violin plots showing single-cell MCT4 and GLUT fluorescence intensities following rSPINK1 or solvent treatment. Scatterplots show the relationship between MCT4 and GLUT intensities within each condition, with Pearson correlation statistics and cell numbers indicated.
(J) Representative confocal images of patient-derived CAFs treated with rSPINK1 or solvent control. LAT1, SNAT2, and MCT1 are shown in cyan, green, and red, respectively, with DAPI in blue. Violin plots quantify single-cell fluorescence intensities of MCT1, SNAT2, and LAT1.
(K) Heatmap of RNA-seq-derived expression of selected metabolic genes in T24 tumor cells following rSPINK1 or solvent treatment. Genes include SREBP1, SLC1A5, SLC2A1, ACLY, SLC16A3, and GLUD1. Values are Z-score normalized by gene. Heatmap of RNA-seq-derived expression of selected metabolic genes in patient-derived CAFs following rSPINK1 or solvent treatment. Genes include SLC7A5, GLS, SLC38A2, GLUL, SLC16A1, and SLC3A2. Values are Z-score normalized by gene.

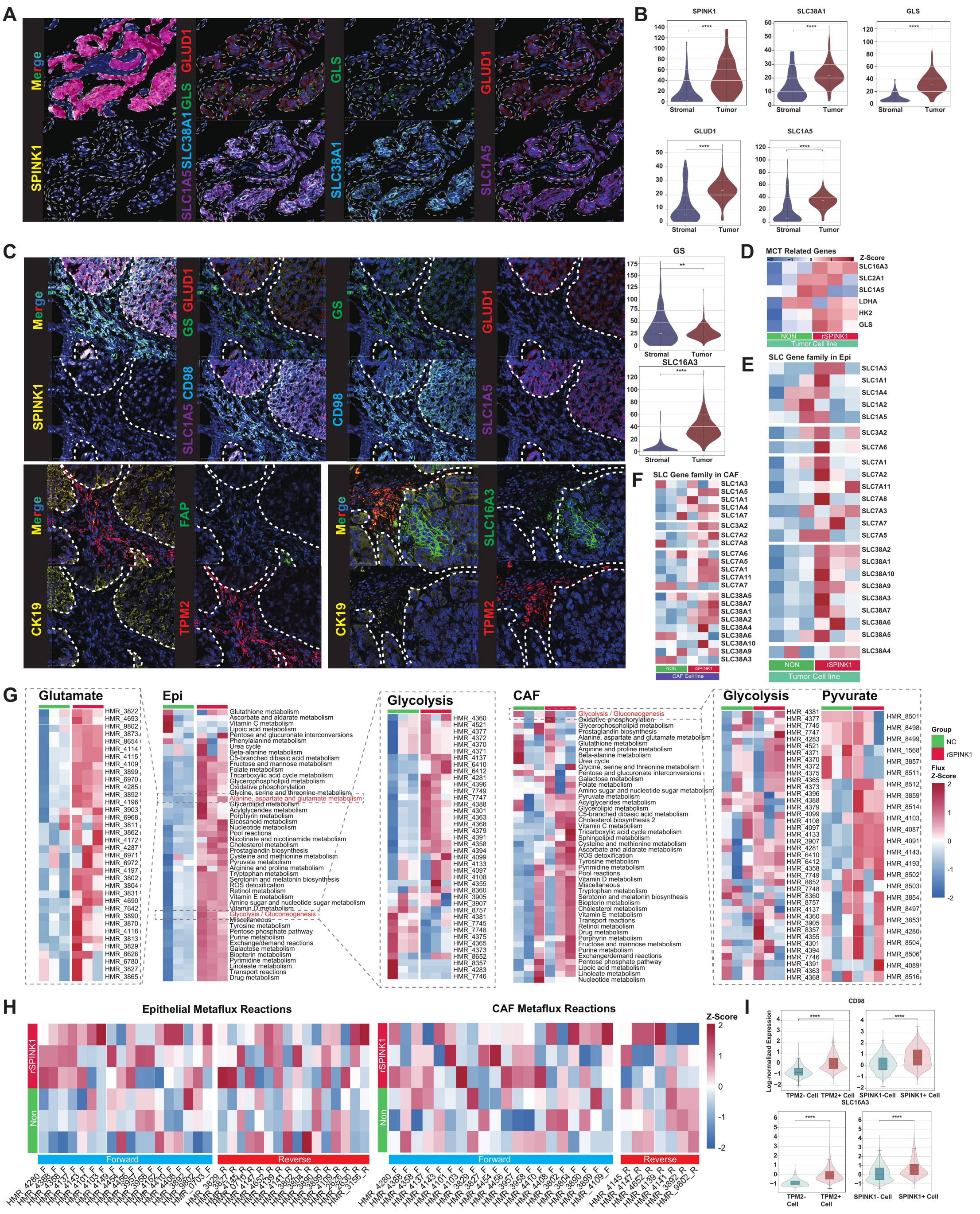


**Figure S13. Spatial and transcriptomic analyses define SPINK1-associated metabolic transporter programs in bladder cancer.**

(A) Representative multiplex immunofluorescence image of a SPINK1+ epithelial–tumor region. SPINK1, SLC38A1, GLS, GLUD1, and SLC1A5 are shown with DAPI nuclear counterstain. The red-shaded area delineates the epithelial–tumor compartment used for quantification.

(B) Single-cell fluorescence intensity distributions of SPINK1, SLC38A1, GLS, GLUD1, and SLC1A5 in stromal and epithelial–tumor compartments corresponding to (A). Statistical significance was evaluated using two-sided Wilcoxon rank-sum tests.

(C) Multiplex immunofluorescence of serial bladder tumor sections stained with complementary panels. SPINK1 and metabolic proteins (GS, GLUD1, SLC1A5, CD98, SLC16A3) are shown together with CK19+ epithelial cells and FAP+/TPM2+ stromal cells. White dashed lines indicate epithelial–tumor boundaries. Violin plots compare GS and SLC16A3 intensities between stromal and epithelial–tumor compartments (two-sided Wilcoxon rank-sum tests).

(D) RNA-seq heatmap of metabolism-related genes in T24 cells treated with recombinant SPINK1 (rSPINK1) or control (Solvent). Genes include *SLC16A3*, *SLC2A1*, *SLC1A5*, *LDHA*, *HK2*, and *GLS*. Values are gene-wise Z-scores.

(E) Heatmap of selected SLC transporter genes in rSPINK1- and control-treated T24 cells. Rows represent *SLC1*, *SLC3*, *SLC7*, and *SLC38* family members; columns represent RNA-seq samples. Values are gene-wise Z-scores.

(F) Heatmap of SLC transporter genes in patient-derived CAFs following rSPINK1 or control treatment. Values are gene-wise Z-scores.

(G) HMR-based transcriptome-inferred metabolic pathway scores in T24 cells and patient-derived CAFs treated with rSPINK1 or solvent.

(H) MetaFlux-inferred metabolic pathway activities in T24 cells and patient-derived CAFs treated with rSPINK1 or solvent.

(I) Single-cell protein abundance in bladder cancer TMA samples. Violin plots show log-normalized CD98 and SLC16A3 levels in TPM2− vs TPM2+ cells and SPINK1− vs SPINK1+ cells. Statistical significance was assessed using two-sided Wilcoxon rank-sum tests.

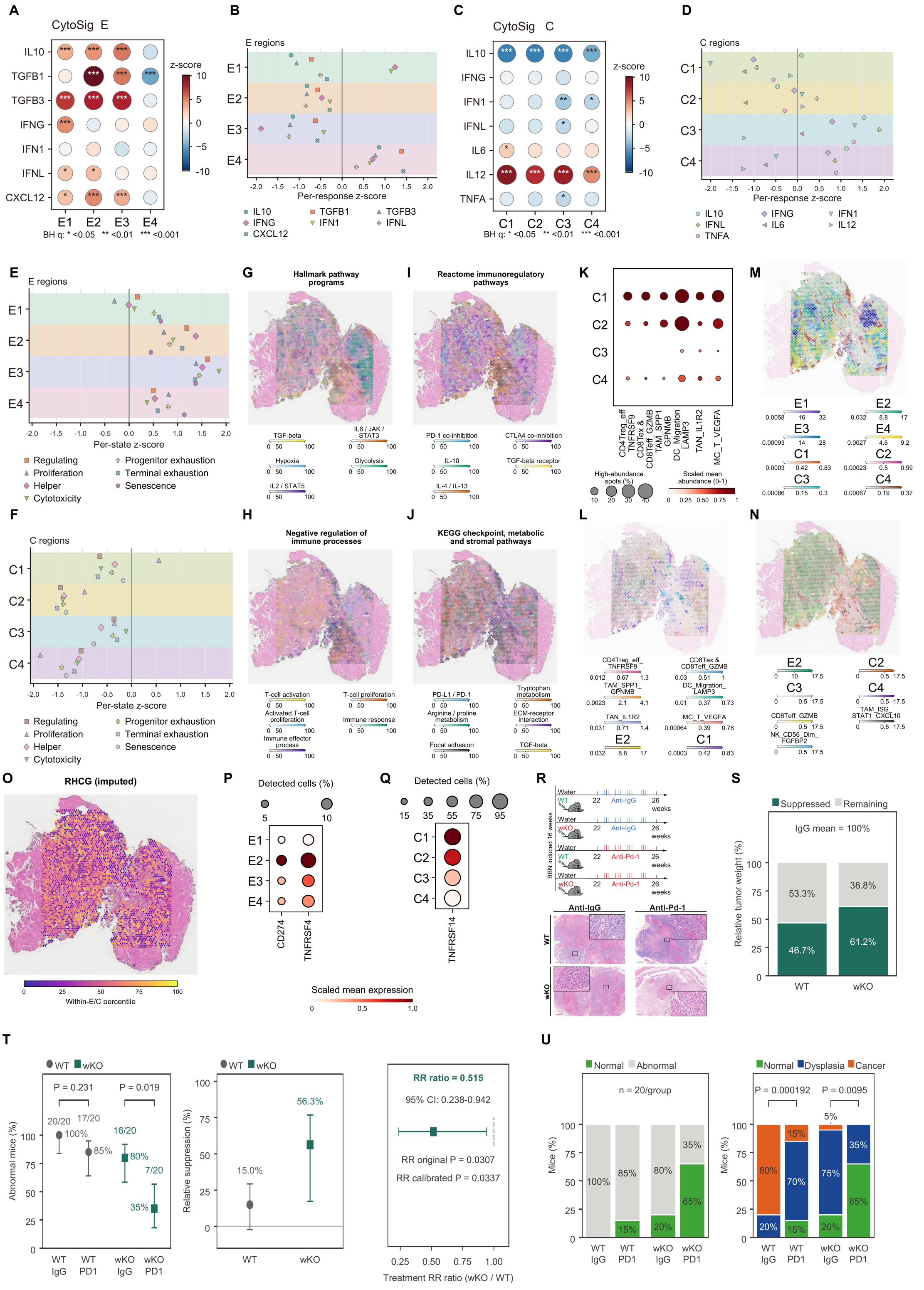

**Figure S14. Spatial immune programs and anti–PD-1 treatment responses in bladder cancer.**

(A) Cytokine response scores inferred using CytoSig in epithelial regions E1 to E4.

(B) Standardized cytokine response profiles in epithelial regions E1 to E4.

(C) Cytokine response scores inferred using CytoSig in CAF regions C1 to C4.

(D) Standardized cytokine response profiles in CAF regions C1 to C4.

(E) TCellSI functional profiles in epithelial regions E1 to E4, with independent standardization of

each program.
(F) Corresponding TCellSI functional profiles in CAF regions C1 to C4.
(G) Spatial distributions of Hallmark signaling and metabolic program scores.
(H) Spatial distributions of gene set scores associated with negative regulation of immune processes.
(I) Spatial distributions of Reactome immunoregulatory pathway scores.
(J) Spatial distributions of KEGG checkpoint, metabolic, and stromal pathway scores.
(K) Inferred immune cell abundance across CAF regions C1 to C4. Dot size represents the proportion of spots exceeding the upper quartile threshold for each population; color indicates scaled mean abundance.
(L) Inferred spatial abundances of E2 epithelial cells, C1 CAFs, and the indicated immune populations.
(M) Spatial abundances of epithelial states E1 to E4 and CAF states C1 to C4 inferred using cell2location and overlaid on H&E images.
(N) Inferred spatial abundances of E2 epithelial cells, CAF states C2 to C4, and selected populations of CD8 T cells, macrophages, and NK cells.
(O) Spatial distribution of imputed RHCG expression overlaid on H&E, expressed as percentile ranks among spots assigned to epithelial or CAF compartments.
(P) Expression of CD274 and TNFRSF4 across epithelial states E1 to E4.
(Q) Expression of TNFRSF14 across CAF states C1 to C4.
(R) Experimental design and representative H&E images of bladder sections from BBN exposed *wild type (WT)* and *Spink1-wKO* mice receiving IgG or anti PD1 treatment.
(S) Relative suppression of tumor weight following PD1 blockade, normalized to the corresponding IgG control within each genotype (n = 6 mice per group).
(T) Prevalence of abnormal histopathology, relative suppression, and the ratio of treatment risk ratios between *Spink1-wKO* and *WT* mice. Error bars represent 95% confidence intervals. Prevalence comparisons used two-sided Fisher exact tests with Holm correction.
(U) Histopathological distributions across treatment groups, classified as normal versus abnormal and as normal, dysplasia, or cancer (n = 20 mice per group). Comparisons across the three categories used two-sided Fisher Freeman Halton exact tests with Holm correction.
In (A) and (C), dot color represents Z scores and dot size represents their absolute magnitudes; *$q < 0.05$, **$q < 0.01$, and ***$q < 0.001$ following correction using the Benjamini and Hochberg procedure. Scores in panels G through J were independently normalized from 0 to 100. In (P) and (Q), dot size represents the percentage of cells with detected expression, and color indicates scaled mean expression. Tumor weight and histopathology analyses were performed in separate cohorts.

**Supplementary Table S1. Summary of clinicopathological information of the included patients.**

| | Patient 1 | Patient 2 | Patient 3 |
|---|---|---|---|
| Tissue Type | Cancer/Adjacent Tissue | Cancer/Adjacent Tissue | Cancer/Adjacent Tissue |
| Surgery Date | October 13, 2023 | October 15, 2023 | October 18, 2023 |
| Surgical Organ | Bladder | Bladder | Bladder |
| Gender | Male | Male | Female |
| Age | 71 | 70 | 67 |
| Pathological Classification | Urothelial Carcinoma of the Bladder | Urothelial Carcinoma of the Bladder | Urothelial Carcinoma of the Bladder |
| T Stage | T4 | T3 | T3 |
| N Stage | N1 | N0 | N0 |
| M Stage | M0 | M0 | M0 |

**Supplementary Table S2. AJCC Seventh Edition anatomic stage grouping for urinary BLCA [1].**

| AJCC-7 stage | TNM combination | Pathological interpretation |
|---|---|---|
| Stage 0a | Ta, N0, M0 | Non-invasive papillary urothelial carcinoma confined to the urothelium, without stromal invasion |
| Stage 0is | Tis, N0, M0 | Flat urothelial carcinoma in situ without stromal invasion |
| Stage I | T1, N0, M0 | Invasion of the subepithelial connective tissue (lamina propria), without muscularis propria involvement |
| Stage II | T2a or T2b, N0, M0 | Invasion of the muscularis propria; pT2a involves the inner half and pT2b the outer half |
| Stage III | T3a, T3b, or T4a; N0, M0 | Microscopic or macroscopic perivesical extension, or direct invasion corresponding to T4a |
| Stage IV | T4b, N0, M0; any T, N1–N3, M0; or any T, any N, M1 | Pelvic-wall or abdominal-wall invasion, regional nodal metastasis, or distant metastatic disease |

Note. The table paraphrases the AJCC-7 urinary bladder chapter. Histological grade is not part of the anatomic stage group and must be recorded separately.

**Supplementary Table S3. AJCC Seventh Edition TNM definitions for urinary BLCA [1].**

| TNM category | Standard pathological definition |
|---|---|
| TX | Primary tumour cannot be assessed |
| T0 | No evidence of a primary tumour |
| Ta | Non-invasive papillary carcinoma |
| Tis | Carcinoma in situ (flat tumour) |
| T1 | Tumour invades the subepithelial connective tissue (lamina propria) |
| T2 | Tumour invades the muscularis propria |
| pT2a | Tumour invades the superficial muscularis propria (inner half) |
| pT2b | Tumour invades the deep muscularis propria (outer half) |
| T3 | Tumour invades the perivesical tissue |
| pT3a | Perivesical invasion is microscopic |
| pT3b | Perivesical invasion is macroscopic and presents as an extravesical mass |
| T4 | Tumour directly invades adjacent structures, as defined by T4a or T4b |
| T4a | Tumour invades the prostatic stroma, uterus, or vagina |
| T4b | Tumour invades the pelvic wall or abdominal wall |
| NX | Regional lymph nodes cannot be assessed |
| N0 | No regional lymph-node metastasis |
| N1 | Metastasis in a single regional lymph node in the true pelvis |
| N2 | Metastases in multiple regional lymph nodes in the true pelvis |
| N3 | Metastasis in one or more common iliac lymph nodes |
| M0 | No distant metastasis |
| M1 | Distant metastasis |

Note. True-pelvic regional nodes include hypogastric, obturator, external iliac, and presacral nodes. AJCC-7 classifies common iliac nodal involvement as N3. The c, p, y, r, and a prefixes should be retained when applicable.

**Supplementary Table S4. Histological grade recorded independently of AJCC stage [1-3].**

| Grade code | Histological classification | Reporting definition |
|---|---|---|
| LG | Low-grade urothelial carcinoma | Urothelial carcinoma classified as low grade under the applicable WHO/ISUP criteria |

| Grade code | Histological classification | Reporting definition |
|---|---|---|
| HG | High-grade urothelial carcinoma | Urothelial carcinoma classified as high grade under WHO/ISUP criteria; carcinoma in situ is high grade |
| GX | Histological grade cannot be assessed | The specimen or pathological information is insufficient for reliable grade assignment |

Note. Grade and TNM stage describe different pathological characteristics. Neither variable should be inferred from the other. AJCC-7 is retained here for historical study consistency; contemporary WHO terminology may be cross-referenced without silently restaging the cohort.

**Supplementary Table S5. Study-defined mouse bladder histopathological progression scale [4-8].**

| **Score** | **Diagnostic category** | **Operational histopathological criteria** | **Carcinoma endpoint** |
|---|---|---|---|
| 0 | No proliferative urothelial lesion | Urothelial thickness, polarity, and cytology within the expected background range | No |
| 1 | Urothelial hyperplasia without atypia | Simple, papillary, or nodular thickening with preserved organization and no definite atypia | No |
| 2 | Atypical hyperplasia or dysplasia-like lesion | Nuclear enlargement or hyperchromasia, polarity loss, irregular arrangement, or increased mitoses, without basement-membrane penetration | No; preneoplastic |
| 3 | Papilloma or low-atypia non-invasive papillary neoplasm | Exophytic or inverted papillary growth without destructive stromal invasion; retain the exact diagnosis | No, unless separately diagnosed as carcinoma |
| 4 | Carcinoma in situ or unequivocal non-invasive urothelial carcinoma | Severe cytological or architectural atypia with an intact basement membrane; record flat and papillary patterns separately | Yes |
| 5 | Lamina propria-invasive urothelial carcinoma | Definite basement-membrane penetration into subepithelial connective tissue, without muscularis propria involvement | Yes |
| 6 | Muscularis propria-invasive urothelial carcinoma | Unequivocal invasion of the muscularis propria or equivalent smooth-muscle layer | Yes |

| Score | Diagnostic category | Operational histopathological criteria | Carcinoma endpoint |
|---|---|---|---|
| 7 | Locally advanced urothelial carcinoma | Extension through the bladder wall into perivesical tissue or an adjacent organ | Yes |

*Note.* This index is a study-defined analytical scale, not an official AJCC-to-mouse conversion and not an INHAND numerical severity grade. The exact morphological diagnosis takes precedence over the score. Carcinoma incidence must be based only on unequivocal carcinoma diagnoses.
